# How synonymous mutations alter enzyme structure and function over long time scales

**DOI:** 10.1101/2021.08.18.456802

**Authors:** Yang Jiang, Syam Sundar Neti, Ian Sitarik, Priya Pradhan, Philip To, Yingzi Xia, Stephen D. Fried, Squire J. Booker, Edward P. O’Brien

## Abstract

The specific activity of enzymes can be altered over long time scales in cells by synonymous mutations, which change an mRNA molecule’s sequence but not the encoded protein’s primary structure. How this happens at the molecular level is unknown. Here, we investigate this issue by applying multiscale modeling to three *E. coli* enzymes - type III chloramphenicol acetyltransferase, D-alanine–D-alanine ligase B, and dihydrofolate reductase. This modeling involves coarse-grained simulations of protein synthesis and post-translational behavior, all-atom simulations as a test of robustness, and QM/MM calculations to characterize enzymatic function. We first demonstrate that our model predicts experimentally measured changes in specific activity due to synonymous mutations. Then, we show that changes in codon translation rates induced by synonymous mutations cause shifts in co-translational and post-translational folding pathways that kinetically partition molecules into subpopulations that very slowly interconvert to the native, functional state. These long-lived states exhibit reduced catalytic activity, as demonstrated by their increased activation energies for the reactions they carry out. Structurally, these states resemble the native state, with localized misfolding near the active sites of the enzymes. The localized misfolding involves noncovalent lasso entanglements - a topology in which the protein backbone forms a loop closed by noncovalent native contacts which is then threaded by another portion of the protein. Such entanglements are often kinetic traps, as they can require a large proportion of the protein to unfold, which is energetically unfavorable, before they can disentangle and attain the native state. The near-native structures of these misfolded states allow them to bypass the proteostasis machinery and remain soluble, as they exhibit similar hydrophobic surface areas as the native state. These entangled structures persist in all-atom simulations as well, indicating that these conclusions are independent of model resolution. Moreover, the structures of long-lived entangled states are supported by agreement with limited-proteolysis mass spectrometry results. Thus, synonymous mutations cause shifts in the co- and post-translational structural ensemble of proteins, whose altered subpopulations lead to long-term changes in the specific activities of some enzymes. The formation of entangled subpopulations is a plausible mechanism through which changes in translation elongation rate alter ensemble-averaged specific activities, which can ultimately affect the efficiency of biochemical pathways and phenotypic traits.

## Introduction

In both *in vitro* and in vivo experiments, the specific activity of protein enzymes – that is, the catalytic turnover per unit time per unit mass of soluble protein – can change depending on the codons used to encode the protein^1, 2, 3, 4, 5, 6^. The specific activity of the *E. coli* enzyme type III chloramphenicol acetyltransferase (CAT-III), for example, decreases by approximately 20% for more than 20 minutes when fast-translating synonymous mutations are introduced into its transcript^1, 6^. This change in activity is considered long-lived as it is longer than *E. coli* cell doubling time (~20 minutes), and can have an impact on daughter cells. Synonymous mutations change the sequence of nucleotides composing an mRNA molecule, which in turn changes the speed at which translation elongation occurs^7^ but not the protein primary structure. Specific activity measurements control for differences in protein expression and the formation of insoluble aggregates through centrifugation or gel separation^1, 2, 3, 8^. For enzymes that do not require post-translational modifications, these observed changes in specific activity indicate that inside cells, newly expressed proteins can populate long-lived conformational states that are not native, have reduced functionality, somehow bypass the chaperone and degradation machinery, and do not aggregate. Furthermore, these observations indicate that the distribution of these kinetically trapped states is sensitive to changes in translation elongation speed.

This alternative state of soluble proteins is distinct from the three typical states^9^ of a protein being either folded and functional, misfolded and aggregated, or degraded. The structural, kinetic, and energetic properties of this alternative state of proteins is a fundamental, unanswered question in biochemistry and molecular biology. Soluble, nonfunctional states do not just occur for enzymes, other protein functions can also be altered. The hormone-transporting protein transthyretin, for example, can have 20% of its soluble fraction in a nonfunctional state^10^. There are hints in the literature of some of the structural properties of this state. NMR-derived structures of β-gamma-crystallin produced from two synonymous mRNA variants found that it populated two soluble conformations differing in the formation of a disulfide bond^11^. Disulfide bonds represent an energetically strong constraint on folding topologies and are uncommon, only 5% and 20%, respectively, of *E. coli* and human cytoplasmic proteins contain disulfide bonds^12^. Furthermore, many enzymes that exhibit specific activity changes due to synonymous mutations do not contain disulfide bonds^1, 2, 3, 5^. Thus, the nature of these structurally altered subpopulations has not yet been resolved, although experimental data rule out misfolding involving quaternary structures. For example, gel separation and chromatography experiments rule out the formation of off-pathway dimers and higher-order oligomers for many enzymes that normally function in a monomeric state^1, 2, 4, 8^.

Here, we use a novel multiscale approach across the dimensions of time, space, and energy to simulate the synthesis, post-translational behavior, and function of enzymes under different translation rate schedules that arise from synonymous codons. We establish that our modeling approach can predict the experimentally measured changes in specific activity for CAT-III, D-alanine–D-alanine ligase B (DDLB), and dihydrofolate reductase (DHFR). We then apply the model to enzymes whose activity is sensitive to synonymous mutations and enzymes whose activity is not. Dissecting the structures and kinetics of co- and post-translational folding that occur in our simulations, we show that synonymous mutations shift the folding pathways and populations of near-native non-covalent lasso entanglements, each of which has its own intrinsic activity (*k*cat) and leads to long-term changes in the enzymatic specific activity.

## Methods

### Coarse-grained simulation model

To model transitions between the native state and the unfolded state we utilized a Gō-based coarse-grained model^13, 14, 15, 16, 17^ for all the proteins studied. Briefly, this coarse-grained model represents each residue as a single interaction site centered on the Cα position and uses a structure-based potential energy function (see Supplementary equation (1)). The force field parameters were optimized by training the parameters to reproduce the folding stability free energies of 18 small single-domain proteins^18^, followed by assignment of the minimum values that can reproduce the structural stability for a given protein. To map the simulation timescale to an experimental timescale, we use the scaling factor *α*, which is the ratio of bulk experimental folding times to simulated folding times (see Supplementary Table 4). The high-resolution crystal structure PDB 4v9d^19^ was used to coarse-grain the *E. coli* ribosome, with the A- and P-site tRNA molecules modeled based on the PDB structure 5jte.^20^ The entire 50S subunit, including the A- and P-site tRNAs, was coarse-grained using the three/four-point model of ribosomal RNA^13^ and the Cα model for ribosomal protein^13^ and then truncated to include only those coarse-grained interaction sites within 30 Å of the centerline of the exit tunnel, within 20 Å of the peptidyl transferase center (PTC, identified as A2602 in the 23S rRNA of the *E. coli* ribosome), and the interaction sites at the ribosome surface near the exit tunnel opening. This resulted in 4,577 interaction sites for the cropped coarse-grained *E. coli* ribosome.

Protein synthesis on the ribosome was modeled using a continuous synthesis protocol that describes A-site tRNA binding, peptidyl transfer, tRNA translocation, and ribosome trafficking^21^ at each nascent chain length using codon translation rates obtained from the model of Fluitt & Viljoen^22^ (see Supplementary equations (11) and (13)). Post-translational dynamics were modeled by simulating the nascent protein in the absence of the ribosome. Simulations for both co- and post-translational folding were performed via Langevin dynamics with a collision frequency of 0.05 ps^-1^ and a time step of 15 fs using OpenMM^23^. Details of the model setup can be found in Supplementary Methods Sections 1 to 8.

### Virtual screening for enzymes that exhibit kinetic traps

To identify candidate enzymes that may have kinetic traps, we created a dataset of well-characterized monomeric *E. coli* enzymes by searching the relevant databases EzCatDB^24, 25^, UniProt^26^ and RCSB PDB^27^. The wild-type enzymes were then parameterized, and their synthesis and post-translational dynamics were simulated with a 14-day wall time. The candidates were selected by using the scoring function

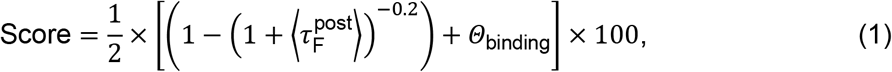

Where 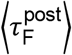 is the mean post-translational folding time calculated by using the double-pathway kinetics scheme of Supplementary equation (6) with no delay time (*i.e.*, *t_1_* = *t*_2_ = 0) and Θ_binding_ is an indicator of whether (equals 1) or not (equals 0) misfolding occurs at or near the substrate binding site and persists to the end of the simulation. We assigned equal weights for these two terms because they are considered equally important in identifying long-lived kinetic traps that have perturbed enzymatic functions. A higher score indicates a higher possibility for the enzyme to have long-lived kinetic traps that have misfolding in the substrate binding pocket during its post-translational folding dynamics. Further details of the screening can be found in Supplementary Methods Section 9.

### Characterizing post-translational misfolded structures

The misfolded structures of the post-translational folding process were characterized by using the metrics *G* and *Q*_act._ *G* is an order parameter that measures the extent to which there is a change of entanglement in a given structure compared to the native structure and is calculated as

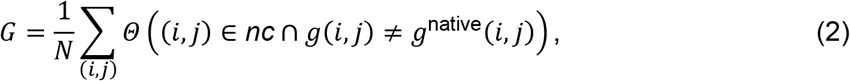

where (*i*, *j*) is one of the native contacts in the native crystal structure; *nc* is the set of native contacts formed in the current structure; *g(i, j)* and *g^native^(i, j)* are, respectively, the total linking number of the native contact (*i*, *j*) in the current and native structures estimated using Supplementary equation (16) (see Supplementary Methods Section 11 for details); *N* is the total number of native contacts within the native structure; and the selection function Θ equals 1 when the condition is true and 0 when it is false. The larger *G* is the larger the number of residues that have changed their entanglement status relative to the native state. That is *G* reports on the presence of non-native entanglements in structures.

*Q*_act_ is the fraction of native contacts that have formed in the enzyme substrate binding pocket. Residues composing the substrate binding pocket were identified as those residues within 8 Å of any atoms of the relevant ligands present in the crystal structure. The *Q*_act_ values were calculated for all the native contacts between one atom within the substrate binding pocket and any other atom, as shown below:

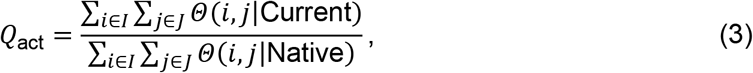

where *i* and *j* are the residue indices and satisfy j > *i* + 3; *I* is the intersection set of residues within secondary structure elements (α-helical or β-strands) and the substrate binding pocket; *J* is the set of all residues within secondary structure elements; Θ(*i, j*|Current) and Θ(*i, j*|Native) are step functions that equal 1 when residue *i* and *j* have native contact and 0 when *i* and *j* do not have native contact in the current structure and native structure, respectively. Native contacts are considered formed when the distance between the Cα atoms of residues *i* and *j* does not exceed 1.2 times their native distance and the native distance does not exceed 8 Å. In the case of CAT-III, residue set *I* also includes native contacts in the trimer interface region (residues 25 to 33 and 150 to 157) to monitor the folding of the trimer interface as well. Note that the native contacts used in estimating *Q*_act_ are restricted to those within secondary structure elements, while the entire set of native contacts is used to estimate *G*. Details of the assessment procedure can be found in Supplementary Methods Section 12.

### Specific activity estimation

For each metastable state *i*, identified using the Markov state modeling procedure reported in Supplementary Methods Section 12, we randomly select five conformations from all the microstates based on the probability distribution of the microstates within the metastable state and back-map them to all-atom structures (see Supplementary Methods Section 13). We then used QM/MM simulations (see Supplementary Methods Section 14) to calculate the transition state barrier for each of the five conformations, and from these, we determined the median activation barrier height 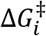. Only the metastable states that form a near-native active site (*Q*_act_ ≥ 0.6), as well as the native state, were taken to estimate ^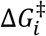^, while the others were considered having an infinite barrier height (zero reaction rate). Assuming the rate constants of each metastable state have similar pre-exponential factors that can be treated as a constant, the specific activity for a protein can be estimated by using the probability distribution of metastable states (state probability p*i*) and the activation free energy barrier height 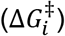 of each state as

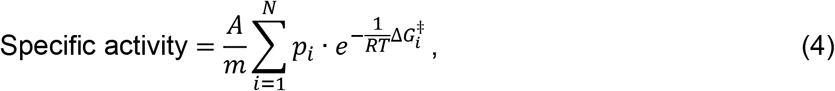

where *m* is the molecular weight of the enzyme; *A* is the pre-exponential factor allowing us to convert 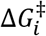 to the reaction rate constant; *N* is the total number of metastable states; *R* is the gas constant; and *T* is the temperature. In many codon usage studies^1, 3, 8^, the relative specific activity is often used to compare the enzymatic activities of a protein produced by synonymous mRNA variants. In this study, the maximum specific activity among the fast and slow synonymous mRNA variants of an enzyme was used for normalization:

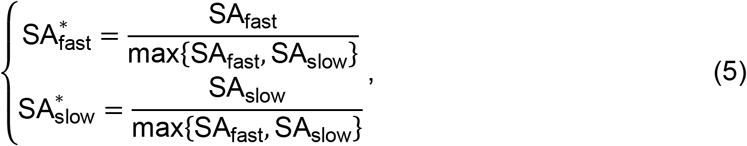

where SA^∗^ is the relative specific activity under saturating conditions. Note that the pre-exponential factor *A* and the molecular weight *m* cancel out when specific activity is normalized; therefore, they do not need to be estimated. The details of estimating the state probability *p_i_* (accounting for the soluble fraction only) and the activation free energy barrier height 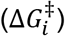 are presented in Supplementary Methods Section 14. The materials and experimental methods for measuring the specific activities of DDLB and DHFR variants are presented in Supplementary Methods Sections 19 and 20, respectively.

## Results

### Recapitulating experimental trends of CAT-III

We first applied our modeling protocol to the *E. coli* enzyme CAT-III to test if the method accurately predicts the influence of synonymous mutations on specific activity. CAT-III has been studied experimentally and shown to have a reduced specific activity when faster-translating codons are introduced through synonymous mutations^1^. We created both fast-and slow-translating synonymous mRNA sequences (denoted, respectively, CAT-III_fast_ and CAT-III_slow_) by replacing each wild-type codon with its fastest or slowest synonymous variant (see Fig. 1b, mRNA sequences are presented in Supplementary Methods Section 22). The resulting slow-mRNA variant takes two times longer to translate than the fast variant (see Fig. 1f). We simulated the synthesis and post-translational behavior of CAT-III resulting from the fast- and slow-mRNA variants and calculated their respective specific activities (equation (4)) at the end of the post-translational simulations. In our model, CAT-III_fast_ exhibited 83.6% (95% confidence interval (CI): [72.0%, 96.5%] from bootstrapping, *p-* value=0.0067 from random permutation test, 10^6^ permutations) of the specific activity of CAT-III_slow_ (Fig. 1g). This comparison, known as the ‘relative specific activity’ (equation (5)), is common in biochemical studies^1, 3^. A 20% decrease in activity has been observed experimentally^1^. Thus, our modeling approach qualitatively recapitulates experimentally observed changes in enzyme activity due to synonymous mutations.

**Figure 1.**
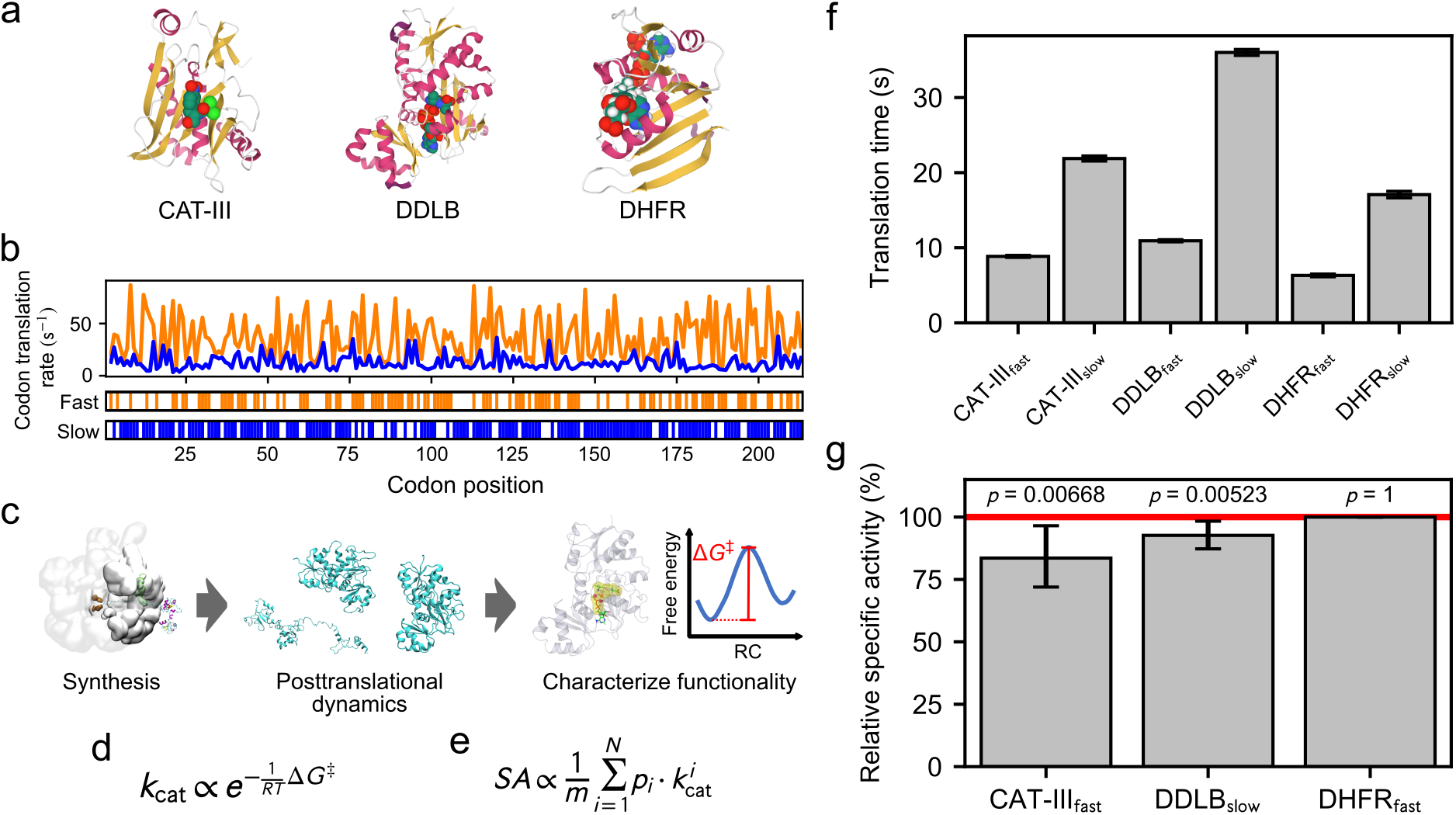
A multiscale approach for understanding the influence of synonymous codons on the structure and function of enzymes. a, Crystal structures of the three enzymes investigated in this study, with secondary structure elements highlighted and substrates presented; b, codon translation rate of CAT-III fast and slow synonymous mRNA variants, with the mutation sites presented on the bottom bars; c scheme diagram of the multiscale approach, including the coarse-grained protein synthesis and post-translational dynamics, and the all-atom characterization of enzymatic functionality; d, enzymatic reaction rate *k*_cat_ estimated from the activation free energy Δ*G*^‡^; e, enzyme specific activity (SA) estimated as the ensemble average of reaction rates *k*cat (see equation (4)); f, comparison of the translation time for the fast and slow synonymous mRNA variants; g, relative SAs for CAT-III, DDLB and DHFR. The SA values are normalized to the higher SA value found in fast- and slow-mRNA variants. Error bars in panels f and g are 95% CIs about the mean, which were estimated using bootstrapping. p-values characterize the difference in SAs between the proteins produced from the fast and slow variants. p-values were calculated using the permutation test. The SAs of CAT-III and DDLB are sensitive to translation speed changes, and that of DHFR is not.

### Other enzymes that are sensitive to changes in translation speed

We next applied our model to identify other *E. coli* enzymes whose specific activities are sensitive to translation speed changes. To do this, we developed a virtual screening strategy that identifies misfolding-prone proteins exhibiting long-lived, post-translational kinetic traps (see Methods), as we hypothesized that the activity of these enzymes is more likely to be sensitive to translation speed changes. Due to the large number of *E. coli* enzymes that have an unknown catalytic mechanism or uncharacterized protein-substrate complex structures, we focused on only 14 well-characterized monomeric enzymes (Table 1). We synthesized each protein on the ribosome using the translation rate arising from their wild-type mRNA sequences, followed by release from the ribosome and a post-translational simulation phase. To identify good candidates, we scored (equation (1)) each of the 14 enzymes based on whether they misfolded near the substrate binding site and whether the misfolding events were long-lived. Porphobilinogen deaminase never exhibited any folding events during the simulations and therefore was excluded from further study. The protein with the highest score, D-alanine–D-alanine ligase B (DDLB), was selected as a candidate whose enzymatic activity we hypothesized is likely to be sensitive to translation speed changes, whereas the protein with the lowest score, dihydrofolate reductase (DHFR), was identified as an enzyme whose activity is likely to be insensitive to synonymous mutations.

**Table 1.** The 14 *E. coli* enzymes studied using our virtual screening strategy. 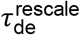, calculated post-translational mean folding time. Θbinding indicates whether misfolding is (‘1’) or is not (‘0’) located at or near the substrate binding site and persists to the end of the simulation. ‘Score’ is the scoring function calculated from equation (1).

| Protein Name | Length (residues) | PDB ID | $\langle \tau_F^{\text{post}} \rangle$ (s) | $\theta_{\text{binding}}$ | Score |
| --- | --- | --- | --- | --- | --- |
| D-alanine—D-alanine ligase B (DDLB) | 306 | 4C5C | $9.8 \times 10^9$ | 1 | 99.5 |
| Phosphoenolpyruvate carboxykinase | 540 | 1AQ2 | $5.2 \times 10^7$ | 1 | 98.6 |
| Quinone oxidoreductase 2 | 303 | 2ZCV | 3.9 | 1 | 63.6 |
| Adenylate kinase | 214 | 1AKE | 9.5 | 0 | 18.7 |
| Dephospho-CoA kinase | 206 | 1VHL | 4.8 | 0 | 14.8 |
| Peptide deformylase | 169 | 3K6L | 3.4 | 0 | 12.8 |
| Protein-L-isoaspartate O-methyltransferase | 208 | 3LBF | 3.4 | 0 | 12.8 |
| Methionyl-tRNA formyltransferase | 315 | 2FMT | 1.0 | 0 | 6.6 |
| Phosphoribosylglycinamide formyltransferase | 212 | 1C2T | 0.65 | 0 | 4.8 |
| Flavodoxin/ferredoxin--NADP reductase | 248 | 1FDR | 0.09 | 0 | 0.9 |
| DNA-3-methyladenine glycosylase 2 | 282 | 3CW7 | 0.07 | 0 | 0.6 |
| 2-Dehydropantoate 2-reductase | 303 | 2OFP | 0.05 | 0 | 0.5 |
| Dihydrofolate reductase (DHFR) | 159 | 4KJK | 0.03 | 0 | 0.3 |
| Porphobilinogen deaminase | 313 | 1PDA | - | - | - |

Next, we simulated the synthesis and post-translational dynamics of DDLB and DHFR from their fast- and slow-translating mRNA variants (sequences are presented in Supplementary Methods Section 22) using the same simulation protocol as applied to CAT-III. The slow variants of DDLB and DHFR translate, respectively, three and two times slower than their fast variants (Fig. 1f). We found that the specific activity of DDLB_slow_ was 92.7% (95% CI: [87.3%, 98.3%]) that of DDLB_fast_ (*p*-value = 0.0052) 60 s after synthesis was completed. DHFR_fast_ exhibited a specific activity 100% (95% CI: [100%, 100%]) that of DHFR_slow_, which were not significantly different (*p*-value = 1). Therefore, our model describes enzymes whose specific activity is either sensitive (DDLB) or insensitive (DHFR) to changes in translation speed over long-time scales.

### Accurate prediction of trends in specific activity

To experimentally test if DDLB_slow_ has a reduced enzymatic activity compared to DDLB_fast_ we recombinantly expressed the fast and slow DDLB variants in *E. coli* using the same mRNA sequences as in the simulations, purified the enzyme, and then assayed the reaction kinetics (see Supplementary Methods Section 19). We observe that the fast variant had a higher level of protein expression 5 hours after induction than the slow variant, consistent with the fast mRNA variant translating faster (Supplementary Fig. 11). The reaction rate constant *k*_cat_ was measured across five biological replicates of the fast and slow variants (see Supplementary Table 11). We find the specific activity of DDLB_slow_ is, on average, 88% 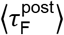, 95% CI: [81.3%, 94.8%]) that of DDLB_fast_ (*p*-value = 0.0186, one-tailed t-test for 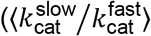. On the other hand, we did the same experiments to measure the *k*cat of fast and slow DHFR variants (see Supplementary Methods Section 20). Consistent with our model prediction, the specific activity of DHFRfast is indistinguishable from that of DHFR_slow_ 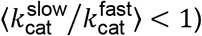 = 91%; 95% CI: [68%, 115%]; *p*-value = 0.5323, two-tailed t-test for 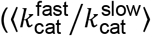; see Supplementary Table 12). Thus, our modeling approach successfully predicts the trends in changes in activity for CAT-III, DDLB and DHFR, indicating it is realistic.

### Near-native, lasso-like entangled structures are populated by the three enzymes

Next, in our simulations we identified the structures, catalytic properties, and folding pathways that cause these activity changes. We first examined the structural distributions of CAT-III, DDLB and DHFR in the post-translational simulations by using a clustering algorithm that uses both structural information and temporal interconversion rates between metastable states. Specifically, after numerous tests of different metrics, we structurally clustered on the basis of the fraction of native contacts formed in substrate binding regions (denoted as *Q*_act_, equation (3)) and the fraction of native contacts that exhibit non-native topological entanglements (denoted as *G*, equation (2), see Methods for description). Microstates were identified by clustering the post-translational trajectories of both synonymous variants together for each protein using the *k*-means algorithm^28^, and then the microstates were coarse-grained to a smaller number of metastable states using the PCCA+ algorithm^29^ (see Fig. 2a and b for CAT-III, Fig. 3a and b for DDLB and Fig. 4a and b for DHFR).

**Figure 2.**
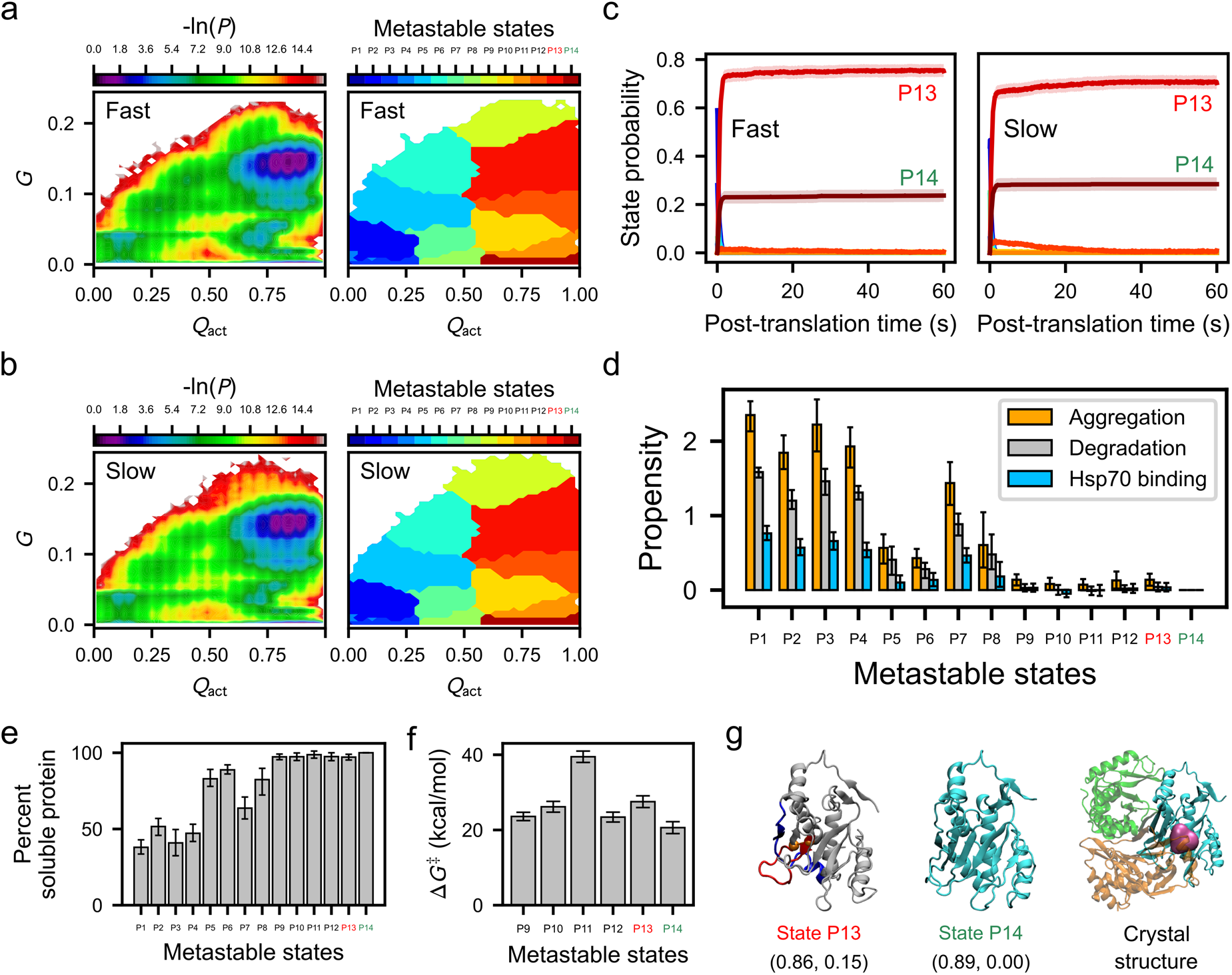
Fast translation partitions more CAT-III into post-translational kinetically trapped entangled states. a and b, Log Probability surfaces (−ln *P*, *P* is the probability of sampling various configurations) for the post-translational folding of fast-translating (a) and slow-translating (b) mRNA variants as a function of the order parameters *Q*_act_ and *G* (left) and the regions corresponding to different metastable states (right). The native state is located in the bottom right-hand corner of these plots. c, Time courses of gross state probabilities (soluble + insoluble; same colors as the metastable states in panels a and b) with error bars shown as transparent stripes for fast (left) and slow (right) variants. d, The aggregation, degradation and Hsp70 binding propensities of each metastable state calculated using Supplementary equation (19). e, The percent soluble protein of each metastable state calculated by Supplementary equation (17). f, The median transition state barrier heights (Δ*G*^‡^) for the native and near-native metastable states calculated from the QM/MM simulations. g, From left to right, the representative structure of near-native kinetically trapped state P13 (the closed loop and threading segment of their entangled regions are colored in red and blue, respectively); the native state P14; and the trimer crystal structure (3cla) with substrate shown in magenta and three monomers shown in green, orange, and cyan. The *Q*_act_ and *G* coordinates of the most probable cluster (microstate) for each state are reported below the structure in the format (*Q*_act_ value, *G* value). For panels a through g, kinetically trapped entangled states are labeled in red; the native state (*i.e.*, state P14) is labeled in green; and others are labeled in black. All error bars represent 95% CIs calculated by bootstrapping. The structure of P13 can be explored interactively at https://obrien-lab.github.io/visualize_entanglements/.

**Figure 3.**
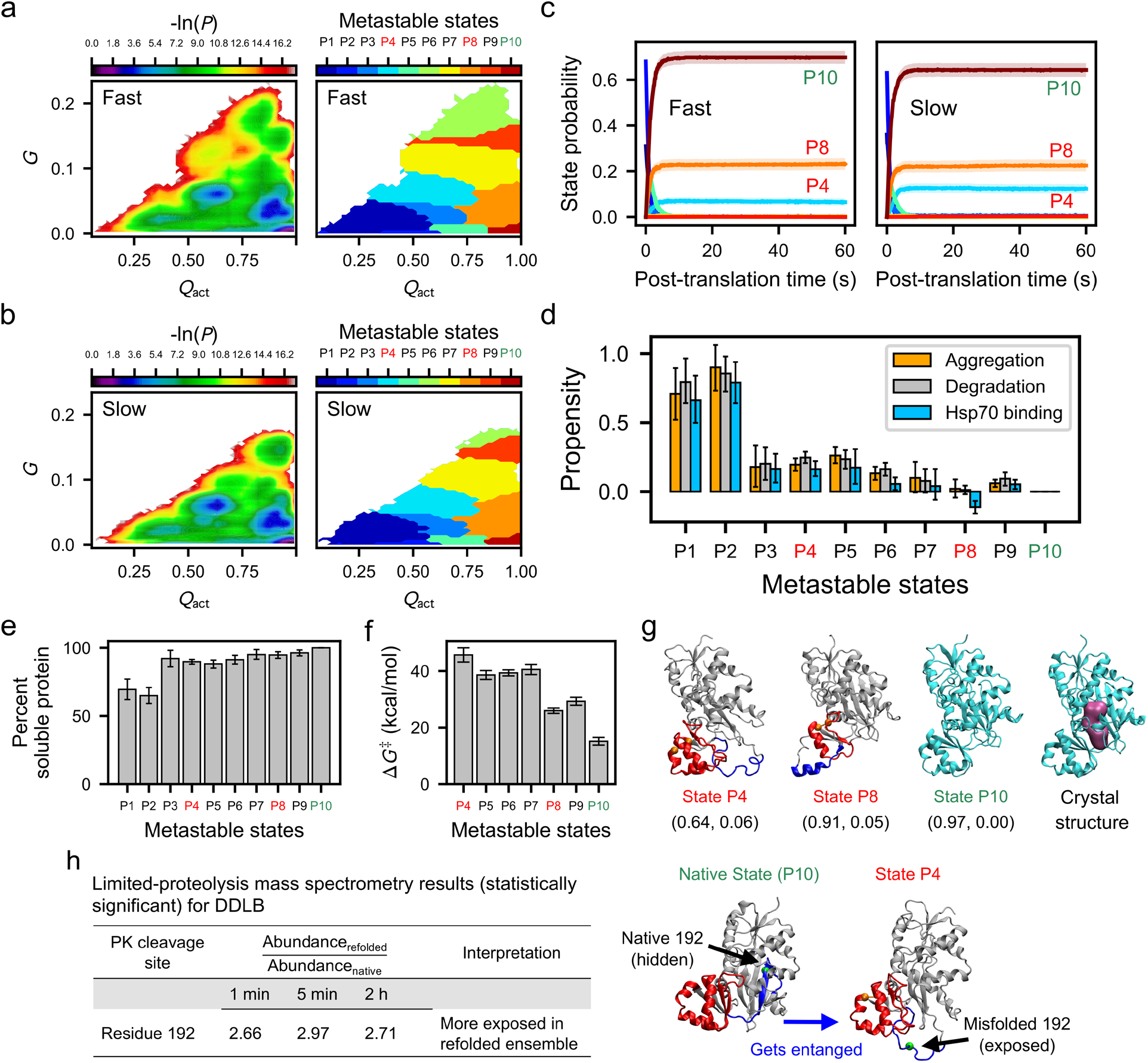
Slow translation partitions more DDLB into post-translational kinetically trapped entangled states. A and b, Log Probability surfaces (-ln*P*, *P* is the probability of sampling various configurations) for the post-translational folding of fast-translating (a) and slow-translating (b) mRNA variants as a function of the order parameters *Q*_act_ and *G* (left) and the regions corresponding to metastable states (right). The native state is located in the bottom right-hand corner of these plots. c, Time courses of gross state probabilities (soluble + insoluble; same colors as the metastable states in panels a and b) with error bars shown as transparent stripes for fast (left) and slow (right) variants. d, The aggregation, degradation and Hsp70 binding propensities of each metastable state calculated using Supplementary equation (19). e, The percent soluble protein of each metastable state calculated by Supplementary equation (17). f, The median transition state barrier heights (Δ*G*^‡^) for the near-native metastable states calculated from the QM/MM simulations. g, From left to right, the representative structure of near-native kinetically trapped states P4 and P8 (the closed loop and threading segment of their entangled regions are colored in red and blue, respectively); the native state P10; and the crystal structure (4c5c) with substrate shown in magenta. The *Q*_act_ and *G* coordinates of the most probable cluster (microstate) for each state are reported below the structure in the format (*Q*_act_ value, *G* value). h, Limited-proteolysis mass spectrometry results for refolded DDLB. Only the peptides showing significantly different abundance (the abundance must have at least a 2-fold difference and the p-value must be less than 0.01) from the refolded protein through all three time points are presented. The representative structure of the native state and the entangled state P4 are presented on the right side. The entanglement in state P4 is represented in the same way of panel g. The native conformation corresponding to the entanglement is highlighted in the native state structure. The PK site residue 192 is shown as a green ball. It is significantly more exposed to solvent in the entangled state P4. For panels a through h, kinetically trapped entangled states are labeled in red; the native state (*i.e.*, state P10) is labeled in green; and others are labeled in black. All error bars represent 95% CIs calculated by bootstrapping. The structures of P4 and P8 can be explored interactively at https://obrien-lab.github.io/visualize_entanglements/.

**Figure 4.**
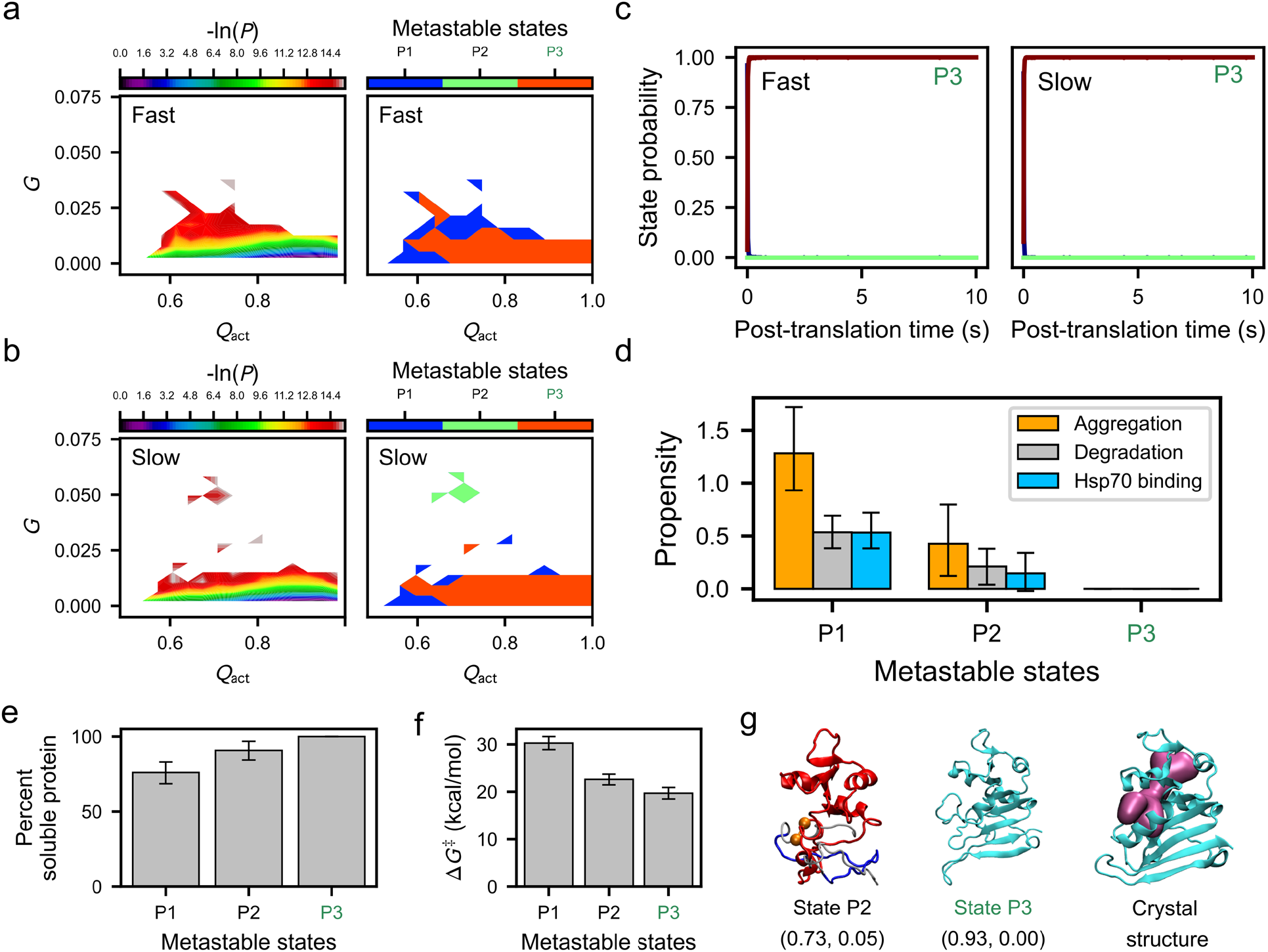
No kinetically trapped states arise in synonymous variants of DHFR. a and b, Log Probability surfaces (−ln(*P)*, *P* is the probability of sampling various configurations) for the post-translational folding of fast (a) and slow (b) mRNA variants as a function of the order parameters *Q*_act_ and *G* (left) and the regions corresponding to metastable states (right). The native state is located in the bottom right-hand corner of these plots. c, Time courses of gross state probabilities (soluble + insoluble; same colors as the metastable states in panels a and b) with error bars shown as transparent stripes for fast (left) and slow (right) variants. d, The aggregation, degradation and Hsp70 binding propensities of each metastable state calculated using Supplementary equation (19). e, The percent soluble protein of each metastable state calculated by Supplementary equation (17). f, The median transition state barrier heights (Δ*G*^‡^) for the near-native metastable states calculated from the QM/MM simulations. g, From left to right, the representative structure of shallow entangled state P2 (the closed loop and threading segment of their entangled regions are colored in red and blue, respectively); the native state P3; and the crystal structure (4kjk) with substrate shown in magenta. The *Q*_act_ and *G* coordinates of the most probable cluster (microstate) for each state are reported below the structure in the format (*Q*_act_ value, *G* value). For panels a through g, kinetically trapped entangled states (if exist) are labeled in red, the native state (*i.e.*, state P3) is labeled in green, and others are labeled in black. All error bars represent 95% CIs calculated by bootstrapping. The structure of P2 can be explored interactively at https://obrien-lab.github.io/visualize_entanglements/.

Across the metastable states, we observed diverse misfolded structures for CAT-III (see Supplementary Fig. 5) and DDLB (see Supplementary Fig. 6). Most of the misfolded structures exhibited topological entanglements, and many of these entangled structures were near native (*Q*_act_ ≥ 0.6 and *G* ≥ 0.02; interactive visualization of some entangled structures is provided on the supplementary website https://obrien-lab.github.io/visualize_entanglements/). Misfolded, metastable states P9, P10, P11, P12 and P13 of CAT-III exhibited *Q*_act_ and *G* values of (0.66, 0.18), (0.72, 0.04), (0.80, 0.02), (0.84, 0.09), and (0.86, 0.15), respectively, while the native state (P14) had values of (0.89, 0.00) (Fig. 2g and Supplementary Fig. 5). The DDLB misfolded states P4, P6, P7, P8, and P9 exhibited values of (0.64, 0.06), (0.83, 0.19), (0.85, 0.09), (0.91, 0.05) and (0.92, 0.15), respectively, while the native state (P10) had values of (0.97, 0.00) (Fig. 3g and Supplementary Fig. 6). In contrast, DHFR exhibited fewer misfolded states and only one entangled, misfolded state, with *Q*_act_ and *G* values of (0.73, 0.05) compared to the values (0.93, 0.00) for its native state (Fig. 4g and Supplementary Fig. 7). All of the topological entanglements that we observe had a noncovalent lasso topology^30, 31, 32, 33, 34^, where native contacts within a certain folded region established a closed loop along the protein backbone and another segment in the same chain threaded through this loop to become entangled (Fig. 5). None of them are topologically knotted, as identified by a knot detection algorithm^35^, meaning that pulling on their termini will result in a fully extended conformation. Thus, all three enzymes sampled near-native, entangled but unknotted structures during folding.

**Figure 5.**
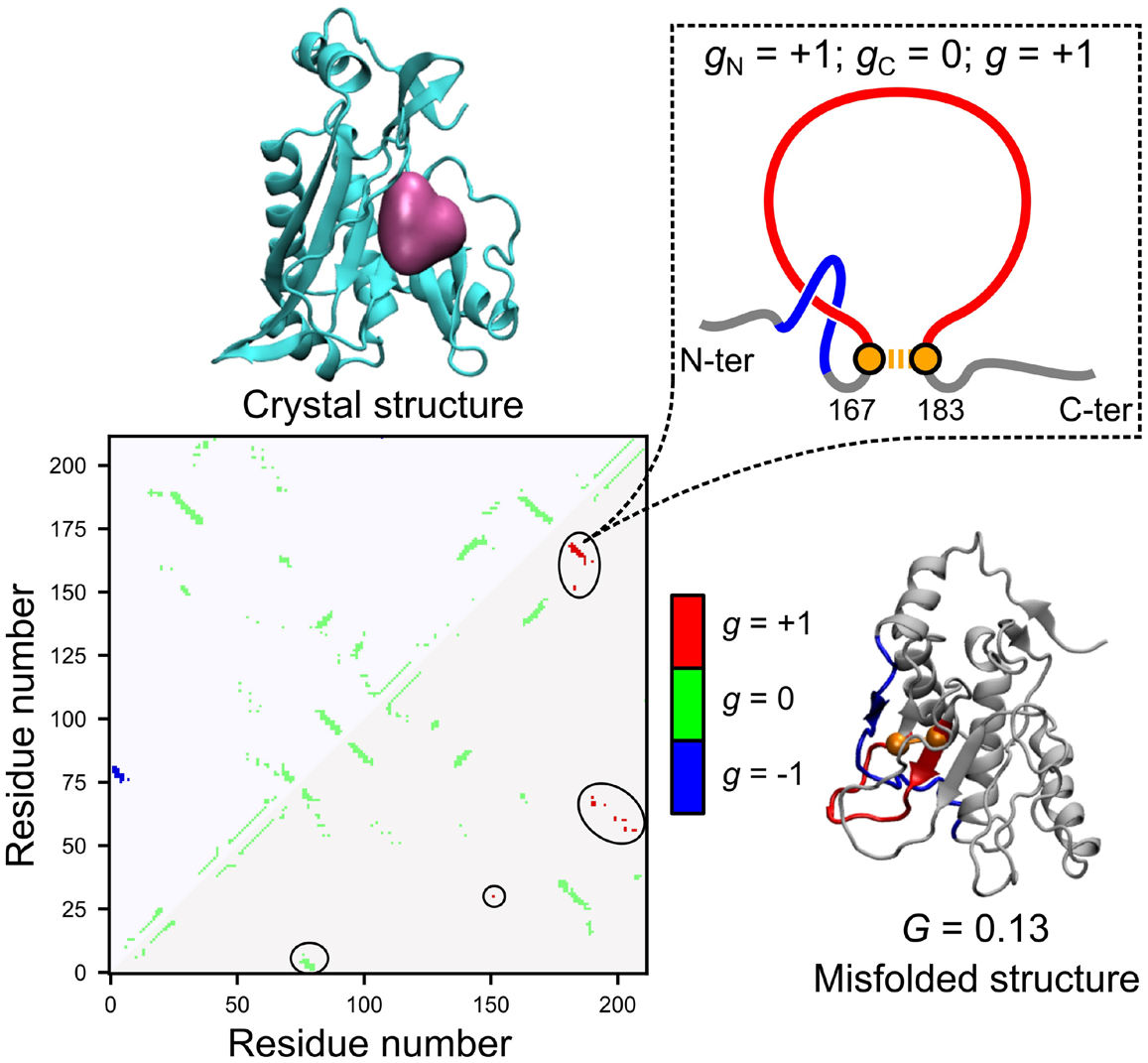
Illustration of the G metric and the noncovalent lasso topology. A contact map of the native contacts is presented on the bottom left, where the upper and lower triangular regions represent the native contacts within the native crystal structure of CAT-III and one of the misfolded structures in state P13, respectively. The native contacts are colored based on the linking number *g* calculated by Supplementary equation (16). A topology diagram of the noncovalent lasso entanglement formed by the native contact of residues 167 and 183, as an instance, is presented at the top right, where the closed loop is shown in red and the threading segment is shown in blue. As described by equation (2), the *G* metric of this misfolded structure, *G* = 0.13, was estimated by counting the number of native contacts within the misfolded structure, whose *g* value is different from that of the same native contact within the crystal structure (*i.e*., the number of native contacts covered by the black circles on the lower triangular region of the contact map), and normalizing them to the total number of native contacts in the crystal structure.

### Some entangled structures are long-lived kinetic traps

To disentangle a misfolded structure, it is often necessary to unfold some portion of the properly folded segments to attain the native state. This can be energetically costly. Therefore, we hypothesized that these near-native, entangled structures are long-lived kinetic traps. To test this hypothesis, we calculated the post-translational probability of being in each metastable state as a function of time and report the results in panels c of Figs. 2, 3 and 4. For CAT-III and DDLB, entangled states (P13) and (P4 and P8), respectively, persisted with appreciable populations (>10%) until 60 s after nascent protein release from the ribosome. For DHFR, the single entangled state (P2), populated only 0.2% of the trajectories, disentangled by 10 s. As a further test, we examined whether any post-translational trajectory of CAT-III or DDLB that reaches an entangled state ever converts to an unentangled state. For CAT-III and DDLB, we find that 92.7% and 100% of trajectories sample an entangled state, and of these, 78.6% and 10.4% do not convert to a state that is not entangled by the end of the simulations. Thus, we conclude that many of these entangled structures represent long-lived kinetic traps that convert to the native state over very longtime scales.

### Entangled structures have altered catalytic properties

The noncovalent lasso entanglement intermediates we observe are a form of misfolding, as they represent ordered structures that are not native. Therefore, we hypothesized that some of these entangled structures have catalytic properties different than those of the native ensemble. To test this hypothesis, we calculated the transition state barrier height of the enzyme reaction of each metastable state that forms native or near-native active site (*Q*_act_ ≥ 0.6) using a back-mapping procedure to an all-atom representation and subsequent QM/MM umbrella sampling simulations of the catalytic reaction (see the potential of mean force plots in Supplementary Fig. 8-10). Thus, for each metastable state, we obtained the activation free energy barrier Δ*G*^‡^ going from reactants to products. For all three proteins, the native state had the lowest activation energy (panel f in Figs. 2, 3 and 4), and the other metastable states had higher barriers. Thus, these misfolded and entangled intermediates contribute to changes in specific activity. Coupled with the observation that entangled structures tend to be long-lived kinetic traps, we also conclude that these specific metastable states lead to reduced specific activity over long time scales.

### Distinctions between shallow and deep entanglements explain why CAT-III and DDLB cannot quickly fold but DHFR can

Not all entangled structures are long-lived kinetic traps; otherwise, the entangled structure of DHFR (state P2 in Fig. 4g and Supplementary Fig. 7) would never disentangle. Sliding a small number of residues out of the closed loop (Fig. 5, also see Supplementary Methods Section 11) tends to be easier than sliding a large number of residues. In addition, unfolding a small number of residues during disentanglement also tends to be easier than unfolding a large number of residues. Therefore, we hypothesized that the minimum number of residues involved in the threading segment, whose reptation can cause disentanglement, and the minimum number of residues needed to unfold during the disentanglement process should correlate with the ability of entangled metastable states to interconvert to unentangled states. To test this hypothesis, we analyzed the representative structures from each metastable state. The entanglement of DHFR involved only, on average, 5 residues in the threading segment, and sliding these segments through the loop should not cause any portion of the protein to unfold. For CAT-III and DDLB, entangled states that never disentangle in the simulations involved, on average, 35 and 8 residues, respectively, in the threading segment and had 31 and 53 residues that need to unfold during disentanglement (see https://obrien-lab.github.io/visualize_entanglements/). These results are consistent with our hypothesis and suggest that because DHFR’s entanglement is ‘shallow’ (*i.e.*, involves a few residues and does not need to unfold to disentangle), thermal fluctuations can easily disentangle this structure, while thermal energy is not sufficient to quickly disentangle the CAT-III and DDLB ‘deep’ entanglements.

### Deep entanglements are usually long-lived regardless of model resolution

To examine how long it will take to disentangle deep and shallow entanglements in a higher resolution model, we back mapped two deep entangled conformations (one from CAT-III state P13 and the other from DDLB state P4) and one shallow entangled conformation (from DHFR state P2) to an all-atom representation (see Supplementary Methods Section 13). We then simulated 30 independent trajectories of each conformation in fully solvated, unrestrained molecular dynamics simulations at 900 K, 800 K, 700 K, 600 K and 550 K (Supplementary Methods Section 15) to monitor the disentangling times. To account for force-field biases affecting kinetics we also simulate and calculate the unfolding time of native DHFR at these temperatures and compared it to its experimentally measured unfolding rate 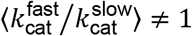, extrapolated to 0M Urea at 298 K)^36, 37^. (Among these three enzymes, only DHFR has an experimentally reported folding kinetics.) The disentangling rates 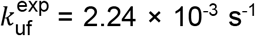 and unfolding rates 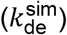 at each temperature were estimated by fitting a single exponential function to the time-dependent survival probability (see panel a in Supplementary Figures 12 and 13 and panels a and b in Supplementary Figure 14). Finally, 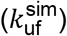 and 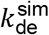 at 298 K were extrapolated using an Arrhenius plot (ln*k* vs. reciprocal temperature). We observe ‘super-Arrhenius’ behavior in the disentangling and unfolding rates, where the Arrhenius plot is better described by a quadratic function than a linear function (Supplementary equation (20); data are presented in panels b and c of Supplementary Figures 12 and 13 and panels c, d and e of Supplementary Figure 14). Such ‘super-Arrhenius’ behavior has been observed in many biological reactions and processes^38, 39^ and arises from the escape rate from local energy minima (which is a quadratic function of reciprocal temperature)^40^. Indeed, experimental measurements of protein unfolding rates over large temperature ranges exhibit super-Arrhenius behavior^41^. We find the all-atom force field accelerates the unfolding of DHFR by 144-fold at 298 K. By correcting *k_de_* for this force-field bias by using this acceleration factor, we estimate the realistic disentangling timescale is 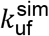. For the deep entanglements in CAT-III and DDLB 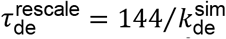 are 6.2 × 10^3^ s (95%CI [29, 1 × 10^6^] s) and 1.3 × 10^4^ s (95%CI [28, 4 × 10^6^] s), respectively. While the shallow entanglement of DHFR gets disentangled with a timescale of 71 s (95%CI [4 × 10^-3^, 2 × 10^6^] s), which is two-orders-of-magnitude faster than the deep entanglements. These results suggest that the deep entangled metastable states we have identified are likely to be long-lived kinetic traps, persisting for hours.

### Native-like entangled states are much less likely than other misfolded states to interact with chaperones, aggregate or be degraded and hence remain soluble

The proteostasis machinery in cells has the potential to catalyze the folding or degradation of these entangled structures. The entangled structures could also potentially aggregate and be removed from the pool of soluble enzymes. We accounted for the effects of these processes by estimating the aggregation, degradation and Hsp70 binding propensities of the different metastable states (see Supplementary Methods Section 14.2, Supplementary equation (19)). We observed, as expected, that the less structured metastable states have a higher propensity to aggregate, be degraded or interact with chaperones. For example, relative to the native state, states P1 through P8 of CAT-III (see structures in Supplementary Fig. 5) are much more likely to experience one of these processes, as indicated by the larger magnitude of the bar plots in Fig. 2d. However, states P9 to P13 exhibit similar propensities as the native state (P14) to aggregate, be degraded, or become chaperone substrates. Similar results are observed for DDLB (Fig. 3d) and DHFR (Fig. 4d). Thus, our model indicates that some of these entangled structures do not interact with the proteostasis machinery any more than the native state does. Therefore, the altered specific activity we observe is likely to persist inside cells over long-time scales. In effect, this prediction is confirmed by our enzyme kinetic assays (see Section “Accurate prediction of trends in specific activity”), since the fast and slow variants of DDLB were expressed in *E. coli* cells possessing the full complement of the proteostasis machinery (see Supplementary Methods Section 19).

We estimated the percent of proteins in each metastable state that are likely to remain soluble by accounting for the aggregation, degradation and Hsp70 binding propensities within each metastable state (see Supplementary Methods Section 14.2, Supplementary equation (17)). We find that for both CAT-III and DDLB, many of the near-native entangled structures remained soluble. At least 97% of the proteins in entangled states P10, P11, P12 and P13 for CAT-III are estimated to remain soluble (Fig. 2e), while at least 89% of the proteins in entangled states P4, P6, P7, P8 and P9 of DDLB are expected to remain soluble (Fig. 3e). Thus, many of the long-lived, near-native entangled states are likely to remain soluble and free from aggregation, degradation, and catalyzed folding by chaperones.

The reason for this is that these near-native structures sequester the residues and sequence motifs that promote these processes to an extent similar to that in the native state. For example, state P13 of CAT-III exposes a similar amount of hydrophobic surface area as the native state ensemble (36.2 nm^2^ versus 34.7 nm^2^). The exposed surface areas of residues that promote aggregation and interactions with the chaperone Hsp70 are also similar between state P13 and the native state, respectively, 21.6 nm^2^ versus 18.9 nm^2^ and 40.0 nm^2^ versus 38.6 nm^2^.

### Limited proteolysis mass spectrometry refolding experiments are consistent with long-lived entangled states

To experimentally test if these entangled states exist, we carried out limited-proteolysis mass spectrometry^42^ (LiP-MS) in which *E. coli* lysates were globally unfolded through incubation in 6 M guanidinium chloride, refolded by dilution, and the conformations of the resulting refolded proteins assessed by their susceptibility to proteolysis with proteinase K (PK). With liquid chromatography tandem mass spectrometry (LC-MS/MS), tens of thousands of fragments were identified and quantified^43^, of which a number were from DDLB and DHFR, enabling an assessment of whether their refolded conformations were similar or not to their native forms. In these experiments, proteins were allowed to refold for 1 min, 5 min, or 2 h following dilution from denaturant, and then pulse proteolysis was conducted, providing a snapshot of their structural ensemble at distinct timescales. Peptide fragments that contain a cleavage site arising from proteinase K are interpreted as demarcating sites within a protein that are solvent-exposed or unstructured; hence, if such a fragment is present in greater abundance in refolded samples (relative to untreated, native samples), it implies that a population of the protein failed to form native-like structure at that site. Additional details can be found in Supplementary Methods Section 21. For DDLB, we find one statistically significant peptide fragment (abundance ratio > 2-fold, p-value < 0.01 by Welch’s t-test), with a proteinase K cleavage site at residue 192, whose abundance is at least 2.6 times greater in the refolded samples at all three time points (Table in Fig.3h, full dataset is provided in Supplementary Table 13), indicating residue 192 is more exposed to solvent than in the native state. Cross-referencing this site against the long-lived metastable states in our simulations (Fig. 3g), we find state P4 contains an entanglement in which residue 192 is part of the threading segment and more exposed to solvent (Fig. 3h). This misfolding results in a 12-fold increase of solvent accessible surface area for residue 192 compared to the native state in which it is part of a β-sheet. Thus, our simulations provide a molecular interpretation of LiP-MS refoldability experiments, and are consistent with the existence of protein entangled states.

In contrast, we predicted DHFR does not exhibit long lived misfolded states, and indeed in the LiP-MS data there is a peptide that exhibits a significant abundance difference in the refolded samples after 5 minutes of refolding time, however, it is not present at the 1 minute and 2-hour time points (see Supplementary Table 14), indicating the refolded protein is indistinguishable from its native conformation. Thus any structural distortions in DHFR during refolding are short-lived. This negative control provides further evidence of the accuracy of our model’s predictions.

### Synonymous mutations alter the post-translational populations of entangled states

Next, we examined how translation speed changes affect the post-translational populations of entangled structures. We observe that for CAT-III, 60 s after translation termination, 76.3% (95% CI: [73.6%, 78.9%]) of the structures are entangled when synthesis was fast, while 71.6% (95% CI: [68.8%, 74.3%]) are entangled when synthesis was slow. Likewise, the entangled population of DDLB accounted for 35.7% (95% CI: [32.7%, 38.7%]) and 30.2% (95% CI: [27.3%, 33.0%]) of the total under slow and fast translation, respectively. In contrast, the population of entangled structures for DHFR remained zero regardless of translation speed 10 s post termination. Thus, synonymous mutations can alter the population distributions of entangled states over long-time scales for deep entanglements.

### Synonymous mutations cause a divergence in the distribution of co-translational structures that persist post-translationally

The post-translational changes in the distribution of entangled structures must have originated co-translationally. We hypothesized that the changes in translation speed alter the co-translational folding pathways of the protein. To test this hypothesis, we first assessed how different were the nascent chain structural distributions under fast and slow translation. To do this, we applied the Jensen-Shannon (JS) divergence metric (Supplementary equation (21)) to the population of microstates identified as part of our Markov state modeling. A value of 0 means there is no difference between the distributions, while a value of ln2 means the distributions are completely different. Calculating this metric at each nascent chain length during synthesis, we found that for all three proteins, the structural divergence induced by synonymous mutations was small at short nascent chain lengths and tended to increase as the nascent chain length increased (Fig. 6b, e and h). The maximum divergence occurred at or near the longest nascent chain length before the nascent chain was released from the ribosome. Specifically, for DHFR, the structural ensembles arising from fast and slow synthesis started to diverge at approximately 110 residues in length, while for CAT-III and DDLB, the two distributions start to diverge at 190 and 210 residues, respectively. Thus, the fast- and slow-translation rates altered the co-translational distributions of conformations for all three enzymes.

**Figure 6.**
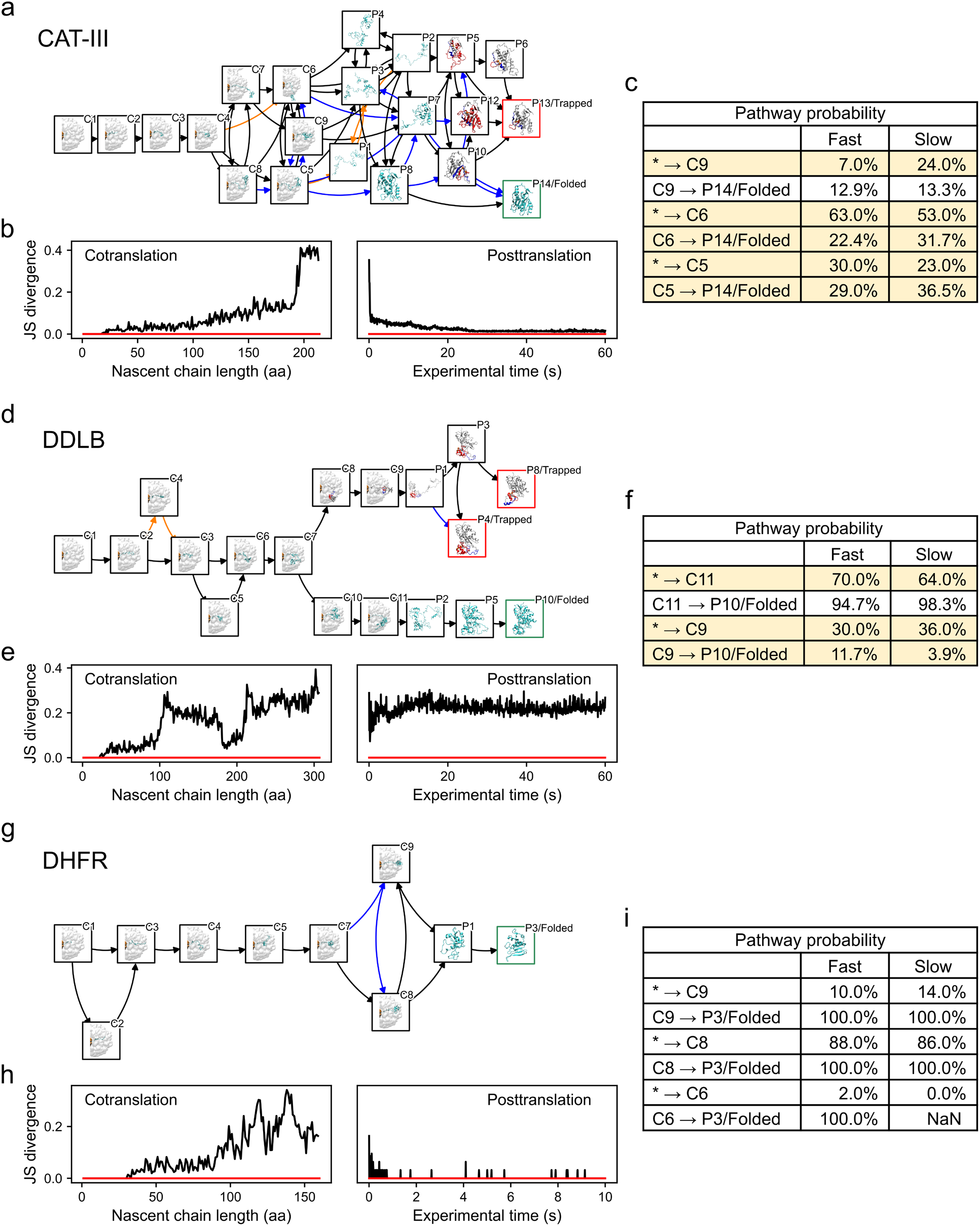
Co- and post-translational folding pathways of CAT-III, DDLB and DHFR. a, d, and g, The most probable folding pathways are presented as a directed graph where the nodes represent metastable states, with representative structures shown in the boxes, and the edges represent the transitions in the 80% most likely pathways. The nodes corresponding to the native state and the kinetically trapped states are framed in green and red, respectively. The transitions (edges) that are observed in only the fast variants or the slow variants are marked in orange and blue, respectively. b, e and h, Jensen-Shannon (JS) divergence of the co-translational microstate distributions (Supplementary equation (21)) and post-translational metastable state distributions (Supplementary equation (22)) comparing fast and slow variants, with a red line indicating zero divergence. c, f and i, The co-translational pathway probabilities (* → end state C*) and the post-translational pathway probabilities from the co-translational end states to the native state (C* → native state) are summarized in the tables. The pathway probabilities whose changes were greater than 5% are highlighted in yellow. The post-translational pathway probabilities are normalized by the corresponding co-translational pathway probabilities that involve the particular co-translational end state.

Applying this divergence metric to the time-dependent post-translational (metastable states) structural distributions (Supplementary equation (22)), we observed that for DHFR, the initially different structural ensembles arising from fast and slow synthesis rapidly converged to the same distribution post-translationally (*i.e.*, have divergence values equal to zero in Fig. 6h). For CAT-III and DDLB, the two distributions do not reconverge 60 s after synthesis (0.02 for CAT-III and 0.27 for DDLB, see Fig. 6b and e). Thus, the changes in conformations caused by synonymous mutations are quickly ‘forgotten’ by DHFR due to its rapid equilibration, while CAT-III and DDLB retain a ‘memory’ of those co-translational differences due to kinetic trapping.

### Changes in co-translational folding pathways give rise to altered post-translational populations

Next, we characterized the co- and post-translational folding pathways arising from changes in the translation rate. To do this, we calculated the populations and pathway probabilities of transitions between metastable states occurring co- and post-translationally under the different translation schedules (see Supplementary Methods Section 17). To display similarities and differences in folding pathways, we used arrows with different colors. Black arrows between metastable states indicate that a transition was observed between those states in the top 80% of populated pathways, blue arrows indicate that the transition was seen only in the slow-translation schedule, and orange arrows indicate that the transition was seen only in the fast translation schedule.

We identified nine co- and three post-translational metastable states (representative structures shown in Fig. 6g) for DHFR. The co- and post-translational folding pathways were very similar, as most of the transitions were seen in both translation schedules (black arrows in Fig. 6g). There were some differences though. Transitions from state C7 to C9 and from C9 to C8 were observed for only the slow schedule. The pathway probabilities, however, demonstrated that even though the initial post-translational structural distribution differed, all states converted to the native state by the end of the post-translational simulations (Fig. 6i). For example, the pathway probabilities of C8→P3 and C9→P3 are 100% for both fast and slow translation. An exception is the transition probability of C6→P3, where state C6 was not sampled by the slow variant, and no such transition was observed in our simulation. However, due to that all C6 ultimately converted to the native state P3 in the fast variant, this difference was negligible. Thus, the co-translational folding of DHFR was slightly affected by differences in translation speed; however, these differences quickly disappeared due to rapid folding from all post-translational metastable states.

In contrast, CAT-III exhibited a co- and post-translational folding reaction network (Fig. 6a) and pathway probabilities that were sensitive to translation speed changes and whose resulting differences persisted post-translationally. Nine co-translational and fourteen post-translational metastable states were identified for CAT-III (Fig. 6a). Six of these post-translational metastable states (P5, P6, P9, P10, P11, P12 and P13) exhibited entangled structures, while no co-translational states exhibited entanglement. Thus, entanglement was a post-translational process for CAT-III (see Supplementary videos I and II visualizing, respectively, the process of entanglement versus correct folding). Very different co-translational folding pathways were observed in the top 80% of pathways starting at a nascent chain length of approximately 195 residues, where the divergence metric exhibited a large increase (Fig. 6b), and transitions into and out of co-translational metastable state C9 started to differ between the fast and slow translation schedules (as indicated by the blue and orange arrows in Fig. 6a). For example, only in the slow schedule can C9 transition to C6, while only in the fast schedule can C5 transition to P1. These differences in allowed transitions persisted post-translationally as well, with the state P13 being an effective sink during the post-translational simulations, *i.e.*, allowing no direct or indirect transitions to the native state (P14) once they were reached. This sink has deep entangled structures, which is consistent with our earlier observation that the deep entangled states tend to be kinetic traps (indicated by the red border around metastable state P13 in Fig. 6a). Thus, the translation speed causes differences in the CAT-III co-translational folding pathways once at least 195 residues have been synthesized, and those differences lead to changes in the populations of post-translationally entangled states, thereby altering the transition state barrier for the reaction this enzyme carries out (Fig. 2f) and affecting the specific activity of CAT-III over long time scales (Fig. 1g).

Finer-grained consideration of the folding pathways shows that post-translationally, CAT-III_fast_ partitioned 5.0% (95% CI [1.1%, 8.9%]) more protein (gross amount, soluble + insoluble, same hereafter) into the near-native kinetic trap P13, while CAT-III_slow_ partitioned 4.7% (95% CI [0.8%, 8.6%]) more protein into native state P14. The transitions towards the native state P14 were enhanced for CAT-III_slow_, whereas the transitions towards the kinetic trap P13 were enhanced for CAT-III_fast_. This is also indicated by the pathway probabilities shown in Fig. 6c. Synonymous mutations also led to smaller changes in other states, including the entangled states P6, P10 and P12.

Similar to CAT-III, DDLB also exhibits co- and post-translational folding pathways that were sensitive to changes in translation speed across its eleven co- and ten post-translational metastable states. However, unlike CAT-III, DDLB co-translationally formed entangled structures (states C8 and C9) that persisted to form entangled post-translational states (states P1, P3, P4 and P8). In CAT-III, the divergence in co-translational folding pathways between fast and slow translation schedules monotonically increased with increasing chain length (Fig. 6b). However, the co-translational folding pathways of DDLB diverged starting at nascent chain length 110 but reconverged around 190 residues, diverging again starting at 210 residues (Fig. 6e). The first divergence involved a transition to state C4 with the fast schedule that did not occur with the slow schedule (orange arrow in Fig. 6d). This state ultimately interconverted to state C3, which is an obligatory intermediate in both schedules. The second divergence started at state C7, in which 6% more nascent chains went to entangled state C8 with the slow schedule than with the fast schedule, and the rest went to state C10. Parallel co- and post-translational folding pathways then arose. In one pathway, transitions were observed from C10→C11→P2, while the other transitions from C8→C9→P1 involved entangled structures in all states (see Fig. 6d, also see Supplementary videos III and IV illustrating the processes of entanglement and folding). The increased probability of this entangled pathway during slow translation decreased the probability that DDLB would post-translationally reach the native state P10 (3.9% versus 11.7% in slow versus fast, Fig. 6f), while DDLB_slow_ partitioned 5.8% (95% CI [3.2%, 8.3%]) more protein (gross amount, soluble + insoluble) into the near-native kinetic trap P4 after post-translation. Thus, we again see that entangled states act as sinks with altered transition state barriers for the enzyme’s reaction (Fig. 3f). This population shifts among these states and the folding pathways they take part in leads to long-lived changes in specific activity.

## Discussion

How synonymous mutations alter the specific activity of enzymes has been a challenging question. A detailed understanding of the molecular mechanisms giving rise to this phenomenon would provide biochemists, molecular biologists, evolutionary biologists, and biomedical researchers a framework to interpret experiments on the influence of codon usage on protein structure, function, cellular phenotype, disease, and the selection pressures shaping mRNA sequence evolution. In this study, we developed a novel multiscale model that qualitatively recapitulates the experimental observations for the specific activity of CAT-III variants^1^ and correctly predicts the trends in specific activity changes of DDLB and DHFR variants, giving us confidence that the model is realistic. This provided us the opportunity to study the structures, pathways, and kinetics that give rise to this phenomenon.

Our key findings are: (1) changes in elongation kinetics induced by synonymous mutations can alter co-translational nascent chain structural ensembles and folding pathways; (2) for some enzymes, such as CAT-III and DDLB, these changes in the structural ensemble can persist long after the nascent chain has been released from the ribosome; (3) this persistence arises from conformational states that are long-lived kinetic traps; (4) these kinetic traps arise at the molecular level from deep entanglements that slowly disentangle because they require the unfolding of already folded protein segments; (5) these entanglements are non-covalent lasso topologies in which a closed loop is formed by a backbone segment connecting two residues that form a non-covalent native contact, and another segment threads through this loop; (6) many of these entangled structures have decreased catalytic efficiencies due to structural perturbation of their active sites; (7) some entangled structures are very similar to the native state – exposing similar extents of hydrophobic surface area and chaperone binding motifs; and (8) because of their near-native conformations, these entangled structures can bypass the chaperone and degradation machinery of the cell and do not exhibit an increased propensity to aggregate. This situation results in a soluble fraction of enzymes with decreased specific activities that can persist for long time periods in cells.

Two concepts central to our explanation – intra-molecular entanglement and subpopulations of kinetically trapped states – are not without precedent. The material properties of entangled synthetic polymers have long been studied and modeled^44, 45^ in the field of polymer physics. And there has been a large amount of research focused on knotted proteins that often contain disulfide bonds or topologies in which when the protein’s ends are pulled in opposite directions the protein does not fully extend^31, 46, 47, 48^. These characteristics, however, are not present in our entanglements. The entanglements we observe form a closed loop due to a non-covalent native contact, and if both ends of the protein were pulled disentanglement would occur. Recently, this type of entanglement has been detected in almost one third of the protein structures deposited in the Protein Data Bank^32^. Thus, the potential for protein segments to form non-native, noncovalent lassos is plausible. Kinetically trapped states of proteins have been observed in single-molecule experiments probing the functioning of flavoenzymes^49, 50^. In one study, heterogeneous populations of the protein cholesterol oxidase were observed to stochastically switch very slowly between active and nonactive states^49^. However, in that study, the influence of protein synthesis on the distribution of conformational states was not probed. Thus, a novel aspect of our discovery is that it combines the phenomena of entanglement and kinetic trapping as essential to the mechanism by which synonymous mutations affect co- and post-translational protein structure and function.

We also observed in our simulations the counterintuitive phenomenon that the folding of some proteins is promoted when their rapid synthesis gives them less time to fold on the ribosome. This phenomenon was first predicted from chemical kinetic models^51^ of co-translational folding. Specifically, we observed that synthesizing DDLB three times faster increased the final post-translational folding probability of DDLB by 5.5% (95% CI [1.4%, 9.7%]) and increased its activity. Consistent with the earlier prediction^51^, this phenomenon arose because having less time to fold on the ribosome decreases DDLB partitioning into the misfolded, kinetically trapped state. This adds to the growing list of proteins for which slowing synthesis decreases folding efficiency, which results in altered function, and challenges the common rule of thumb that slow-translating codons promote folding.

An important aspect of our choice of protein systems is that we included a protein, DHFR, whose activity is not sensitive to synonymous mutations. This provided us with the opportunity to gain insights by comparing and contrasting its behavior with those of CAT-III and DDLB. A key insight from this comparison is that even though the DHFR co-translational structures and folding pathways were sensitive to synonymous mutations, these differences did not persist post-translationally because the only entangled state that DHFR populates and disentangled rapidly. This is consistent with previous experimental and theoretical studies that show that DHFR has fast folding kinetics and an absence of significant off-pathway intermediates^52^ and kinetic traps^53^. We demonstrated that the two reasons for this are that DHFR only negligibly populates an entangled conformation and that the entanglement it forms is ‘shallow’ - its disentanglement requires only 5 residues to slide out of the closed loop and does not need to unfold any other portions of the protein to do so. Thus, this shallow entanglement can be easily disentangled without large structural rearrangements. In contrast, CAT-III and DDLB entanglements have many more residues involved, requiring large structural rearrangements to become disentangled, leading to more-persistent entangled structures. Thus, the presence of entanglement is not sufficient to guarantee a kinetic trap. What is important is the nature of the entanglement and the native structure surrounding it.

Like any simulation study, limitations of force fields, time scales, and sampling statistics are of concern. In the coarse-grained simulations, we used a structure-based force field to encode the native state as the global free energy minimum. One of the many benefits of this is that we automatically know that the entangled structures we observe are metastable states, as the native state is encoded to be the global free energy minimum below the protein’s melting temperature. However, one potential drawback is that this force field does not allow ordered, nonnative secondary and tertiary structures to form (*e.g.*, the model does not allow β-sheets to switch to α-helical bundles). This means that additional structures beyond entanglement could also contribute to the influence of synonymous mutations on enzyme activity. At the monomeric protein level, however, such nonnative ordered structure seems unlikely. Except for fold-switching proteins, which are rare in nature (only 0.5 to 4% of proteins in the Protein Data Bank are estimated to switch folds^54^) and can require quaternary interactions to induce structural changes^55^, there have been no reported experimental data for alternative, ordered tertiary structures occurring on the folding pathways of monomeric proteins in bulk solution. Also, with absence of attractive nonnative interactions in our model, the diversity of metastable states might be underestimated.

However, this is not likely to change our conclusions because non-native interactions increase the roughness of the energy landscape making kinetic trapping more likely. In the QM/MM simulations, we used the 3rd-order density-functional tight-binding (DFTB) method^56^ as a balance between computational speed and accuracy. Transition state barrier heights can change depending on the level of theory and basis set used. However, it is often the case that trends across related compounds or conformations are robust. Therefore, the differences in the barrier heights between metastable states are likely to be qualitatively correct, meaning our overall conclusions would remain unchanged even if a higher level of theory was used.

Another limitation of our simulations is that we only simulated one protein chain at a time. Thus, we can never observe homodimer-swapped structures that could also be long-lived kinetic traps and influence function^57^. However, as noted, in the experimental examples provided in the Introduction, such dimer-swapped structures were ruled out. To address sampling issues, we applied rigorous statistical analyses of all our reported quantities, including confidence intervals and p-values. All our conclusions were drawn from statistically significant signals, indicating that sampling was not an issue. For these reasons, our results are robust and statistically meaningful, giving us confidence that they represent realistic scenarios of what is happening at the molecular level.

The predictions that monomeric enzymes can become intramolecularly entangled and populate long-lived states can be tested experimentally. Ensemble-level experiments, such as NMR and X-ray crystallography, often need appreciable populations of a state to detect that state, which would make entanglement detection difficult. Many biophysical techniques used in protein folding (such as FRET, tryptophan fluorescence, and circular dichroism) are sensitive to sub-populations, but tend to have low structural resolution, which would limit their capacity to distinguish near-native conformations. Therefore, hydrogen-deuterium exchange mass spectrometry as well as LiP-MS (as we show here), which can localize conformational differences to specific regions within a protein in a temporally-resolved manner, seem the most promising techniques to experimentally detect these entanglements. Cryo-EM, with its ability to build structural classes from heterogeneous populations, would also be effective for systems of suitable size and resolution.

It is reasonable to expect that the enzymatic activity of initially misfolded proteins should increase with time as the protein relaxes to its native state. Based on the disentangling timescale of 10^4^ s (see Section “Deep entanglements are usually long-lived regardless of model resolution”), we estimate that it would take more than 3 hours for the specific activities between fast and slow variants to converge. This is consistent with what was found for CAT-III, where the specific activity of synonymous mutant CAT-III didn’t converge to that of the wild type within 20 minutes^1^. For DDLB, because the limited-proteolysis mass spectrometry results confirmed that the misfolded states can persist longer than 2 hours, it is reasonable to anticipate that the specific activities between fast and slow DDLB variants will take more than 2 hours to converge, which is also consistent with our prediction. Further experiments, such as a time-dependent activity assay coupled with pulse-chase protein expression and production, could be applied to measure such timescales.

This study provides a plausible explanation of how synonymous mutations can alter enzyme activity in cells. Synonymous mutations alter translation elongation speeds and change the population of nascent chain conformations in entangled states that are near native but have lower catalytic efficiencies than that of the native state. Hence, the specific activity, a quantity averaged over the populations of proteins in different conformational states, can increase or decrease due to synonymous mutations. The experimental search for these entangled structures and their roles in influencing protein structure, function, and phenotypes in cells is likely to be a fruitful area of research in the future.

## Acknowledgements

S.D.F. acknowledges support from the NIH Director’s New Innovator Award (DP2GM140926) and from the National Science Foundation Division of Molecular and Cellular Biology (MCB-2045844).

S.J.B. acknowledges support from the NIH (GM-122595), the Eberly Family Distinguished Chair in Science, and the Howard Hughes Medical Institute. E.P.O. acknowledges support from the National Science Foundation (MCB-1553291) as well as the National Institutes of Health (R35-GM124818). Computations in this work have been carried out on the Extreme Science and Engineering Discovery Environment (XSEDE) supercomputer^58^, which is supported by MCB-160069, and the Pennsylvania State University’s Institute for Computational and Data Sciences’ Roar supercomputer.

## Author contributions

E.P.O. designed the research; Y.J. developed the computational methods with contributions from E.P.O.; Y.J. wrote the computer code and carried out the simulations and computations; S.S.N. and S.J.B. designed the experimental validation for DDLB variants; S.S.N., I.S., E.P.O and S.J.B. designed the experimental validation for DHFR variants; S.S.N., I.S. and P.P. performed the specific activity experiments; S.D.F. designed the limited-proteolysis mass spectrometry experiments for DDLB and DHFR; P.T. and Y.X. performed the limited-proteolysis mass spectrometry experiments; All of the authors analyzed the data and wrote the manuscript.

## Code availability

All computer code developed in this work is available in the GitHub repositories https://github.com/obrien-lab/cg_simtk_protein_folding and https://github.com/obrien-lab/Activation-Energy-Estimation-Workflow, under the MIT license. Detailed instructions of code usage, basic theory and examples of the input/output are available in the wiki pages of the above repositories.

## Data and biological materials availability

Raw data for Figures 1 to 6 are available in the supplementary data. The input data that we used to perform the simulations in this study are available in the repository subdirectory https://github.com/obrien-lab/cg_simtk_protein_folding/blob/master/example/input_data.tar.xz.

All the data that support the findings of this study, as well as the biological materials that were used for testing the enzymatic activity of DDLB and DHFR variants, are available from the corresponding author upon reasonable request. The raw mass spectrometry data of DDLB and DHFR have been deposited to the ProteomeXchange Consortium via the PRIDE partner repository with the dataset identifier PXD031425.

## Graphical Abstract

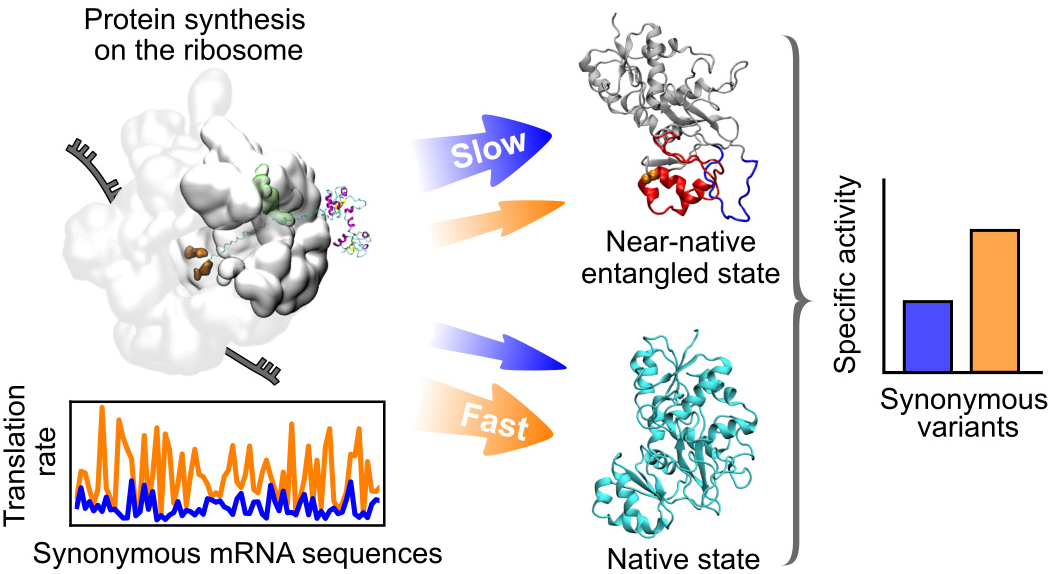

## Notes

### Competing Interest Statement

The authors have declared no competing interest.

## References

1. Komar AA, Lesnik T, Reiss, C. Synonymous codon substitutions affect ribosome traffic and protein folding during in vitro translation. FEBS Lett 1999, 462(3): 387–391.

2. Zhao F, Yu C-h, Liu, Y. Codon usage regulates protein structure and function by affecting translation elongation speed in Drosophila cells. Nucleic Acids Res 2017, 45(14): 8484–8492.

3. Spencer PS, Siller E, Anderson JF, Barral, JM. Silent substitutions predictably alter translation elongation rates and protein folding efficiencies. J Mol Biol 2012, 422(3): 328–335.

4. Hunt R, Hettiarachchi G, Katneni U, Hernandez N, Holcomb D, Kames J, et al. A single synonymous variant (c. 354G> A [p. P118P]) in ADAMTS13 confers enhanced specific activity. International journal of molecular sciences 2019, 20(22): 5734.

5. Crombie T, Boyle JP, Coggins JR, Brown, AJ. The folding of the bifunctional TRP3 protein in yeast is influenced by a translational pause which lies in a region of structural divergence with Escherichia coli indoleglycerol‐phosphate synthase. Eur J Biochem 1994, 226(2): 657–664.

6. Walsh IM. Testing the Effects of Synonymous Codon Usage on Co-Translational Protein Folding Using Novel Experimental and Computational Techniques. University Of Notre Dame, 2019.

7. Yu C-H, Dang Y, Zhou Z, Wu C, Zhao F, Sachs MS, et al. Codon usage influences the local rate of translation elongation to regulate co-translational protein folding. Mol Cell 2015, 59(5): 744–754.

8. Walsh IM, Bowman MA, Santarriaga IFS, Rodriguez A, Clark, PL. Synonymous codon substitutions perturb cotranslational protein folding in vivo and impair cell fitness. Proc Natl Acad Sci 2020, 117(7): 3528–3534.

9. Sala AJ, Bott LC, Morimoto, RI. Shaping proteostasis at the cellular, tissue, and organismal level. J Cell Biol 2017, 216(5): 1231–1241.

10. Liu Y, Tan YL, Zhang X, Bhabha G, Ekiert DC, Genereux JC, et al. Small molecule probes to quantify the functional fraction of a specific protein in a cell with minimal folding equilibrium shifts. Proc Natl Acad Sci 2014, 111(12): 4449–4454.

11. Buhr F, Jha S, Thommen M, Mittelstaet J, Kutz F, Schwalbe H, et al. Synonymous codons direct cotranslational folding toward different protein conformations. Mol Cell 2016, 61(3): 341–351.

12. Martelli PL, Fariselli P, Casadio, R. Prediction of disulfide‐bonded cysteines in proteomes with a hidden neural network. Proteomics 2004, 4(6): 1665–1671.

13. O’Brien EP, Christodoulou J, Vendruscolo M, Dobson, CM. Trigger factor slows co-translational folding through kinetic trapping while sterically protecting the nascent chain from aberrant cytosolic interactions. J Am Chem Soc 2012, 134(26): 10920–10932.

14. Sharma AK, Bukau B, O’Brien EP. Physical origins of codon positions that strongly influence cotranslational folding: A framework for controlling nascent-protein folding. J Am Chem Soc 2016, 138(4): 1180–1195.

15. Fritch B, Kosolapov A, Hudson P, Nissley DA, Woodcock HL, Deutsch C, et al. Origins of the mechanochemical coupling of peptide bond formation to protein synthesis. J Am Chem Soc 2018, 140(15): 5077–5087.

16. Nissley DA, O’Brien EP. Structural Origins of FRET-Observed Nascent Chain Compaction on the Ribosome. J Phys Chem B 2018, 122(43): 9927–9937.

17. Leininger SE, Trovato F, Nissley DA, O’Brien EP. Domain topology, stability, and translation speed determine mechanical force generation on the ribosome. Proc Natl Acad Sci 2019, 116(12): 5523–5532.

18. Nissley DA, Vu QV, Trovato F, Ahmed N, Jiang Y, Li MS, et al. Electrostatic interactions govern extreme nascent protein ejection times from ribosomes and can delay ribosome recycling. J Am Chem Soc 2020, 142(13): 6103–6110.

19. Dunkle JA, Wang L, Feldman MB, Pulk A, Chen VB, Kapral GJ, et al. Structures of the bacterial ribosome in classical and hybrid states of tRNA binding. Science 2011, 332(6032): 981–984.

20. Arenz S, Bock LV, Graf M, Innis CA, Beckmann R, Grubmüller H, et al. A combined cryo-EM and molecular dynamics approach reveals the mechanism of ErmBL-mediated translation arrest. Nat Commun 2016, **7:** 12026.

21. Sharma AK, Sormanni P, Ahmed N, Ciryam P, Friedrich UA, Kramer G, et al. A chemical kinetic basis for measuring translation initiation and elongation rates from ribosome profiling data. PLoS Comp Biol 2019, 15(5): e1007070.

22. Fluitt A, Pienaar E, Viljoen, H. Ribosome kinetics and aa-tRNA competition determine rate and fidelity of peptide synthesis. Comput Biol Chem 2007, 31(5-6): 335–346.

23. Eastman P, Swails J, Chodera JD, McGibbon RT, Zhao Y, Beauchamp KA, et al. OpenMM 7: Rapid development of high performance algorithms for molecular dynamics. PLoS Comp Biol 2017, 13(7): e1005659.

24. Nagano N. EzCatDB: the enzyme catalytic-mechanism database. Nucleic Acids Res 2005, 33(suppl_1): D407–D412.

25. Nagano N, Nakayama N, Ikeda K, Fukuie M, Yokota K, Doi T, et al. EzCatDB: the enzyme reaction database, 2015 update. Nucleic Acids Res 2014, 43(D1): D453–D458.

26. Consortium U. UniProt: the universal protein knowledgebase. Nucleic Acids Res 2018, 46(5): 2699.

27. Berman HM, Westbrook J, Feng Z, Gilliland G, Bhat TN, Weissig H, et al. The protein data bank. Nucleic Acids Res 2000, 28(1): 235–242.

28. MacQueen J. Some methods for classification and analysis of multivariate observations. Proceedings of the fifth Berkeley symposium on mathematical statistics and probability; 1967: Oakland, CA, USA; 1967. p. 281–297.

29. Röblitz S, Weber, M. Fuzzy spectral clustering by PCCA+: application to Markov state models and data classification. Advances in Data Analysis and Classification 2013, 7(2): 147–179.

30. Niemyska W, Dabrowski-Tumanski P, Kadlof M, Haglund E, Sułkowski P, Sulkowska, JI. Complex lasso: new entangled motifs in proteins. Sci Rep 2016, 6: 36895.

31. Sulkowska JI. On folding of entangled proteins: knots, lassos, links and θ-curves. Curr Opin Struct Biol 2020, 60: 131–141.

32. Baiesi M, Orlandini E, Seno F, Trovato, A. Sequence and structural patterns detected in entangled proteins reveal the importance of co-translational folding. Sci Rep 2019, 9(1): 1–12.

33. Baiesi M, Orlandini E, Seno F, Trovato, A. Exploring the correlation between the folding rates of proteins and the entanglement of their native states. Journal of Physics A: Mathematical and Theoretical 2017, 50(50): 504001.

34. Connolly ML, Kuntz I, Crippen, GM. Linked and threaded loops in proteins. Biopolymers: Original Research on Biomolecules 1980, 19(6): 1167–1182.

35. Jarmolinska AI, Gambin A, Sulkowska, JI. Knot_pull—python package for biopolymer smoothing and knot detection. Bioinformatics 2020, 36(3): 953–955.

36. Jennings PA, Finn BE, Jones BE, Matthews, CR. A reexamination of the folding mechanism of dihydrofolate reductase from Escherichia coli: verification and refinement of a four-channel model. Biochemistry 1993, 32(14): 3783–3789.

37. Garbuzynskiy SO, Ivankov DN, Bogatyreva NS, Finkelstein, AV. Golden triangle for folding rates of globular proteins. Proc Natl Acad Sci 2013, 110(1): 147–150.

38. Aquilanti V, Mundim KC, Elango M, Kleijn S, Kasai, T. Temperature dependence of chemical and biophysical rate processes: Phenomenological approach to deviations from Arrhenius law. Chem Phys Lett 2010, 498(1-3): 209–213.

39. Wallace MI, Ying L, Balasubramanian S, Klenerman, D. Non-Arrhenius kinetics for the loop closure of a DNA hairpin. Proc Natl Acad Sci 2001, 98(10): 5584–5589.

40. Onuchic JN, Luthey-Schulten Z, Wolynes, PG. Theory of protein folding: the energy landscape perspective. Annu Rev Phys Chem 1997, 48(1): 545–600.

41. Sarkar D, Kang P, Nielsen SO, Qin, Z. Non-Arrhenius Reaction-Diffusion Kinetics for Protein Inactivation over a Large Temperature Range#. ACS nano 2019, 13(8): 8669–8679.

42. Feng Y, De Franceschi G, Kahraman A, Soste M, Melnik A, Boersema PJ, et al. Global analysis of protein structural changes in complex proteomes. Nat Biotechnol 2014, 32(10): 1036–1044.

43. Nissley DA, Jiang Y, Trovato F, Sitarik I, Narayan K, To P, et al. Universal protein misfolding intermediates can bypass the proteostasis network and remain soluble and less-functional. bioRxiv 2022: 2021.2008.2018.456613.

44. Kröger M. Developments in Polymer Theory and Simulation. Polymers (Basel) 2019, 12(1):30.

45. Pawlak A. The Entanglements of Macromolecules and Their Influence on the Properties of Polymers. Macromol Chem Phys 2019, 220(10): 1900043.

46. Sułkowska JI, Sułkowski P, Onuchic, J. Dodging the crisis of folding proteins with knots. Proc Natl Acad Sci 2009, 106(9): 3119–3124.

47. Haglund E, Sulkowska JI, Noel JK, Lammert H, Onuchic JN, Jennings, PA. Pierced lasso bundles are a new class of knot-like motifs. PLoS Comput Biol 2014, 10(6): e1003613.

48. Haglund E, Sułkowska JI, He Z, Feng G-S, Jennings PA, Onuchic, JN. The unique cysteine knot regulates the pleotropic hormone leptin. PLoS One 2012, 7(9): e45654.

49. Lu HP, Xun L, Xie, XS. Single-molecule enzymatic dynamics. Science 1998, 282(5395): 1877–1882.

50. Yang H, Luo G, Karnchanaphanurach P, Louie T-M, Rech I, Cova S, et al. Protein conformational dynamics probed by single-molecule electron transfer. Science 2003, 302(5643): 262–266.

51. O’Brien EP, Vendruscolo M, Dobson, CM. Kinetic modelling indicates that fast-translating codons can coordinate cotranslational protein folding by avoiding misfolded intermediates. Nat Commun 2014, 5(1): 1–11.

52. Heidary DK, O’Neill JC, Roy M, Jennings, PA. An essential intermediate in the folding of dihydrofolate reductase. Proc Natl Acad Sci 2000, 97(11): 5866–5870.

53. Bitran A, Jacobs WM, Zhai X, Shakhnovich, E. Cotranslational folding allows misfolding-prone proteins to circumvent deep kinetic traps. Proc Natl Acad Sci 2020, 117(3): 1485–1495.

54. Porter LL, Looger, LL. Extant fold-switching proteins are widespread. Proc Natl Acad Sci 2018, 115(23): 5968–5973.

55. Bryan PN, Orban, J. Proteins that switch folds. Curr Opin Struct Biol 2010, 20(4): 482–488.

56. Gaus M, Cui Q, Elstner, M. DFTB3: extension of the self-consistent-charge density-functional tight-binding method (SCC-DFTB). J Chem Theory Comput 2011, 7(4): 931–948.

57. Liu Y, Eisenberg, D. 3D domain swapping: as domains continue to swap. Protein Sci 2002, 11(6): 1285–1299.

58. Towns J, Cockerill T, Dahan M, Foster I, Gaither K, Grimshaw A, et al. XSEDE: accelerating scientific discovery. Computing in science & engineering 2014, 16(5): 62–74.

