## Supplementary Information for "How synonymous mutations alter enzyme structure and function over long time scales"

### Supplementary Methods

#### 1. Coarse-grained (CG) protein model

We utilized a Gō-based force field[1, 2, 3, 4, 5] for all the proteins investigated in this study. Briefly, this CG model represents each residue as a single interaction site centered on the Cα atom position. The potential energy function is:

$$
\begin{aligned}
E_{\text{tot}} &= E_{\text{bond}} + E_{\text{angle}} + E_{\text{dihedral}} + E_{\text{elec}} + E_{\text{vdW}} \\
&= \sum_i K_{\text{b}}(b_i - b_0)^2 + \sum_i -\frac{1}{\gamma}\ln\left\{e^{-\gamma[K_\alpha(\theta_i-\theta_\alpha)^2+\varepsilon_\alpha]} + e^{-\gamma K_\beta(\theta_i-\theta_\beta)^2}\right\} \\
&\quad + \sum_i \sum_{j=1}^{4} K_{\text{D}_j}\left[1+\cos\left(j\varphi_i - \delta_j\right)\right] + \sum_{i,j} \frac{q_i q_j}{4\pi\varepsilon_0\varepsilon_\gamma r_{ij}} \cdot e^{-\frac{r_{ij}}{l_{\text{D}}}} \\
&\quad + \sum_{i,j} \varepsilon_{ij}\left[13\left(\frac{R_{ij}}{r_{ij}}\right)^{12} - 18\left(\frac{R_{ij}}{r_{ij}}\right)^{10} + 4\left(\frac{R_{ij}}{r_{ij}}\right)^{6}\right], \qquad (1)
\end{aligned}
$$

where $K_{\text{b}}$ is the bond force constant and equals 50 kcal/mol/Å$^2$; $b_i$ is the $i^{\text{th}}$ pseudo bond length between two adjacent interaction sites; $b_0$ is the equilibrium pseudo bond length and is set to 3.81 Å, which is the average distance between two adjacent Cα atoms in the protein sequence; $\gamma$, $K_\alpha$, $\theta_\alpha$, $\varepsilon_\alpha$, $K_\beta$ and $\theta_\beta$ are all constants of the double-well angle potential[6], which was designed to describe bond angles associated with both α-helix and β-sheet conformations; $\theta_i$ is the $i^{\text{th}}$ angle of two adjacent pseudo bonds; $K_{\text{D}_j}$ and $\delta_j$ are the dihedral force constant and the phase at periodicity $j$, respectively; $\varphi_i$ is the $i^{\text{th}}$ pseudo dihedral angle; $q_i$ is the net charge of the $i^{\text{th}}$ interaction site, which equals the net charge of the corresponding amino acid residue; $\varepsilon_0$ and $\varepsilon_\gamma$ are the dielectric constants of vacuum and water, respectively; $l_{\text{D}}$ is the Debye length and is set[1] to 10 Å; $\varepsilon_{ij}$ and $R_{ij}$ are the well depth and the vdW radius in the LJ 12-10-6 potential[7], which takes into account of desolvation barriers between interaction sites *i* and *j* and $r_{ij}$ is their distance. The 1-4 nonbonding interactions were included in our model without scaling any parameters. The nonbonding interactions were smoothly switched to zero starting at a distance of 18 Å and ending at 20 Å. Constraints on the bond length were applied in all the simulations.

Parameters $\varepsilon_{ij}$ and $R_{ij}$ were set according to the interaction types. For the interaction sites that have native contacts, $\varepsilon_{ij} = \varepsilon_{ij}^{\text{HB}} + n_{\text{scal}} \cdot \varepsilon_{ij}^{\text{SC-SC}} + \varepsilon_{ij}^{\text{BB-SC}}$, where the hydrogen bond (HB) contact well depth $\varepsilon_{ij}^{\text{HB}}$ was set as 0.75 kcal/mol for a single HB contact and 1.5 kcal/mol for multiple HB contacts; the sidechain-sidechain (SC-SC) interaction well depth $\varepsilon_{ij}^{\text{SC-SC}}$ was set according to the Betancourt−Thirumalai statistical potential[8] and then scaled by a multiplicative factor $n_{\mathrm{scal}}$ to achieve a realistic native-state stability for a particular protein; the backbone-sidechain (BB-SC) interaction well depth $\varepsilon_{ij}^{\text{BB-SC}}$ was set as 0.37 kcal/mol. All the native contacts were identified for the residue pairs that are separated by no less than 2 residues within the native structure. The native HB contacts were identified by STRIDE[9]. The native SC-SC contacts and BB-SC contacts were identified as those residue pairs that have any heavy atom within 4.5 Å in the crystal structure. The value of $R_{ij}$ for a native contact was set as the native distance $d_{ij}$ between interaction sites *i* and *j*. For the non-native contacts, to make the interaction mostly repulsive, we used $\varepsilon_{ij} = \sqrt{\varepsilon_i \cdot \varepsilon_j}$ and $R_{ij} = R_i + R_j$, where $\varepsilon_i$ was set as 0.000132 kcal/mol and $R_i$ was set as the non-native collision diameter $\sigma_i$ of interaction sites *i* multiplied by $2^{1/6}$ and divided by 2. The collision diameters were identified based on the protein's native structure according to Karanicolas−Brooks method[7]. All the CG force field parameters are summarized in Supplementary Table 1. All the CG simulations in this study were performed in OpenMM[10].

### 2. Defining native-state stabilities by parameterizing $n_{\mathrm{scal}}$

Setting a realistic free energy of stability is important for accurately modeling protein folding in the CG model. However, due to the time-consuming parameterization process, especially for a large multi-domain protein, it is impractical to parameterize all the proteins that we are interested in. We therefore utilized a training set that contains 18 small single-domain proteins to optimize each $n_{\mathrm{scal}}$ to reproduce the corresponding experimental protein stability[5, 11, 12, 13, 14] ($\Delta G_{\mathrm{UN}}^{\mathrm{exp}}$). The training-set proteins can be classified into 3 structural classes: mainly α-helix (marked as α class, see Supplementary Table 2a), mainly β-sheet (marked as β class, see Supplementary Table 2b) and α/β fold (marked as α/β class, see Supplementary Table 2c). We used the resulting $n_{\mathrm{scal}}$ data of those structural classes to further parameterize the proteins that we are interested in.

Each training-set protein was coarse-grained using the interaction sites at every Cα position within its crystal structure. The CG structures were then parameterized using 4 different $n_{\mathrm{scal}}$ values, resulting 4 CG models for each training-set protein. We utilized parallel temperature replica exchange molecular dynamic (pt-REMD)[15] simulations to efficiently sample the conformational space of each CG model. 100,000 replica exchange attempts were performed with the sampling time 75 ps between two attempts, resulting the total sampling time as 7.5 μs, for each CG model. The temperature windows for each pt-REMD simulations were chosen within the range of 280 K to 400 K and progressively optimized so that the successful exchange ratios are no less than 0.6 by using the minimum number of windows. The heat capacity $C_{\mathrm{v}}$ and the probability of protein folding $P_{\mathrm{N}}$ were then estimated using the Weighted Histogram Analysis Method[16] (WHAM). The melting temperature $T_{\mathrm{m}}$ was identified as the temperature where the $C_{\mathrm{v}}$

of the particular protein reaches its maxima. We used the fraction of native contact $Q$ within secondary structure elements as the order parameter (calculated using the main text equation (3), where *I* and *J* are both the set of residues within secondary structure elements) and the threshold $Q_{\mathrm{eq}}$ used to identify the folded and unfolded states was chosen as the $Q$ value where the cumulative probability reaches 0.5 at $T_{\mathrm{m}}$. The folded states were then identified as the structure whose $Q > Q_{\mathrm{eq}}$. The protein stability $\Delta G_{\mathrm{UN}}^{\mathrm{sim}}$ at the experimental temperature $T^{\mathrm{exp}}$ was then estimated by

$$\Delta G_{\mathrm{UN}}^{\mathrm{sim}}(T^{\mathrm{exp}}) = -k_{\mathrm{B}} T^{\mathrm{exp}} \cdot \ln\left[\frac{P_{\mathrm{N}}(T^{\mathrm{exp}})}{1 - P_{\mathrm{N}}(T^{\mathrm{exp}})}\right], \quad (2)$$

where $k_{\mathrm{B}}$ is Boltzmann's constant.

There are several proteins in α and α/β classes whose conformational space cannot be sufficiently sampled within the 100,000 exchange attempts, i.e., replicas cannot cross the free energy barrier to visit both of the folded and unfolded states frequently. To improve the reliability of the estimated protein stability, we performed an extra pt-REMD simulation at each $n_{\mathrm{scal}}$ value starting from the unfolded states that were obtained from the MD simulations at 800 K. The final $\Delta G_{\mathrm{UN}}^{\mathrm{sim}}$ at a particular $n_{\mathrm{scal}}$ value for those insufficiently sampled proteins (shown in Supplementary Table 3) were calculated as the average of the protein stability values estimated using the pt-REMD simulations starting from their native states and unfolded states.

The optimized $n_{\mathrm{scal}}$ for each training-set protein (shown in Supplementary Table 2) was determined through extrapolation as

$$n_{\mathrm{scal}}^{*} = \frac{\Delta G_{\mathrm{UN}}^{\mathrm{exp}} - b}{m}, \quad (3)$$

where *m* and *b* are the slope and intercept obtained by the linear regression of $\Delta G_{\mathrm{UN}}^{\mathrm{sim}}$ as a function of $n_{\mathrm{scal}}$, respectively.

#### 3. Stepwise $n_{\mathrm{scal}}$ optimization strategy for arbitrary proteins

To further parameterize the proteins that are more complex than the training set proteins, we used a stepwise optimization strategy. Five levels of $n_{\mathrm{scal}}$ for each structural class were predefined as shown in Supplementary Table 5. At the first level, $n_{\mathrm{scal}}$ was chosen as the mean value of the structural class $\langle n_{\mathrm{scal}}^{*} \rangle_{\mathrm{class}}$, as well as the overall mean value of the 18 training-set proteins for the domain interface. At the second level, $n_{\mathrm{scal}}$ was chosen as $\langle n_{\mathrm{scal}}^{*} \rangle_{\mathrm{class}}$ increased by $\langle \Delta\Delta G / \Delta G_{\mathrm{UN}}^{\mathrm{class}} \rangle_{-} \times 100\% = 23\%$, where $\Delta\Delta G = \Delta G_{\mathrm{UN}}^{\mathrm{exp}} - \Delta\Delta G_{\mathrm{UN}}^{\mathrm{class}}$, $\Delta\Delta G_{\mathrm{UN}}^{\mathrm{class}} = m \cdot \langle n_{\mathrm{scal}} \rangle_{\mathrm{class}} + b$

and $\langle \ldots \rangle_-$ means the average is only calculated for the proteins that are destabilized using $\langle n^*_{\text{scal}} \rangle_{\text{class}}$ within the training set. At the third level, $n_{\text{scal}}$ was chosen as $\langle n^*_{\text{scal}} \rangle_{\text{class}}$ increased by $\underline{\max}\{\Delta\Delta G/\Delta G^{\text{class}}_{\text{UN}}\} \times 100\% = 46\%$, where $\underline{\max}\{\ldots\}$ means the maximum value only accounts for the proteins that are destabilized using $\langle n^*_{\text{scal}} \rangle_{\text{class}}$ within the training set. At the fourth level, $n_{\text{scal}}$ was chosen as $\langle n_{\text{scal}} \rangle_{\text{class}}$ increased by $\min\{\Delta\Delta G\}/\langle \Delta G^{\text{class}}_{\text{UN}} \rangle \times 100\% = 70\%$, which is the largest destabilization relative to the overall average $\Delta G^{\text{class}}_{\text{UN}}$. At the last level, $n_{\text{scal}}$ was chosen as the overall maximum value of all levels.

An arbitrary protein was parameterized using the following strategy: The protein was coarsen-grained using our CG force field (Supplementary equation (1)). The native vdW interactions were divided into groups based on the domains defined by CATH[17]; $n_{\text{scal}}$ was assigned as the first-level value according to the domain's structural class (or interface). We then performed 10 parallel 1-μs MD simulations at 310 K for this CG model and monitor the *Q* value of each domain and interface. The domain or interface was regarded stabilized when all the 10 trajectories have the *Q* values greater than the threshold $\langle Q_{\text{kin}} \rangle$ for no less than 98% of the simulation time. We then increased the $n_{\text{scal}}$ values of the destabilized domains and interfaces to the next level and keep the other's $n_{\text{scal}}$ values invariant and generate the next CG model. We repeated the above procedure until all the domains and interfaces were stabilized. If a domain or interface cannot be stabilized even using the highest level of $n_{\text{scal}}$, the median $n_{\text{scal}}$ value (i.e., level 3) of the corresponding structural class was used to generate the final CG model regardless of the stability.

### 4. Estimation of *in silico* protein folding times

The *in silico* protein folding time $\langle \tau^{\text{sim}}_{\text{F}} \rangle$ was estimated as the mean first passage time (MFPT) of protein folding for each training-set protein by using temperature quenching simulations. We performed MD simulations for all the CG protein models built using corresponding $n^*_{\text{scal}}$ at 800 K to force them unfolded, followed by running MD simulations at 310 K to monitor protein folding. The trajectory where the *Q* value is higher than a threshold $Q_{\text{kin}}$ for at least 150 ps was identified as a folded trajectory and then terminated. $Q_{\text{kin}}$ is defined as $Q_{\text{kin}} = \frac{1}{2}\left(Q_{\text{eq}} + Q_{310}\right)$, where $Q_{310}$ is the most probable *Q* value at 310 K for each training-set protein estimated by using WHAM in the pt-REMD trial whose $n^*_{\text{scal}}$ is closest to the optimized value $n^*_{\text{scal}}$.

500 trajectories were run for each training-set protein and the proportion of unfolded trajectories ($S_{\text{U}}$) was fit to an exponential decay as a function of simulation time. We first

considered the simplest folding kinetics with one pathway $\mathrm{U} \overset{k_{\mathrm{F}}}{\rightarrow} \mathrm{F}$. In this case, $S_{\mathrm{U}}$ follows a single-exponential equation

$$S_{\mathrm{U}}(t) = \begin{cases} 1, & 0 \le t < t_0 \\ e^{-k_{\mathrm{F}}(t-t_0)}, & t \ge t_0 \end{cases}, \tag{4}$$

where $t_0$ is the lag time of protein folding. The single-exponential fitting works well except for protein ABP1, SH3, and Tenascin, whose fitting quality $R^2$ are both lower than 0.99. We further considered the protein folding kinetics that has two parallel pathways that start from two isolated unfolded states $U_1$ and $U_2$ (with extremely slow inter-transitions between $U_1$ and $U_2$ that can be neglected) to F with folding rates $k_1$ and $k_2$ for each. In this case, $S_{\mathrm{U}}$ follows the double-exponential equation

$$S_{\mathrm{U}}(t) = \begin{cases} 1 & 0 \le t < t_1 \\ f_1 e^{-k_1(t-t_1)} + f_2, & t_1 \le t < t_2 \\ f_1 e^{-k_1(t-t_1)} + f_2 e^{-k_2(t-t_2)}, & t \ge t_2 \end{cases}, \tag{5}$$

where $f_1$ and $f_2$ are the initial proportions of the unfolded states $U_1$ and $U_2$, respectively, and thus satisfy $0 < f_1, f_2 < 1$ and $f_1 + f_2 \equiv 1$; $t_1$ and $t_2$ are the corresponding lag times for the first and second folding pathway. This kinetic scheme separates the fast folding and slow folding events. Using the double-exponential equation, the fitting quality $R^2$ of protein ABP1, SH3, and Tenascin was above 0.99.

The *in silico* protein folding time $\langle \tau_{\mathrm{F}}^{\mathrm{sim}} \rangle$ was then estimated by

$$\begin{aligned} \langle \tau_{\mathrm{F}}^{\mathrm{sim}} \rangle &= \int_0^\infty \frac{d[1 - S_{\mathrm{U}}(t)]}{dt} \cdot t\,\mathrm{d}t \\ &= \begin{cases} t_0 + \dfrac{1}{k_{\mathrm{F}}}, & \text{for single-pathway kinetics;} \\ f_1\left(t_1 + \dfrac{1}{k_1}\right) + f_2\left(t_2 + \dfrac{1}{k_2}\right), & \text{for double-pathway kinetics;} \end{cases} \end{aligned} . \tag{6}$$

The scaling factor $\alpha$ of the *in silico* protein folding time $\langle \tau_{\mathrm{F}}^{\mathrm{sim}} \rangle$ with respect to the experimental protein folding time $\langle \tau_{\mathrm{F}}^{\mathrm{exp}} \rangle$ was calculated as $\alpha = \overline{\langle \tau_{\mathrm{F}}^{\mathrm{exp}} \rangle / \langle \tau_{\mathrm{F}}^{\mathrm{sim}} \rangle}$. All the relevant quantities for calculating $\langle \tau_{\mathrm{F}}^{\mathrm{sim}} \rangle$ and $\alpha$ are listed in Supplementary Table 4.

### 5. CG model of ribosome

The high-resolution crystal structure PDB 4v9d[18] was used to build a coarse-grain model of the *E. coli* ribosome. To model the tRNA molecules present in the A-site and P-site, we inserted the A- and P-site tRNAs obtained from PDB 5jte[19] into the A-site and P-site of PDB 4v9d by aligning the 50S subunit of PDB 4v9d to the 50S subunit of PDB 5jte. The entire 50S subunit, including 23S rRNA, 5S rRNA, ribosomal proteins, A-site tRNA and P-site tRNA, was coarsen-grained using the three/four-point model of RNA[1] and the Cα model of protein[1]. Briefly, nucleotides containing pyrimidines and purines were represented as 3 and 4 interaction sites, respectively, with one interaction site (P) located at the phosphate position and having a q = -1*e* charge, another (R) at the centroid of the ribose ring, and one (named as PU1 and PU2, or PY) at the centroid of each conjugated ring in the base; and amino acid residues were represented as one interaction site at the Cα position with corresponding net charge. To accelerate the simulations we truncated the large subunit of the CG ribosome model by retaining those interaction sites within 30 Å of the center line of the exit tunnel, the interaction sites within 20 Å of the peptidyl transferase center (PTC, selected as A2602 in the 23S rRNA of the *E. coli* ribosome) and the interaction sites at the ribosome surface near the tunnel exit, resulting in 4,577 interaction sites in the cropped CG *E. coli* ribosome. The location and shape of the exit tunnel were detected by MOLE*online*[20, 21].

#### 6. CG force field for ribosome-nascent-chain complex (RNC)

The CG ribosome model was held fixed during the simulations except the ribosomal protein L24, which has a long loop that overhangs the exit tunnel opening. We therefore used the CG protein forcefield shown in Supplementary equation (1) to parameterize the *E. coli* ribosomal protein L24 and made residues 42-59 flexible. The fixed part of the CG ribosome model has no intra-ribosome interactions but has interactions with the nascent chain and the flexible part of the ribosome. The potential energy of the CG RNC model is thus described as

$$\begin{aligned} E_{\text{tot}} &= E_{\text{tot}}^{\text{flex-flex}} + E_{\text{tot}}^{\text{flex-fixed}} + E_{\text{tot}}^{\text{R-NC}} + E_{\text{tot}}^{\text{NC-NC}} \\ &= E_{\text{bonding}}^{\text{R-NC}} + \sum_{i\in\{\text{flex} \cup \text{flex}|\text{fixed} \cup \text{NC}\}} K_{\text{b}}(b_i - b_0)^2 \\ &\quad + \sum_{i\in\{\text{flex} \cup \text{flex}|\text{fixed} \cup \text{NC}\}} -\frac{1}{\gamma}\ln\left\{e^{-\gamma\left[K_\alpha(\theta_i-\theta_\alpha)^2+\varepsilon_\alpha\right]} + e^{-\gamma K_\beta(\theta_i-\theta_\beta)^2}\right\} \\ &\quad + \sum_{i\in\{\text{flex} \cup \text{flex}|\text{fixed} \cup \text{NC}\}} \sum_{j=1}^{4} K_{\text{D}_j}\left[1+\cos(j\varphi_i - \delta_j)\right] \\ &\quad + \sum_{\substack{i\in\{\text{R} \cup \text{NC}\},\\ j\in\{\text{NC}\}, i\neq j}} \left\{\frac{q_i q_j}{4\pi\varepsilon_0\varepsilon_\gamma r_{ij}} \cdot e^{-\frac{r_{ij}}{l_\text{D}}} + \varepsilon_{ij}\left[13\left(\frac{R_{ij}}{r_{ij}}\right)^{12} - 18\left(\frac{R_{ij}}{r_{ij}}\right)^{10} + 4\left(\frac{R_{ij}}{r_{ij}}\right)^{6}\right]\right\}, \quad (7) \end{aligned}$$

where $E_{\text{tot}}^{\text{flex-flex}}$ represents the intra-ribosome interactions of the flexible part (ribosomal protein L24 of the *E. coli* ribosome) using Supplementary equation (1); $E_{\text{tot}}^{\text{NC-NC}}$ represents the intra-NC interactions using Supplementary equation (1); $E_{\text{tot}}^{\text{flex-fixed}}$ represents the intra-ribosome interactions between the flexible part and the fixed part; and $E_{\text{R-NC}}$ represents the interactions between NC and the ribosome. Parameters of both $E_{\text{tot}}^{\text{flex-flex}}$ and $E_{\text{tot}}^{\text{NC-NC}}$were obtained from the corresponding native structures using the method described in Supplementary Methods Sections 1 to 3. All the nonbonded interactions in $E_{\text{tot}}^{\text{flex-fixed}}$ and $E_{\text{tot}}^{\text{R-NC}}$ were treated as non-native contacts and calculated in the same manner described in Supplementary Methods Section 1 using the parameters shown in Supplementary Table 6. The parameter $R_i$ of the ribosomal protein interaction sites were derived from the collision diameters $\sigma_i$ used in our previous work[1], while $R_i$ of the RNA interaction sites were calculated from the collision diameters as $\sigma_i = 2 \cdot \langle \bar{d}_{ij}(\text{R-NC}) - \frac{1}{2}\sigma_j \rangle$. The term $\bar{d}_{ij}(\text{R-NC})$ is the average minimum distance between the interaction site type *i* observed in the ribosome and interaction site type *j* observed in NC, whose collision diameter is $\sigma_j$, and was measured by using 6 high-resolution cryo-EM structures of RNCs (PDB 3jbu [22], 4uy8 [23], 5jte [19], 5ju8 [19], 5nwy [24] and 6i0y [25]). $E_{\text{tot}}^{\text{flex-fixed}}$ has extra bonding interactions (including bond, angle and dihedral terms, same as those in Supplementary equation (1)) for the linking bonds at flexible-fixed interface (flex|fixed), i.e., the bond between residue 41 and 42 and the bond between residue 59 and 60 of the ribosomal protein L24 of the *E. coli* ribosome. $E_{\text{tot}}^{\text{R-NC}}$ also has bonded interactions ($E_{\text{bonding}}^{\text{R-NC}}$) for the linking bonds at the NC-tRNA interface, where the force field varies at different steps within the continuous synthesis process, and is described in Supplementary Methods Section 7.

### 7. Protein synthesis simulation protocol

The on-ribosome protein synthesis process was simplified as containing three sub-steps at each time the nascent chain elongates, as shown in Supplementary Fig. 1: (1) the cognate aminoacyl tRNA (aa-tRNA) binds at the A-site; (2) the *l*-length nascent-chain-tRNA complex ($\text{NC}_l$-tRNA) at the P-site and the aa-tRNA at the A-site undergo the peptidyl transfer reaction catalyzed by the PTC and then forms $\text{NC}_{l+1}$-tRNA at the A-site; (3) the P-site tRNA is translocated to the E-site and the A-site tRNA is translocated to the P-site, resulting in an empty A-site for the next round of aa-tRNA binding. We denote the processes from step (1) to step (2), step (2) to step (3) and step (3) to next round step (1) as PT, TL and AB, respectively.

The aa-tRNA molecule was modeled by adding an extra harmonic bond between the amino acid interaction site (CA) and the last nucleotide ribose interaction site (tRNA:76@R, where ':' refers to residues and '@' refers to atoms; similarly hereafter), which introduces extra angle and dihedral energy terms into the aa-tRNA bonding potential energy $E_{\text{aa-tRNA}}^{\text{bonding}}$, resulting in

$$E_{\text{aa-tRNA}}^{\text{bonding}} = E_{\text{bond}} + E_{\text{angle}} + E_{\text{dihedral}}$$
$$= K_{\text{b}}(b - b_0)^2 + \sum_i K_{\theta}(\theta_i - \theta_0)^2 + K_{\varphi}(\varphi - \varphi_0)^2, \quad (8)$$

where $K_{\text{b}}$ is the bond force constant and set as 200 kcal/mol/Å$^2$; $b$ is the pseudo bond length between CA and tRNA:76@R; $b_0$ is the corresponding equilibrium pseudo bond length; $K_{\theta}$ is the angle force constant and set as 25 kcal/mol/rad$^2$; $\theta_i$ includes the angle of CA–tRNA:76@R–tRNA:76@PU2 and the angle of CA–tRNA:76@R–tRNA:76@P; $\theta_0$ is the corresponding equilibrium angle; $K_{\varphi}$ is the dihedral force constant and set as 25 kcal/mol/rad$^2$; $\varphi$ is the dihedral angle of plane CA–tRNA:76@R–tRNA:76@P and plane tRNA:76@R–tRNA:76@P–tRNA:76@PU2; $\varphi_0$ is the corresponding equilibrium dihedral angle.

The NC$_l$-tRNA molecule was modeled by adding an extra harmonic bond between the last amino acid interaction site (NC:*l*@CA) and tRNA:76@R, which introduces the same energy terms with $E_{\text{aa-tRNA}}^{\text{bonding}}$ when *l* = 1. When *l* > 1, this bond introduces an extra double-well angle energy term into the $E_{\text{NC}_l\text{-tRNA}}^{\text{bonding}}$, resulting

$$E_{\text{NC}_l\text{-tRNA}}^{\text{bonding}} = E_{\text{aa-tRNA}}^{\text{bonding}} - \frac{1}{\gamma}\ln\left\{e^{-\gamma\left[K_{\alpha}(\theta-\theta_{\alpha})^2+\varepsilon_{\alpha}\right]} + e^{-\gamma K_{\beta}(\theta-\theta_{\beta})^2}\right\}, \quad (9)$$

where $\theta$ is the angle of NC:(*l*-1)@CA–NC:*l*@CA–tRNA:76@R and the parameters were set as those listed in Supplementary Table 1. All the equilibrium values for both P-site and A-site NC-tRNA (or aa-tRNA) presented in Supplementary equation (8) were set as the average values observed in high-resolution experimental structures: 3jbu [22], 4uy8 [23], 4v5c [26], 5jte [19], 5ju8 [19], 5nwy [24], 6enj [27], 6enu [27] and 6i0y [25]; and are shown in Supplementary Table 7.

The bonding interaction energy term $E_{\text{bonding}}^{\text{R-NC}}$ in Supplementary equation (7) during synthesizing at NC length *l* therefore can be calculated as

$$E_{\text{bonding}}^{\text{R-NC}} = \begin{cases} E_{\text{NC}_{l-1}\text{-tRNA}}^{\text{bonding}}(\text{P-site}) + E_{\text{aa-tRNA}}^{\text{bonding}}(\text{A-site}), & \text{within the process PT} \\ E_{\text{NC}_l\text{-tRNA}}^{\text{bonding}}(\text{A-site}), & \text{within the process TL} \\ E_{\text{NC}_l\text{-tRNA}}^{\text{bonding}}(\text{P-site}), & \text{within the process AB} \end{cases}, \quad (10)$$

where (P-site) means to use the equilibrium bond lengths, angles and dihedrals obtained from P-site tRNA; and (A-site) means to use the equilibrium values obtained from A-site tRNA (see

Supplementary Table 7). Note that $E^{\text{bonding}}_{\text{NC}_{l-1}\text{-tRNA}}(\text{P-site}) = 0$ when $l$ = 1. There are no constraints added on the bond lengths in $E^{\text{R-NC}}_{\text{bonding}}$.

To setup a continuous synthesis simulation, we used the corresponding mRNA sequence of the protein to be synthesized, the truncated CG ribosome structure of the corresponding organism and the CG RNC force field with optimized $n_{\text{scal}}$ for this protein. The synthesis begins at the start codon and ends at the stop codon. At each codon position, we obtained the initial RNC structure from the final structure calculated at the previous codon position, followed by putting the new amino acid interaction site near the A-site tRNA:76@R with an initial orientation based on the experimental data listed in Supplementary Table 7. At each sub-step, an energy minimization was performed on the last 15 residues of the C-terminal of NC, as well as the new amino acid at step 1, followed by running a MD simulation for a time $\tau^{\text{sim}}$. At step 1, we connected the bond between the new amino acid and the A-site tRNA. At step 2, we connect the ($l$-1)$^{\text{th}}$ amino acid interaction site to the new amino acid at A-site tRNA and break the original bond between the ($l$-1)$^{\text{th}}$ amino acid interaction site and the P-site tRNA. At step 3, the $l$-length NC was covalently connected to the P-site tRNA and the empty A-site was modeled by excluding all the nonbonded interactions of A-site tRNA with other interaction sites. The simulation time $\tau^{\text{sim}}$ at each step of each codon position was randomly sampled from the exponential distribution with the expectation $\langle\tau^{\text{sim}}\rangle$. Details on derivation of $\langle\tau^{\text{sim}}\rangle$ can be found in the Supplementary Methods Section 8.

Once the nascent chain was fully synthesized, the bond between NC:$l$@CA and tRNA:76@R was removed, as well as the corresponding bonding interaction energy terms. The nascent chain was ejected from the exit tunnel and then dissociated from the ribosome surface. We identified the complete dissociation as when the minimum distance between the nascent chain and the ribosome surface is greater than 20 Å for at least 750 ps. The final nascent chain structure was then used to study the off-ribosome post-translational folding.

### 8. Setting realistic *in silico* codon translation times

Even though the folding process is accelerated in the coarse-grained model, we maintain a realistic ratio of folding and translation times. To set a realistic *in silico* mean translation time $\langle\tau^{\text{sim}}_{\text{trans}}(S)\rangle_i$ for codon $i$ from the corresponding real mean translation time $\langle\tau^{\text{real}}_{\text{trans}}(S)\rangle_i$ given the mRNA sequence $S$ and the organism, we use the scaling factor $\alpha$ from *in silico* protein folding times (see Supplementary Methods Section 4 and Supplementary Table 4) as $\langle\tau^{\text{sim}}_{\text{trans}}(S)\rangle_i =$

$\frac{1}{\alpha}\langle\tau_{\text{trans}}^{\text{real}}(S)\rangle_i$. This takes into account of the influence of ribosome traffic $\langle\tau_{\text{trans}}^{\text{ribo-traffic}}(S)\rangle_i$ on the intrinsic translation time $\langle\tau_{\text{trans}}^{\text{intrinsic}}\rangle_i$ and is expressed as

$$\begin{aligned}\langle\tau_{\text{trans}}^{\text{real}}(S)\rangle_i &= \langle\tau_{\text{trans}}^{\text{intrinsic}}\rangle_i + \langle\tau_{\text{trans}}^{\text{ribo-traffic}}(S)\rangle_i \\ &= \langle\tau_{\text{AB}}^{\text{intrinsic}}\rangle_i + \langle\tau_{\text{PT}}^{\text{intrinsic}}\rangle + \langle\tau_{\text{TL}}^{\text{intrinsic}}\rangle + \langle\tau_{\text{trans}}^{\text{ribo-traffic}}(S)\rangle_i \\ &= \langle\tau_{\text{AB}}^{\text{intrinsic}}\rangle_i + \langle\tau_{\text{PT}}^{\text{intrinsic}}\rangle + \langle\tau_{\text{TL}}^{\text{real}}(S)\rangle_i, \qquad (11)\end{aligned}$$

where $\langle\tau_{\text{AB}}^{\text{intrinsic}}\rangle_i$ is the intrinsic mean dwell time of the A-site aa-tRNA binding at codon *i* taking into account the competition of the non- and near-cognate tRNAs binding; $\langle\tau_{\text{PT}}^{\text{intrinsic}}\rangle$ is the intrinsic mean dwell time of the peptidyl transfer reaction at the PTC after A-site aa-tRNA binding and is independent of the codon identity; $\langle\tau_{\text{TL}}^{\text{real}}(S)\rangle_i$ is the real mean dwell time of tRNA translocation after the peptidyl transfer reaction and can be calculated as the summation of the intrinsic mean dwell time of translocation $\langle\tau_{\text{TL}}^{\text{intrinsic}}\rangle$ and the mean delay time due to the ribosome traffic $\langle\tau_{\text{trans}}^{\text{ribo-traffic}}(S)\rangle_i$ at codon *i*. $\langle\tau_{\text{trans}}^{\text{real}}(S)\rangle_i$ was estimated by a kinetic model proposed in our previous study[28] using the particular mRNA sequence *S*, the translation initiation rate and the intrinsic codon translation time of the particular organism. The translation initiation rate was set as the median value observed in experiments and simulations, which is 0.083 $s^{-1}$ for *E. coli*[29].

The intrinsic codon translation times $\langle\tau_{\text{trans}}^{\text{intrinsic}}\rangle_i$ of *E. coli*, as well as the intrinsic dwell times of sub-steps ($\langle\tau_{\text{PT}}^{\text{intrinsic}}\rangle$ and $\langle\tau_{\text{TL}}^{\text{intrinsic}}\rangle$), were obtained from Fluitt & Viljoen's work[30] (denoted as $\langle\tau_{\text{Fluitt}}^{\text{intrinsic}}\rangle_i$), followed by rescaling the data so that the average codon translation time equals the experimental average $\overline{\tau_{\text{trans}}^{\text{exp}}}$:

$$\langle\tau_{\text{trans}}^{\text{intrinsic}}\rangle_i = \frac{\overline{\tau_{\text{trans}}^{\text{exp}}}}{\sum_{k=1}^{64} w_k \cdot \langle\tau_{\text{Fluitt}}^{\text{intrinsic}}\rangle_k} \cdot \langle\tau_{\text{Fluitt}}^{\text{intrinsic}}\rangle_i, \qquad (12)$$

where $w_k$ is the normalized codon usage frequency[31] of the $k^{\text{th}}$ codon in *E. coli*. The experimental codon translation rate varies from 12 to 21 $s^{-1}$ in *E. coli* at 310 K[32], resulting in an average rate of 16.5 $s^{-1}$. Thus, $\overline{\tau_{\text{trans}}^{\text{exp}}}$ was set as 0.061 s. All of the values for $\langle\tau_{\text{trans}}^{\text{intrinsic}}\rangle_i$, $\langle\tau_{\text{PT}}^{\text{intrinsic}}\rangle$, $\langle\tau_{\text{TL}}^{\text{intrinsic}}\rangle$ and $w_k$ can be found in Supplementary Table 8.

From Supplementary equation (11), the *in silico* mean dwell times of the sub-steps during translating codon *i* were then calculated as

$$\begin{cases} \langle \tau_{\mathrm{AB}}^{\mathrm{sim}} \rangle_i = \dfrac{1}{\alpha} \langle \tau_{\mathrm{AB}}^{\mathrm{intrinsic}} \rangle_i \\ \langle \tau_{\mathrm{PT}}^{\mathrm{sim}} \rangle = \dfrac{1}{\alpha} \langle \tau_{\mathrm{PT}}^{\mathrm{intrinsic}} \rangle \\ \langle \tau_{\mathrm{TL}}^{\mathrm{sim}}(S) \rangle_i = \dfrac{1}{\alpha} \langle \tau_{\mathrm{TL}}^{\mathrm{real}}(S) \rangle_i = \dfrac{1}{\alpha} \left[ \langle \tau_{\mathrm{TL}}^{\mathrm{intrinsic}} \rangle + \langle \tau_{\mathrm{trans}}^{\mathrm{ribo\text{-}traffic}}(S) \rangle_i \right] \\ \qquad = \dfrac{1}{\alpha} \left[ \langle \tau_{\mathrm{TL}}^{\mathrm{intrinsic}} \rangle + \langle \tau_{\mathrm{trans}}^{\mathrm{real}}(S) \rangle_i - \langle \tau_{\mathrm{trans}}^{\mathrm{intrinsic}} \rangle_i \right] \end{cases}, \qquad (13)$$

### 9. Virtual screening for proteins that get kinetically trapped during folding

To identify candidates whose catalytic property might be affected by synonymous mutations, we identified an initial set of enzymes by searching the databases EzCatDB[33, 34], UniProt[35] and PDB[36]. This initial set was of enzymes that (1) are endogenetic enzymes from *E. coli*; (2) have a known catalytic mechanism; (3) can be active as a monomer; (4) are located in the cytoplasm and not transmembrane; (5) have resolved crystal or NMR structures with bound substrates; and (6) do not have missing domains in their resolved structures (having several missing loops or fragments is acceptable). The catalytic mechanism information was obtained from EzCatDB; the enzymology information was obtained from UniProt; and the structural information was obtained from PDB. We found 14 *E. coli* enzymes that met these criteria. We chose the resolved structure that has the highest resolution and has the most relevant substrate binding, for those enzymes that have more than one resolved structure. The structural domains of those initial enzymes were then identified by using the CATH database and parameterized using our CG protein model (see Supplementary Methods Section 1) via the stepwise parameterization strategy (see Supplementary Methods Section 3). All the structural information and the parameterization information for these enzymes can be found in Supplementary Table 9. The wild-type enzymes were then continuously synthesized on the corresponding ribosome, followed by post-translational folding simulations (both have 10 replicas). The post-translational folding simulations for all the initial enzymes were stopped at a 14-day wall time. The candidates were then selected by using the score function equation (1) in the main text. The first term in the scoring function adopts the form $1 - \left(1 + \langle \tau_{\mathrm{F}}^{\mathrm{post}} \rangle\right)^{-k}$, where *k* is a positive parameter controlling the slop. In order to distinguish both large (~$10^{10}$) and small (~$10^{-2}$) $\langle \tau_{\mathrm{F}}^{\mathrm{post}} \rangle$ values, we constrain $1 - (1 + 10^{10})^{-k} < 0.9999$ and $1 - (1 + 10^{-2})^{-k} > 0.0001$, which result in $0.01 < k < 0.4$. We thus set *k* as the average $k = 0.2$. The time point when the protein reaches the folded state was used to estimate $\langle \tau_{\mathrm{F}}^{\mathrm{post}} \rangle$ in the score function and was identified as the time when $Q^{\mathrm{mod}}$ is higher than $\langle Q_{\mathrm{native}}^{\mathrm{mod}} \rangle - 3\sigma$ for 7.5 ns in simulation timescale. $Q^{\mathrm{mod}}$ is the mode of Q-values calculated by histogramming a moving window of 200 data points along with the post-translational time course using a bin width 0.02; $\langle Q_{\text{native}}^{\text{mod}} \rangle$ is the average of mode-filtered $Q$ values of the native state calculated by using the 10 trajectories obtained from the final iteration of the stepwise parameterization strategy; and $\sigma$ is the standard deviation. $Q_{\text{act}}^{\text{mod}}$ (the mode of $Q_{\text{act}}$, see main text equation (3)) is used to identify whether or not the misfolding occurs at the substrate binding site. $\Theta_{\text{binding}} = 1$ when at least one trajectory has $Q_{\text{act}}^{\text{mod}} < \langle Q_{\text{act, native}}^{\text{mod}} \rangle - 3\sigma$ at the last 7.5 ns; otherwise $\Theta_{\text{binding}} = 0$.

**10. Parameterization of the CG models for CAT-III**

CAT-III monomer is a single domain protein and was parameterized using the same approach as the training-set proteins (Supplementary Methods Section 2). The folding stability, -10.1 kcal/mol, was estimated via PREFUR algorithm[37] at 298 K. The native structure was obtained from PDB ID 3cla[38] and chain A was selected to create the CG model. Three $n_{\text{scal}}$ values (1.0, 1.1 and 1.2) were utilized to search for the optimized $n_{\text{scal}}^{*}$ value. At each $n_{\text{scal}}$, two pt-REMD simulations were initialized from native state and unfolded state, respectively. The parameterization results for CAT-III are shown in Supplementary Table 10.

**11. Detecting noncovalent lasso entanglements in protein structures**

To detect noncovalent lasso entanglements we use linking numbers[39], which requires at least one closed loop as an argument. We define this loop as being composed of the backbone trace connecting residues *i* and *j*, which have formed a native contact in the given protein conformation. The native contact between *i* and *j* is considered to close this loop, even though there is no covalent bond between these two residues. Outside this loop is an N-terminal segment, composed of residues 1 through *i*-1, and C-terminal segment composed of residues *j*+1 through *N*. These two segments represent open curves, whose entanglement through the closed loop we characterize with linking numbers denoted $g_{\text{N}}$ and $g_{\text{C}}$. We calculate these numbers using the partial Gauss double integration method proposed by Baiesi and co-workers[40]. For a given structure of an *N*-length protein, with a native contact present at residues (*i*, *j*), the coordinates $\boldsymbol{R}_l$ and the gradient $\mathrm{d}\boldsymbol{R}_l$ of the point *l* on the curves were calculated as

$$\begin{cases} \boldsymbol{R}_l = \dfrac{1}{2}(\boldsymbol{r}_l + \boldsymbol{r}_{l+1}) \\ \mathrm{d}\boldsymbol{R}_l = \boldsymbol{r}_{l+1} - \boldsymbol{r}_l \end{cases}, \tag{14}$$

where $\boldsymbol{r}_l$ is the coordinates of the Cα atom in residue *l*. The linking numbers $g_{\text{N}}(i,j)$ and $g_{\text{C}}(i,j)$ were calculated as

$$
\begin{cases}
g_{\mathrm{N}}(i,j) = \dfrac{1}{4\pi}\displaystyle\sum_{m=6}^{i-5}\sum_{n=i}^{j-1}\dfrac{\boldsymbol{R}_m - \boldsymbol{R}_n}{|\boldsymbol{R}_m - \boldsymbol{R}_n|^3}\cdot(\mathrm{d}\boldsymbol{R}_m \times \mathrm{d}\boldsymbol{R}_n) \\
g_{\mathrm{C}}(i,j) = \dfrac{1}{4\pi}\displaystyle\sum_{m=i}^{j-1}\sum_{n=j+4}^{N-6}\dfrac{\boldsymbol{R}_m - \boldsymbol{R}_n}{|\boldsymbol{R}_m - \boldsymbol{R}_n|^3}\cdot(\mathrm{d}\boldsymbol{R}_m \times \mathrm{d}\boldsymbol{R}_n)
\end{cases}, \tag{15}
$$

where we excluded the first 5 residues on the N-terminal curve, last 5 residues on the C-terminal curve and 4 residues before and after the native contact for the purpose of eliminating the error induced by both the high flexibility and contiguity of those tails. The above integrations yield two non-integer values, therefore the total linking number for a native contact ($i$, $j$) was estimated as

$$
g(i,j) = \mathrm{round}\big(g_{\mathrm{N}}(i,j)\big) + \mathrm{round}\big(g_{\mathrm{C}}(i,j)\big). \tag{16}
$$

As an illustration and test case we analyzed the post-translational trajectories of fast CAT-III variant. We calculated the maximum $g(i,j)$ for every frame of the trajectories and found that the maximum $g(i,j)$ varies from -2 to 2. We randomly chose structures corresponding to each maximum $g(i,j)$ detected and compared the structure with the corresponding linking diagram shown in Supplementary Fig. 2. We clearly see entanglements in these structures. As shown in Supplementary Fig. 2 b-f, all the maximum $g(i,j)$ values estimated from Supplementary equation (16) are consistent with those calculated as the total number of all the +1 and -1 crossings in the link diagram (left side in each panel) divided by 2.

### 12. Assessing the influence of synonymous mutations on co- and post-translational folding

The translation schedules for the fastest (denoted as fast variant) and the slowest synonymous mRNA variant (denoted as slow variant) were predicted according to Fluitt & Viljoen's codon translation time in Supplementary Table 8. We performed continuous synthesis simulations for the fast and slow variants using 50 trajectories for DHFR and 100 trajectories for CAT-III and DDLB, followed by post-translational folding simulations in the bulk environment for 10 seconds for DHRF (in which all trajectories are folded) and 60 seconds for CAT-III and DDLB on the experimental timescale (2.31 μs and 13.85 μs of simulation time, respectively). The number of post-translational trajectories was increased to 10 times larger than the continuous synthesis simulations (500 trajectories for DHFR and 1,000 trajectories for CAT-III and DDLB) for each variant.

The structural distribution from the post-translational simulations was assessed as -ln$P$, ($P$ is the probability of sampling various configurations) along the order parameters $G$ and $Q_{\mathrm{act}}$ as described in the main text (equations (2) and (3)). To further analyze the post-translational kinetics, 400 clusters (micro-states) were grouped from the combined set of post-translational trajectories of both synonymous variants using the $k$-means algorithm[41, 42]. A Markov state model (MSM) was built and the clusters were coarsen-grained into a small number of metastable states using the PCCA+ algorithm[43]. The number of metastable states was chosen based on the existence of a gap in the eigenvalue spectrum of the transition probability matrix[44]. Five representative structures of each metastable state were randomly sampled from all microstates according to the probability distribution of the microstates within the given metastable state. The time evolution of the probability distribution of metastable states (see Supplementary Methods Section 14.1), as well as the representative structures, were used to estimate the specific activity of a synonymous variant. All the clustering and MSM building were performed by using PyEmma package[45].

#### 13. Back-mapping the CG protein model to an all-atom (AA) model

To be able to study enzymatic catalysis at the AA resolution, a customized back-mapping procedure was utilized to rebuild AA models for those CG protein structures. The CG interaction sites that represent the side-chain center-of-mass were first built near the corresponding Cα interaction sites, whose orientation was further refined by using energy minimizations under the Cα-SCM force field[46] with all Cα positions restrained. The backbone atoms were then rebuilt by Prodart2[47] and the side-chain atoms were rebuilt by Pulchra[48]. The final AA structure was obtained after a further energy minimization in vacuum with all Cα positions restrained.

#### 14. Calculating the specific activity from the structural distribution of enzyme conformations

##### 14.1. Estimation of specific activity for an enzyme

The specific activity (SA) for a given protein variant is estimated by utilizing the distribution of metastable states (whose state probability is $p_i$ for metastable state *i*) and the activation free energy barrier height ($\Delta G_i^{\ddagger}$) of each state as described in the main text (equation (4)), as well as the relative specific activity (SA*, see main text (equation (5)). To reduce the error of estimated reaction rates, $\Delta G_i^{\ddagger}$ was obtained as the median value of the successfully estimated activation barrier heights of the representative structures for each metastable state. To account for only the soluble conformations, the state probability $p_i$ was estimated as $p_i = p_i^0 \cdot f_i^{\text{sol}} / [\sum_i p_i^0 \cdot f_i^{\text{sol}}]$, where $f_i^{\text{sol}}$ is the percent soluble protein of state *i* (see details in Supplementary Methods Section 14.2) and $p_i^0$ is the raw probability of state *i* at the end of post-translational simulations (10 s for DHFR and 60 s for CAT-III and DDLB) that was obtained from discrete metastable state trajectories in Supplementary Methods Section 12.

#### 14.2. The percent of soluble protein from each metastable state

The percent soluble protein $f_i^{\text{sol}}$ was estimated as

$$f_i^{\text{sol}} = \frac{\chi_{\text{disordered}}^{\text{insol}} - \chi_i^{\text{insol}}}{\chi_{\text{disordered}}^{\text{insol}} - \min\left(\chi_i^{\text{insol}}\right)} \times 100\%, \tag{17}$$

where $\chi_i^{\text{insol}}$, $\chi_{\text{disordered}}^{\text{insol}}$ and $\min\left(\chi_i^{\text{insol}}\right)$ are the insolubility propensities of state *i*, the fully disordered structure and the minimum propensity value throughout all states, respectively. The fully disordered protein has no secondary and tertiary structure elements and was built as a linear sequence of amino acid residues. The insolubility propensity $\chi^{\text{insol}}$ was estimated by taking into account of the aggregation propensity $\chi^{\text{agg}}$, degradation propensity $\chi^{\text{deg}}$ and subtracting the Hsp70 binding propensity $\chi^{\text{Hsp70}}$ that is considered to prevent the misfolding protein from aggregation:

$$\chi^{\text{insol}} = \chi^{\text{agg}} + \chi^{\text{deg}} - \chi^{\text{Hsp70}}. \tag{18}$$

For a given metastable state, the aggregation, degradation and Hsp70 binding propensities were evaluated as the relative change of the mean solvent accessible surface area (SASA) of the aggregation-prone, degradation-prone and Hsp70 binding regions of the representative protein structures (or the modeled fully disordered structure) against the mean SASA of the native state, respectively:

$$\chi = \frac{\langle \text{SASA} \rangle}{\langle \text{SASA} \rangle_{\text{native}}} - 1. \tag{19}$$

A negative $\chi$ value indicates a lower propensity than the native state. The aggregation-prone region was predicted by the AmylPred2 server[49]. The degradation-prone region was defined as the hydrophobic residues (Ile, Val, Leu, Phe, Cys, Met, Ala, Gly and Trp). The Hsp70 binding region was predicted by using ChaperISM[50]. Note that, for CAT-III, the trimerization interface area (residue 25 to 33 and 150 to 157) was subtracted from the above SASAs. The 95% CIs of the propensities and the percent soluble protein were estimated using the bootstrapping method with 10,000 iterations.

#### 14.3. Estimating the activation free energy barrier for a protein conformation

We developed the following workflow (illustrated in Supplementary Fig. 3) to estimate activation free energy barrier height for a given enzyme structure:

(1) Take the representative structure (CG protein structure) of a metastable state. Back-map the CG structure to its AA structure.

(2) Refine the rebuilt AA structure by running energy minimizations in vacuum for 1,000 steps

with all Cα positions restrained using a force constant 30 kcal/mol/Å$^2$ under a dielectric constant 78.5 to relax the side chain orientations, followed by running energy minimizations in a water box for 2,000 steps with all Cα positions restrained using a force constant 100 kcal/mol/Å$^2$ to relax water molecules and ions. The final refined AA structure was obtained after running energy minimizations in water for 2,000 steps with all Cα positions restrained using a force constant 10 kcal/mol/Å$^2$.

(3) For an enzyme that can only carry out the reaction in its homomultimer form, an estimation of the multimer structure was performed by using SymmDock program[51]. For example, CAT-III needs to form a trimer to conduct the reaction. The monomer structure of CAT-III obtained from step (2) was first refined to form a native-like trimer interface by running targeted energy minimizations in vacuum for 2,000 steps, where an RMSD restraint on the trimer interface region towards the corresponding native structure (crystal structure) with a force constant 100 kcal/mol/Å$^2$ was applied. The refined monomer structure was then used to predict the trimer structure using SymmDock with distance restraints between monomers at the interface region to force the trimer to be formed on the native interface. If the prediction succeeded, the best candidate was obtained and the trimer interface was refined by running targeted energy minimizations in vacuum for 2,000 steps, where an RMSD restraint on the trimer interface region towards the corresponding native structure with a force constant 10 kcal/mol/Å$^2$ were applied. The final refined trimer structure was obtained after running a series of MD simulations in a water box: First, run energy minimizations for 2,000 steps with all Cα positions restrained using a force constant 10 kcal/mol/Å$^2$; Second, heat the system to 310 K in NVT ensemble with all Cα positions restrained using a force constant 10 kcal/mol/Å$^2$; Third, equilibrate the system at 310 K in the NPT ensemble for 0.5 ns with all Cα positions restrained using a force constant 5 kcal/mol/Å$^2$; Finally, run energy minimizations again for 500 steps with all Cα positions restrained using a force constant 10 kcal/mol/Å$^2$.

(4) After the AA structure of the monomer or multimer was refined, the structure was then used to predict the ligand-binding poses using AutoDock vina[52]. The docking boxes were set as the minimum box that covers each ligand presented in the crystal structure. The best binding pose (highest score) of each ligand was used to build the protein-ligand complex structure. The complex structure was then refined by running the following MD simulations: First, solvate the complex in TIP3P water molecules and neutralization ions (Na+ or Cl-); Second, run energy minimizations for 2,000 steps with all Cα positions restrained using a force constant 100 kcal/mol/Å$^2$, followed by running energy minimizations for 2,000 steps with all Cα positions

restrained using a force constant 5 kcal/mol/Å$^2$; Third, heat the system to 310 K in the NVT ensemble with all Cα positions restrained using a force constant 3 kcal/mol/Å$^2$ for 20 ps; Forth, equilibrate the system in the NPT ensemble with all Cα positions restrained using a force constant 1 kcal/mol/Å$^2$ for 400 ps. A few distance restraints between ligands and surrounding residues were added to force the orientation of ligands to be the most native-like. The distance restraints for refining the binding poses of the reactants were set as the following: If the reactant is co-crystalized in the crystal structure, some of the representative interactions were chosen and the distances were restrained to the exact value; and if the reactant is not presented in any crystal structure, the possible interactions were obtained from the literature or inferred theoretically and the distances were restrained less than an expected value. The distance restraint force constant was stepwise increased to 500 kcal/mol/Å$^2$ in the first 200 ps and kept 500 kcal/mol/Å$^2$ in the later 200 ps; Finally, the refined complex structure was obtained after running a classical energy minimization for 2,000 steps, followed by a QM/MM energy minimization for 5,000 steps with the distance restraints between ligands and catalytic residues. The distance restraint force constant was stepwise increased to 10 kcal/mol/Å$^2$ in the first 2,500 steps and kept 10 kcal/mol/Å$^2$ in the later 2,500 steps.

The activation free energy profile was estimated by running QM/MM umbrella sampling simulations along the predefined reaction coordinates (RCs). In each umbrella window, the simulation was run in the NPT ensemble at 310 K for 20 ps after a 1,000-step energy minimization. The umbrella restraint force constant was set as 250 kcal/mol/Å$^2$. The potential of mean force (PMF) was unbiased and estimated by the WHAM equation[16] using the last 10 ps trajectories (see Supplementary Figures 8, 9, 10). The activation free energy barrier height was obtained as the difference between the maximum PMF value and the minima before the maximum. The 95% CIs were estimated by using the Monte Carlo bootstrap error analysis[53] with 100 trials.

#### 14.4. Classical Molecular Dynamics Simulations

For a vacuum system in the above workflow, the MD simulations were performed without a nonbonded cutoff and using the dielectric constant set as 78.5. For a solvated system, the solute was embedded in a periodic TIP3P water box. Several ions were added to neutralize the system. Particle Mesh Ewald (PME) method[54] was used to calculate the long-range electrostatic interactions with a 10 Å cutoff. The NPT ensemble simulations were performed at 310 K temperature and 1 bar pressure via Langevin dynamics (the collision frequency is 1.0 ps$^{-1}$), with a coupling constant of 0.2 ps for both parameters. The SHAKE algorithm[55] was applied to the bonds involving hydrogen, which ensures the integral timestep to be 2 fs. All the classical MD simulations in the workflow were performed by Amber17[56] with *ff14SB* protein force field[57]. The ligands were parameterized by the general force field *gaff*[58]. The atomic charges of the ligands were estimated as the RESP charges[59] derived from the QM optimization with Hartree–Fock (HF) method at 6-31G* level using Gaussian 09[60].

#### 14.5. QM/MM Simulations

For QM/MM umbrella sampling simulations, the QM region (including ligands and catalytic residues) was simulated using the third-order density functional tight binding (DFTB3) Hamiltonian[61] with 3ob-3-1 parameter set[62, 63, 64, 65]. The MM region was simulated using *ff14SB* protein force field[57]. The QM/MM interface was built by inserting explicit link atoms (hydrogen atoms). The interaction on QM/MM interface was estimated using the electrostatic embedding scheme. Particle Mesh Ewald (PME) method[54] was used to calculate the long-range electrostatic interactions with a 10-Å cutoff. The NPT ensemble simulations were performed at 310 K temperature and 1 bar pressure via the Langevin dynamics (the collision frequency is 1.0 $ps^{-1}$), with a coupling constant of 0.2 ps for both parameters. The SHAKE algorithm[55] was applied to the bonds involving hydrogen in the MM region and no constraints were applied in the QM region to enable the proton transfer. The MD integration timestep was thus set as 1 fs. All the QM/MM umbrella sampling simulations were performed by Amber17[56].

#### 14.6. Detailed setup for each system

***CAT-III.*** The chain A of the crystal structure 3CLA was used as the template for back-mapping. The trimer structure of 3CLA was used as the template for estimating the trimer of representative structures. The trimer interface region in the monomer was identified as residue 25 to 33 and 150 to 157 (the residue index starts from 1 here, whereas it starts from 6 in the PDB file). The distance restraints applied during the prediction of trimer structure are on the distance between residue 25 in two monomers, distance between residue 32 in two monomers, distance between residue 150 in two monomers, distance between residue 157 in two monomers, distance between residue 25 in one monomer and 157 in another, and distance between residue 32 in one monomer and 150 in another. The binding poses of the reactant chloramphenicol (CLM) were searched within the box centered at (-1.000, 14.600, 11.200) Å with the dimension 14 Å × 14 Å × 14 Å. The binding poses of the reactant acetyl coenzyme A (ACO) were searched within the box centered at (-2.062, 16.000, 30.000) Å with the dimension 15 Å × 15 Å × 30 Å. The following distance restraints were used to refine the binding poses with classical MD simulations (residue indices start from 1 and are accumulated throughout the trimer. The Amber-style masking syntax is used to denote the selected atoms, where ':' denotes residuals and '@' denotes atoms, and the unit of distance is in

Å, similarly hereinafter): *d*(:402@NE2, :CLM@O1) = 2.830; *d*(:402@NE2, :CLM@N2) = 8.750; *d*(:402@NE2, :CLM@C6) = 5.450; *d*(:402@NE2, :CLM@C2) = 4.020; *d*(:402@NE2, :CLM@C4) = 7.270; *d*(:402@NE2, :CLM@O5) = 5.650; *d*(:402@CG, :CLM@O1) = 4.940; *d*(:402@CG, :CLM@N2) = 9.180; *d*(:402@CG, :CLM@C6) = 6.680; *d*(:402@CG, :CLM@C2) = 5.800; *d*(:402@CG, :CLM@C4) = 8.720; *d*(:402@CG, :CLM@O5) = 7.400; *d*(:402@ND1, :CLM@O1) = 4.890; *d*(:402@ND1, :CLM@N2) = 10.090; *d*(:402@ND1, :CLM@C6) = 7.290; *d*(:402@ND1, :CLM@C2) = 5.970; *d*(:402@ND1, :CLM@C4) = 8.790; *d*(:402@ND1, :CLM@O5) = 7.750; *d*(:CLM@O1, :ACO@C22) < 3.000; *d*(:402@NE2, :ACO@S1) < 3.000; *d*(:402@ND1, :406@OD1) = 3.380. The following distance restraints were used to refine the active site with QM/MM energy minimization: *d*(:402@NE2, :CLM@O1) < 2.830; *d*(:CLM@O1, :ACO@C22) < 3.000; *d*(:402@NE2, :ACO@S1) < 3.000; *d*(:402@ND1, :406@OD1) < 3.000. The QM region (see Supplementary Fig. 4a) includes reactant CLM, a part of ACO with the acetyl group (from C4 to C22), the side chain of His402[38, 66] and the side chain of Asp406[66]. The reaction coordinate (RC) was defined as $\mathrm{RC} = d(\text{:ACO@C22, :ACO@S1}) - d(\text{:CLM@O1, :ACO@C22})$ and discretized into 34 windows: RC = -2.6, -2.4, -2.2, -2.0, -1.8, -1.6, -1.4, -1.2, -1.0, -0.9, -0.8, -0.7, -0.6, -0.5, -0.4, -0.3, -0.2, -0.1, 0.0, 0.1, 0.2, 0.3, 0.4, 0.5, 0.6, 0.7, 0.8, 0.9, 1.0, 1.2, 1.4, 1.6, 1.8, 2.0. The metastable states P9, P10, P11, P12, P13 and P14 were taken to evaluate $\Delta G_i^{\ddagger}$.

***DDLB.*** The chain A of the crystal structure 4C5C was used as the template for back-mapping. Two $Mg^{2+}$ ions (denoted as MC) that facilitate the proper orientation of ATP were docked at position (5.523, 31.727, 65.029) Å and (7.170, 33.772, 62.509) Å, respectively. The $Mg^{2+}$ ions were parameterized by the dummy-atom model[67], which has an enhanced performance in modeling metalloenzymes. The binding poses of reactant ATP were searched within the box centered at (7.538, 30.141, 68.311) Å with the dimension 7 Å × 11 Å × 16 Å. The binding poses of reactant zwitterionic D-alanine ($-NH_3^+$ and $-COO^-$, DAL) were searched within the box centered at (4.876, 31.014, 58.470) Å with the dimension 4 Å × 4 Å × 4 Å. The binding poses of reactant anionic D-alanine ($-COO^-$, DAN) were searched within the box centered at (2.031, 29.261, 60.338) Å with the dimension 4 Å × 4 Å × 4 Å. The following distance restraints were used to refine the binding poses with classical MD simulations: *d*(:272@OD1, :307@MC) = 2.16; *d*(:270@OE1, :307@MC) = 2.20; *d*(:270@OE2, :307@MC) = 2.13; *d*(:270@OE2, :308@MC) = 2.13; *d*(:257@OD2, :308@MC) = 2.05; *d*(:307@MC, :308@MC) = 3.64; *d*(:210@CA, :ATP@O2) = 3.68; *d*(:187@OE1, :ATP@O2) = 2.66; *d*(:187@OE2, :ATP@O1) = 2.43; *d*(:181@C, :ATP@N3) = 4.03; *d*(:144@NZ, :ATP@N2) = 3.00; *d*(:180@OE1, :ATP@N2) = 3.37; *d*(:180@C, :ATP@N3) = 4.67; *d*(:215@NZ, :ATP@O11) = 2.77; *d*(:ATP@O11, :308@MC) = 2.19;

*d*(:ATP@O10, :151@CA) = 3.26; *d*(:ATP@O9, :97@NZ) = 2.74; *d*(:ATP@O13, :308@MC) = 2.03; *d*(:ATP@O7, :308@MC) = 1.97; *d*(:ATP@O12, :307@MC) = 1.96; *d*(:ATP@O10, :307@MC) = 2.02; *d*(:255@CZ, :DAL@O1) = 3.73; *d*(:15@OE2, :DAL@N1) = 2.95; *d*(:281@OG, :DAN@O1) = 2.61; *d*(:DAL@C1, :DAN@N1) < 3.50; *d*(:DAL@O2, :ATP@P3) < 3.00; *d*(:DAL@N1, :ATP@O12) = 2.80; *d*(:DAL@C1, :307@MC) = 5.66. The following distance restraints were used to refine the active site with QM/MM energy minimization: *d*(:DAL@C1 :DAN@N1) < 3.00; *d*(:DAL@O2, :ATP@P3) < 3.50; *d*(:255@CZ, :DAL@O1) < 3.00; *d*(:281@OG, :DAN@O1) < 3.00; *d*(:15@OE2, :DAL@N1) < 3.00, as well as all the P-O bonds in the ATP molecule. The QM region (see Supplementary Fig. 4b) includes reactant DAL, DAN, a part of ATP with all the phosphate groups (from C1 to P3) and the side chain of Arg255 (play an important role in stabilizing transition states and intermediates[68]). The reaction coordinate (RC) was defined as $\mathrm{RC} = 0.6d(\text{:ATP@O11}, \text{:ATP@P3}) - 0.6d(\text{:ATP@P3}, \text{:DAL@O2}) + 0.4d(\text{:DAL@C1}, \text{:DAL@O2}) - 0.4d(\text{:DAN@N1}, \text{:DAL@C1})$ and discretized into 33 windows: RC = -1.7, -1.5, -1.4, -1.3, -1.2, -1.1, -1.0, -0.9, -0.8, -0.7, -0.6, -0.5, -0.4, -0.3, -0.2, -0.1, 0.0, 0.2, 0.4, 0.6, 0.8, 0.9, 1.0, 1.1, 1.2, 1.3, 1.4, 1.5, 1.6, 1.7, 1.8, 1.9, 2.1. To ensure the reaction occurs within the RCs, a few extra distance restraints were applied in the umbrella sampling simulations: *d*(:ATP@O11, :ATP@P3) < 4.00; *d*(:ATP@O6, :DAL@C1) > 2.00; *d*(:ATP@O12, :DAL@C1) > 2.00; *d*(:ATP@O13, :DAL@C1) > 2.00; *d*(:DAL@C1, :DAL@O2) < 3.00, as well as all the P-O bonds in the ATP molecule except for the P3-O11 bond. The metastable states P4, P5, P6, P7, P8, P9 and P10 were taken to evaluate $\Delta G_i^{\ddagger}$.

***DHFR.*** The crystal structure 4KJK was used as the template for back-mapping. The reaction mechanism of DHFR includes a proton transfer step, followed by a hydride transfer[69]. The hydride transfer from NADPH to the protonated dihydrofolate (DHF-H+) is the rate-limiting step[69]. We therefore only studied the hydride transfer step. The binding poses of reactant NADPH (NPH) were searched within the box centered at (4.333, 6.861, -16.028) Å with the dimension 40 Å × 54 Å × 20 Å. The binding poses of reactant DHF-H+ (DFH) were searched within the box centered at (0.861, -3.250, -6.861) Å with the dimension 20 Å × 28 Å × 32 Å. The following distance restraints were used to refine the binding poses with classical MD simulations: *d*(:78@CA, :NPH@N5) = 4.93; *d*(:44@CZ, :NPH@P3) = 4.09; *d*(:98@CZ, :NPH@O13) = 3.18; *d*(:46@OG1, :NPH@O10) = 2.70; *d*(:49@OG, :NPH@O2) = 3.71; *d*(:7@CA, :NPH@N1) = 4.01; *d*(:27@CG, :DFH@N5) = 3.54; *d*(:6@CA, :DFH@N4) = 3.63; *d*(:28@CA, :DFH@C1) = 4.32; *d*(:57@CZ, :DFH@C5) = 3.80; *d*(:DFH@C14, :NPH@C3) = 3.19. The following distance restraints were used to refine the active site with QM/MM energy minimization: *d*(:27@CG, : DFH @N5) = 3.54; *d*(:DFH @C14, :NPH@C3) = 3.19. The QM region (see Supplementary Fig. 4c) includes a part of DHF-H+ with the protonated pterin group, a part of NADPH with the dihydronicotinamide group and the side chain of Asp27 (play an important role in facilitating both the proton transfer and hydride transfer steps[69]). The reaction coordinate (RC) was defined as RC $= d$(:NPH@C3, :NPH@H42) $- d$(:NPH@H4, :DFH@C14) and discretized into 31 windows: RC = -2.0, -1.8, -1.6, -1.4, -1.2, -1.0, -0.9, -0.8, -0.7, -0.6, -0.5, -0.4, -0.3, -0.2, -0.1, 0.0, 0.1, 0.2, 0.3, 0.4, 0.5, 0.6, 0.7, 0.8, 0.9, 1.0, 1.2, 1.4, 1.6, 1.8, 2.0. The metastable states P1, P2, and P3 were taken to evaluate $\Delta G_i^{\ddagger}$.

**15. Estimation of disentangling rates and unfolding rates using all-atom simulations**

We simulated 30 independent trajectories of each conformation in fully solvated, unrestrained molecular dynamics at each of the following temperatures, 900 K, 800 K, 700K, 600 K and 550 K to monitor the disentangling process. The setup of the all-atom simulations can be found in Supplementary Methods Section14.4. For DHFR, the same number of simulations were performed for the crystal structure (PDBID 4KJK) to monitor the unfolding process at these temperatures. The systems were first energy minimized and the density and box size were then equilibrated at 310 K with all C-alpha atoms restrained harmonically to maintain the original topology. The production run was then performed at high temperatures without any restraints for each system. The disentangling events were identified as when the fraction of entanglements that gain from the native structure ($G_{\text{gain}}$) is 0 and lasts for at least 1 ns, where $G_{\text{gain}} = \frac{1}{N}\sum_{(i,j)} \Theta\left((i,j) \in nc \cap |g(i,j)| > \left|g^{\text{native}}(i,j)\right|\right)$. While the unfolding events were identified as when $Q$ is less than 0.3 and lasts for at least 1 ns. A list of hit times was obtained from the trajectories by using the thresholds for disentangling and unfolding, respectively. The survival probabilities were then calculated by using the hit times. The disentangling rates ($k_{\text{de}}$) and unfolding rates ($k_{\text{uf}}$) at each temperature were estimated by fitting a single exponential function (see Supplementary equation (4)) to the survival probabilities of the entangled states and native states, respectively. Finally, $k_{\text{de}}$ and $k_{\text{uf}}$ at 298 K were extrapolated by fitting the Arrhenius plot (ln$k$ vs. reciprocal of temperatures) using a quadratic function

$$\ln k = \frac{A}{T^2} + \frac{B}{T} + C, \quad (20)$$

where $A$, $B$ and $C$ are fitting parameters and $A \le 0$ ($A < 0$ refers to the super-Arrhenius behavior and $A = 0$ refers to the linear-Arrhenius behavior).

### 16. Error estimation and statistical tests

The 95% CIs of the raw state probability time courses $p_i^0(t)$ were estimated using the following bootstrapping method: (1) Randomly resample the indices of post-translational metastable state discrete trajectories 10,000 times with replacement; (2) Reconstruct the set of post-translational metastable state discrete trajectories using the resampled indices, resulting in 10,000 resampled ensembles of trajectories for each variant; (3) For each sample, calculate the time series of raw state probability $p_i^0(t)$ , resulting 10,000 resampled $p_i^0(t)$ for each state; (4) At a given time $t$, estimate the lower bound and upper bound of 95% CI as the 2.5% and 97.5% percentiles of the distribution of the 10,000 resampled $p_i^0(t)$ for each variant, respectively.

The 95% CIs of the relative specific activities SA* were estimated using the following bootstrapping method: (1) Create a set of $\Delta G_i^{\ddagger}$ values ($\{\Delta G_i^{\ddagger}\}$) that reproduces the observed distribution probabilities $\{p_i\}$ (with solubility correction) for each variant with a sample size equal to the number of post-translational trajectories; (2) Randomly resample the set $\{\Delta G_i^{\ddagger}\}$ 10,000 times with replacement; (3) For each sample, estimate SA* using the main text equations (4) and (5), where the state probabilities $p_i$ were calculated as the fraction of state *i* in the resampled $\{\Delta G_i^{\ddagger}\}$ set, resulting in 10,000 resampled SA*; (5) Estimate the lower bound and upper bound of 95% CI as the 2.5% and 97.5% percentiles of the distribution of the 10,000 resampled SA* for each variant, respectively.

To test whether or not the observed change of SA* at end of the post-translational simulations is statistically significant, a p-value was calculated for each candidate enzyme using the permutation test method. The null hypothesis is that $SA_{fast}$ and $SA_{slow}$ do not have significant difference; and the alternative hypothesis is that $SA_{fast}$ is lower than $SA_{slow}$ or $SA_{slow}$ is lower than $SA_{fast}$, depending on which variant has the lower observed SA. The permutation test was performed as follows: (1) Create a set of $\Delta G_i^{\ddagger}$ values ($\{\Delta G_i^{\ddagger}\}$) that reproduces the observed distribution probabilities $\{p_i\}$ (with solubility correction) at end of the post-translational simulations for each variant with the sample size equals to the number of post-translational trajectories; (2) Combine $\{\Delta G_i^{\ddagger}\}_{fast}$ and $\{\Delta G_i^{\ddagger}\}_{slow}$ into one set and resample the combined set for 1,000,000 times without replacement, i.e., permutate the indices, resulting 1,000,000 resampled combined sets; (3) For each resampled combined set, partition the first half as the resampled $\{\Delta G_i^{\ddagger}\}_{fast}$ and the second half as the resampled $\{\Delta G_i^{\ddagger}\}_{slow}$, resulting 1,000,000 resampled $\{\Delta G_i^{\ddagger}\}_{fast}$ and $\{\Delta G_i^{\ddagger}\}_{slow}$; (4) For each sample, estimate $SA_{fast}$ and $SA_{slow}$, respectively, using the main text equation (4), where the state probabilities $p_i$ were calculated as the fraction of state *i* in the resampled $\{\Delta G_i^{\ddagger}\}_{\text{fast}}$ and $\{\Delta G_i^{\ddagger}\}_{\text{slow}}$ sets. Calculate $\text{SA}_{\text{slow}} - \text{SA}_{\text{fast}}$ (in the case that the alternative hypothesis is $\text{SA}_{\text{fast}} < \text{SA}_{\text{slow}}$) or $\text{SA}_{\text{fast}} - \text{SA}_{\text{slow}}$ (in the case that the alternative hypothesis is $\text{SA}_{\text{fast}} > \text{SA}_{\text{slow}}$); (5) Estimate the p-value as the frequency of observing the sampled difference that is greater than or equals to the observed difference. If the p-value is less than the significant level 0.05 then we reject the null hypothesis.

The 95% CIs of $k_{\text{de}}^{\text{sim}}$ and $k_{\text{uf}}^{\text{sim}}$ were estimated using the following bootstrapping method: (1) At each temperature, randomly resample the hit times for 10,000 times with replacement; (2) Reconstruct the list of hit times and the survival probability, resulting in 10,000 resampled survival probabilities at each temperature; (3) For each sample of the survival probabilities at all temperatures, extrapolate the rate at 298 K, resulting 10,000 resampled rates; (4) Estimate the lower bound and upper bound of 95% CI as the 2.5% and 97.5% percentiles of the distribution of the 10,000 resampled rates. The 95% CIs of the rescaled disentangling timescale $\tau_{\text{de}}^{\text{rescale}}$ is estimated in the same way by applying the bootstrapping samples of $k_{\text{de}}^{\text{sim}}$ in the equation $\tau_{\text{de}}^{\text{rescale}} = 144/k_{\text{de}}^{\text{sim}}$.

**17. Folding pathway analysis for co- and post-translational simulations**

To analyze the co-translational folding pathways, the synthesis simulations were divided into blocks of nascent chain lengths. Three blocks of nascent chain lengths were used for CAT-III (1-100, 101-203 and 204-213 residues), whereas four and three blocks were used for DDLB (1-90, 91-180,181-296 and 297-306) and DHFR (1-80, 81-149 and 150-159). At each block, 200 clusters were grouped via a *k*-means algorithm[41, 42] based on the order parameters *Q* and *G* that were calculated for all the trajectories of fast and slow variants; and at maximum 3 metastable states were identified via the PCCA+ algorithm[43]. The co-translational discrete trajectories were constructed based on the metastable states assigned in all the blocks. The combined discrete trajectories for co- and post-translational folding were then constructed by appending the post-translational discrete trajectories (obtained in Supplementary Methods Section 12) to the corresponding co-translational discrete trajectories. The folding pathways were identified as follows: (1) For each discrete trajectory, put the starting state of the first frame into the pathway; (2) Move forward along the trajectory and find the state that is different from the last state recoded in the pathway. If the state has not yet been recorded in the pathway, then put it into the pathway. Otherwise, cut the pathway at the first place where this state is recorded and then move forward; (3) Repeat step (2) until the end of the trajectory. This will yield a pathway that has no loop on the route and only records the on-pathway states for each discrete trajectory. The distribution of distinct pathways and the pathway probabilities from one state to another can be estimated from the pathways of all the discrete trajectories.

**18. Structural distribution divergence analysis**

To quantify the divergence of the co- and post-translational structural distributions between the fast variant and slow variant, the Jensen-Shannon divergence (JSD) metric[70] was applied on the probability distributions ($P_i^{\text{fast}}$ and $P_i^{\text{slow}}$) of the microstates ($m$) assigned by the *k*-means clusters at a given nascent chain length *i* and the metastable states ($M$) assigned by coarse-graining the *k*-means clusters at post-translational time step *i*, respectively:

$$\text{JSD}\left(P_i^{\text{fast}} \| P_i^{\text{slow}}\right) = \frac{1}{2} \sum_{x \in m,M} \left[ P_i^{\text{fast}}(x) \ln \frac{P_i^{\text{fast}}(x)}{\frac{1}{2}\left(P_i^{\text{fast}}(x) + P_i^{\text{slow}}(x)\right)} + P_i^{\text{slow}}(x) \ln \frac{P_i^{\text{slow}}(x)}{\frac{1}{2}\left(P_i^{\text{fast}}(x) + P_i^{\text{slow}}(x)\right)} \right] \tag{21}$$

where $P_i^{\text{fast}}$ and $P_i^{\text{slow}}$ were obtained from the last frame of the co-translational simulation trajectories at nascent chain length *i* and from frame *i* of the post-translational simulation trajectories, respectively. For the purpose of better comparison, the post-translational JSD values were normalized to achieve that the starting value is identical to the end value of the co-translational JSD. That is,

$$\text{JSD}_{\text{norm}}^{\text{post-trans}}(i) = \frac{\text{JSD}^{\text{post-trans}}(i)}{\text{JSD}^{\text{post-trans}}(0)} \cdot \text{JSD}^{\text{co-trans}}(N)\,, \tag{22}$$

where *N* is the protein chain length. We choose metastable states for calculating the post-translational divergence because metastable states better represent the structural distribution that is relevant to the enzymatic activity in the post-translation.

**19. Experimental validation for the predicted specific activity of DDLB variants**

**Materials.** All commercial materials were used as received unless otherwise noted. Tris(hydroxymethyl)aminomethane (Tris), sodium chloride, and glycerol was purchased from Fisher Scientific. Imidazole was purchased from J. T. Baker Chemical Co. Isopropyl β-D-1-thiogalactopyranoside (IPTG), and dithiothreitol (DTT) were purchased from Gold Biotechnology. Ni-NTA resin was purchased from Qiagen. 2-Mercaptoethanol, dihydrofolic acid, D-alanine, and pyruvate kinase were purchased from Sigma-Aldrich. Lactate dehydrogenase, and phosphoenol pyruvate (PEP) were purchased from Roche.

**General methods.** UV-visible spectra were recorded on a Cary 60 spectrometer from Varian (Agilent Technologies, Santa Clara, CA) using the WinUV software package to control the instrument.

**Cloning, overexpression and purification of slow and fast translational variants of DDLB.** DNA sequences encoding the slow and fast variants of DDLB were obtained from our simulations and cloned into pET-26b(+) vector (New England Biolabs) containing C-terminal hexahistidine ($His_6$) tag using *Nde*I and *Sac*I restriction sites. The resulting constructs, pET-26b(+)-slow and pET-26(+)-fast DDLB, were verified by DNA sequencing and used to transform *E. coli* BL21 (DE3). A single colony was used to inoculate 200 ml of a lysogeny broth (LB) starter culture containing 50 mg/L kanamycin, which was shaken at 37 °C and 180 rpm for 12 h. 15 mL of this starter culture was used to inoculate 3 L of LB medium containing 50 mg/L kanamycin, which was incubated at 37 °C and 180 rpm until an optical density at 600 nm ($OD_{600}$) of ~0.6 was reached. Protein expression was induced by adding 0.25 mM isopropylthio-β-D-galactoside (IPTG), and incubation was continued at 25 °C at 180 rpm for an additional 12-15 h. The cells were harvested by centrifugation (4 °C, 15000 × *g*, 15 min). For protein purification, the cells were resuspended in lysis buffer (20 mM Tris-HCl, 200 mM NaCl, 5 mM imidazole, 10 mM $MgCl_2$, 2 mM β-mercaptoethanol pH 7.5), and were then disrupted by sonication with an ultrasonic cell disruptor (Branson Sonifier II "Modell W- 250", Heinemann) before the lysates were clarified by centrifugation (4 °C, 45000 × *g*, 45 min). The C-terminally $His_6$-tagged slow and fast DDLB variants were purified via immobilized metal affinity chromatography (IMAC) at 4 °C using Ni-NTA resin. The Ni-NTA resin was pre-equilibrated with lysis buffer. After loading the supernatant onto the Ni-NTA column, the resin was washed with 200 mL wash buffer-A (20 mM Tris-HCl, 200 mM NaCl, 20 mM imidazole, 5 mM $MgCl_2$, 2 mM β-mercaptoethanol pH 7.5) followed by a 50 mL wash with wash buffer-B (20 mM Tris-HCl, 200 mM NaCl, 40 mM imidazole, 5 mM $MgCl_2$, 2 mM β-mercaptoethanol pH 7.5) and a 50 mL wash with wash buffer-C (20 mM Tris-HCl, 200 mM NaCl, 80 mM imidazole, 5 mM $MgCl_2$, 2 mM β-mercaptoethanol pH 7.5). The slow and fast variant proteins were then eluted from the column with 30 mL of elution buffer (20 mM Tris-HCl, 200 mM NaCl, 300 mM imidazole, 2 mM β-mercaptoethanol pH 7.5). The proteins were concentrated using 10000 kDa MWCO filters (Millipore), and imidazole was removed using a PD-10 column (GE Healthcare) pre-equilibrated in storage buffer (20 mM Tris-HCl, 200 mM NaCl, 10% glycerol, 2 mM β-mercaptoethanol pH 7.5) according to the manufacturer's protocol. The enzyme was divided into 50 µL aliquots before it was snap-frozen in liquid $N_2$ and stored at −80 °C until use.

**DDLB kinetic assays.** Enzyme activity was measured spectrophotometrically using a previously described procedure with a few amendments[71, 72, 73]. Reactions contained the following in a final volume of 120 µL: 100 mM Tris–HCl (pH 7.8), 20 mM ATP, 10 mM $MgCl_2$, 10 mM KCl, 0.15 mg/mL lactate dehydrogenase, 0.3 mg/mL pyruvate kinase, 0.3 mM NADH, 7.5 mM phosphoenol pyruvate (PEP), and ~13-14 nM of DDLB with varying concentrations (0 – 8 mM) of D-alanine. Reactions were initiated by addition of D- alanine, and the disappearance of NADH was monitored over 5 min at 25 °C by the loss in absorbance at 340 nm ($\varepsilon_{340}$=6440 $M^{-1}$ $cm^{-1}$). Kinetic assays were conducted in triplicate (technical replicates) and the data were fitted to the Michaelis-Menten equation to extract the values of rate constant ($k_{cat}$) and Michaelis constant ($K_M$).

**Relative specific activity and hypothesis test.** In total 5 biological replicates were prepared and assayed for both the fast and slow variants. For each biological replicate, the relative specific activity of the slow variant with regard to the fast variant was calculated as $k_{cat}^{slow}/k_{cat}^{fast}$. The hypothesis that the average relative specific activity ($\langle k_{cat}^{slow}/k_{cat}^{fast} \rangle$) is less than 1 was tested by using the one-tailed Student's t-test.

**Time-course analysis of overexpression.** A single colony was used to inoculate 200 ml of a lysogeny broth (LB) starter culture containing 50 mg/L kanamycin, which was shaken at 37 °C and 180 rpm for 12 h. 1 mL of this starter culture was used to inoculate 100 mL of LB medium containing 50 mg/L kanamycin, which was incubated at 37 °C and 180 rpm until an optical density at 600 nm ($OD_{600}$) of ~0.6 was reached. Protein expression was induced by adding 0.25 mM isopropylthio-β-D-galactoside (IPTG), and 5 mL aliquots were collected at time points 0.5h, 1h, 2h, 3h, 4h, 5h after IPTG addition and the cells were harvested by centrifugation (4 °C, 4000 × *g*, 15 min). As a control, 5 mL of cells were also collected before IPTG addition. The aliquots were diluted accordingly based on the $OD_{600}$ so that the number of cells is controlled to be same. Protein expression at all time points including control was analyzed by both SDS-PAGE and western blot as the DDLB variants are $His_6$-tagged and the antibodies binds the $His_6$-tag are commercially available and were used to check the expression of both slow and fast variants.

**20. Experimental validation for the predicted specific activity of DHFR variants**

**Cloning, overexpression and purification of slow and fast translational variants of DHFR.** DNA sequences encoding the slow and fast variants of DHFR were obtained from the mRNA sequences used in our simulations (see Supplementary Methods Section 22) and cloned into pET-26b(+) vector (Genescript) containing C-terminal hexahistidine ($His_6$) tag using NdeI and SacI restriction sites. The resulting constructs, pET26b(+)-slow and pET-26b(+)-fast DHFR, were verified by DNA sequencing and used to transform *E. coli* BL21 (DE3). A single colony was used to inoculate 50 ml of a lysogeny broth (LB) starter culture containing 50 mg/L kanamycin, which was shaken at 37 °C and 180 rpm for 12 h. 5mL of this starter culture was used to inoculate each of the three 2L flasks, each containing 1L of LB medium and 50 mg/L kanamycin. Each flask was incubated at 37 °C and 180 rpm until an optical density at 600 nm ($OD_{600}$) of ~0.6 was reached. Protein expression was induced by adding 0.25 mM isopropylthio-β-Dgalactoside (IPTG), and incubation was continued at 25 °C at 180 rpm for an additional 12-15 h. The cells were harvested by centrifugation (4 °C, 15000 × *g*, 15 min). For protein purification, 9 grams of cells were resuspended in lysis buffer (20 mM Tris-HCl, 200 mM NaCl, 5 mM imidazole, 10 mM MgCl2, 2 mM β-mercaptoethanol pH 7.5), and were then disrupted by sonication with an ultrasonic cell disruptor (Branson Sonifier 450) before the lysates were clarified by centrifugation (4 °C, 45000 × *g*, 45 min). The C-terminally $His_6$-tagged slow and fast DHFR variants were purified via immobilized metal affinity chromatography (IMAC) at 4 °C using Ni-NTA resin. The Ni-NTA resin was pre-equilibrated with lysis buffer. After loading the supernatant onto the Ni-NTA column, the resin was washed with 200 mL wash buffer-A (20 mM Tris-HCl, 300 mM NaCl, 20 mM imidazole, 5 mM MgCl2, 2 mM β-mercaptoethanol pH 7.5) followed by a 100 mL wash with wash buffer-B (20 mM Tris-HCl, 200 mM NaCl, 40 mM imidazole, 5 mM MgCl2, 2 mM β-mercaptoethanol pH 7.5) and a 100 mL wash with wash buffer-C (20 mM Tris-HCl, 200 mM NaCl, 80 mM imidazole, 5 mM MgCl2, 2 mM β-mercaptoethanol pH 7.5). The slow and fast variant proteins were then eluted from the column with 80 mL of elution buffer (20 mM Tris-HCl, 200 mM NaCl, 300 mM imidazole, 2 mM β-mercaptoethanol pH 7.5). The proteins were concentrated using 10000 kDa MWCO filters (Cytiva) and exchanged 3 times with the storage buffer containing 10-20% glycerol. The enzyme was divided into 75 µL aliquots before it was snap-frozen in liquid $N_2$ and stored at −80 °C until use. Across all the five biological replicates, the fast variant yields more DHFR proteins than the slow variant (1.3-fold more protein yield on average), indicating that the fast variant indeed translates faster.

**DHFR kinetic assays.** Spectrophotometric enzymatic assay to determine the catalytic constant ($k_{cat}$) of slow and fast DHFR is adapted from the reported protocols by two independent groups namely Benkovic[74] and Skolnick[75] and coworkers with minor modifications. Reactions were performed in a 0.15 mL reaction volume containing 100 mM Tris buffer pH 7.5, 16 nM slow or fast DHFR, 0.5 mM DTT, 100 µM NADPH, 40 µM dihydrofolic acid. The enzyme was preincubated with NADPH for 3 minutes, and then the reaction was initiated by the addition of 40 µM dihydrofolic acid and the decrease in absorbance was monitored at 340 nm ($\Delta\varepsilon_{340}$ = 6220 $M^{-1}$ $cm^{-1}$) for 120 seconds at 25 °C. All measurements were performed in quadruplicate on five independent biological replicates of slow and fast variants of DHFR.

**Relative specific activity and hypothesis test.** In total 5 biological replicates were prepared and assayed for both the fast and slow variants. For each biological replicate, the relative specific activity of the slow variant with regard to the fast variant was calculated as $k_{\mathrm{cat}}^{\mathrm{fast}}/k_{\mathrm{cat}}^{\mathrm{slow}}$. The hypothesis that the average relative specific activity ($\langle k_{\mathrm{cat}}^{\mathrm{fast}}/k_{\mathrm{cat}}^{\mathrm{slow}}\rangle$) is not equal 1 was tested by using the two-tailed Student's t-test.

**21. Limited-proteolysis mass spectrometry experiments for DDLB and DHFR**

Details of the experimental methods are described in Ref. 76 (without molecular chaperones added). Briefly, *E. coli* K12 cells (in which DDLB and DHFR are endogenous proteins) were grown in MOPS defined media[77] in 2 sets of 3 biological replicates. One set was supplemented with 0.5 mM [$^{13}C_6$]L-Arginine and 0.4 mM [$^{13}C_6$]L-Lysine and the other with 0.5 mM L-Arginine and 0.4 mM L-Lysine. Cells were cultured at 37˚C and each heavy/light pair was then pooled together. Cells were collected by centrifugation at 4˚C, supernatants were removed, and cell pellets were stored at -20˚C. Frozen cell pellets were resuspended in a lysis buffer consisting of 900 μL of 20 mM Tris pH 8.2 (20 mM Tris pH 8.2, 100 mM NaCl, 2 mM $MgCl_2$ and supplemented with DNase I to a final concentration (f.c.) of 0.1 mg mL$^{-1}$) and then cryogenically pulverized with a freezer mill (SPEX Sample Prep). Lysates were then clarified (16000 *g* for 15 min at 4 °C) to remove insoluble cell debris and ultracentrifuged to deplete ribosome particles (33,300 rpm for 90 min at 4 °C using a SW55 Ti rotor). Protein concentrations were measured by BCA assay and diluted to a standard concentration of 3.3 mg mL$^{-1}$ using Tris lysis buffer.

To prepare native samples, 3.5 μL of normalized lysates were diluted with 96.5 μL of Tris native dilution buffer (20 mM Tris pH 8.2, 100 mM NaCl, 10.288 mM $MgCl_2$, 10.36 mM KCl, 2.07 mM ATP, 1.04 mM DTT, 62 mM GdmCl) to a final protein concentration of 0.115 mg mL$^{-1}$. Native samples were then equilibrated by incubating for 90 min at room temperature. To prepare unfolded samples, 600 μL of normalized lysates, 100 mg of solid guanidinium chloride, and 2.4 μL of a freshly prepared 700 mM DTT stock solution were combined, and solvent was removed using a vacufuge plus to a final volume of 170 μL. Unfolded lysates were incubated overnight at room temperature. 99 μL of refolding dilution buffer were rapidly added to 1 μL of unfolded extract to let it refold. Refolded samples were then incubated at room temperature for 1 min, 5 min or 2 h.

100 µL of the native or refolded lysates was added to Proteinase K (enzyme:substrate ratio of 1:100 w/w ratio[78]), incubated for 1 min at room temperature, and quenched by boiling in a mineral oil bath at 110°C for 5 min. Boiled samples were transferred to tubes containing 76 mg urea. To prepare samples for mass spectrometry, dithiothreitol was added to a final concentration of 10 mM and samples were incubated at 37°C for 30 minutes. Iodoacetamide was added to a final concentration of 40 mM and samples were incubated at room temperature in the dark for 45 minutes. LysC was added to a 1:100 enzyme:substrate (w/w) ratio and samples were incubated at 37˚C for 2 h, urea was diluted to 2 M using 100 mM ammonium bicarbonate pH 8, then trypsin was added to a 1:50 enzyme:substrate (w/w) ratio and incubated overnight at 25°C.

Peptides were acidified, desalted with Sep-Pak C18 1 cc Vac Cartridges, dried down, and resuspend in 0.1% formic acid. LC-MS/MS acquisition was conducted on a Thermo Ultimate3000 UHPLC system with an Acclaim Pepmap RSLC C18 column (75 µm × 25 cm, 2 µm, 100 Å) in line with a Thermo Q-Exctive HF-X Orbitrap.

Proteome Discoverer (PD) Software Suite (v2.4, Thermo Fisher) and the Minora Algorithm were used to analyze mass spectra and perform Label Free Quantification (LFQ) of detected peptides. Details pertaining to the settings used for the analysis nodes are specified elsewhere[76]. The data were searched against *Escherichia coli* (UP000000625, Uniprot) reference proteome database. For peptide identification, the PD MSFragger node was used, using a semi-tryptic search allowing up to 2 missed cleavages[79]. A precursor mass tolerance of 10 ppm was used for the MS1 level, and a fragment ion tolerance was set to 0.02 Da at the MS2 level for both search algorithms. Peptide lengths between 7 and 50 amino acid residues was allowed with a peptide mass between 500 and 5000 Da. Additionally, a maximum charge state for theoretical fragments was set at 2. Oxidation of methionine and acetylation of the N-terminus were allowed as dynamic modifications while carbamidomethylation on cysteines was set as a static modification. Heavy isotope labeling ($^{13}C_6$) of Arginine and Lysine were allowed as dynamic modifications. The Philosopher PD node was used for FDR validation. Raw normalized extracted ion intensity data for the identified peptides were exported from the .pdResult file using a three-level hierarchy (protein > peptide group > consensus feature). These data were further processed utilizing custom Python analyzer scripts (available on GitHub: https://github.com/FriedLabJHU/Refoldibility-Tools/, and described in depth previously[80]). Briefly, normalized ion counts were collected across the refolded replicates and the native replicates for each successfully identified peptide group. Effect sizes are the ratio of averages (reported in $\log_2$) and p-values (reported as $-\log_{10}$) were assessed using *t* tests with Welch's correction for unequal population variances. Coefficients of variation (CV) for the peptide abundance in the three replicate refolded samples are also calculated. Analyzer returns a file listing all the peptides that can be confidently quantified, and provides their effect-size, p-value, refolded CV, proteinase K site (if half-tryptic), and associated protein metadata. The data corresponding to DDLB and DHFR were obtained from the output file and presented in Supplementary Tables 13 and 14, respectively.

**22. mRNA sequences used in this study**

All the mRNA sequences used in the simulations and experiments are presented below. All sequences begin with the start codon and end with the stop codon.

**CAT-III fast variant:**

```
AUGAAUUAUACUAAAUUUGAUGUUAAAAAUUGGGUUCGUCGUGAGCACUUUGAGUUUUAUCGUCACCGUC
UGCCGUGUGGUUUUUCUCUGACUUCUAAAAUUGAUAUUACUACUCUGAAAAAAUCUCUGGAUGAUUCUGC
UUAUAAAUUUUAUCCGGUUAUGAUUUAUCUGAUUGCUCAAGCUGUUAAUCAAUUUGAUGAGCUGCGUAUG
GCUAUUAAAGAUGAUGAGCUGAUUGUUUGGGAUUCUGUUGAUCCGCAAUUUACUGUUUUUCACCAAGAGA
CUGAGACUUUUUCUGCUCUGUCUUGUCCGUAUUCUUCUGAUAUUGAUCAAUUUAUGGUUAAUUAUCUGUC
UGUUAUGGAGCGUUAUAAAUCUGAUACUAAACUGUUUCCGCAAGGUGUUACUCCGGAGAAUCACCUGAAU
AUUUCUGCUCUGCCGUGGGUUAAUUUUGAUUCUUUUAAUCUGAAUGUUGCUAAUUUUACUGAUUAUUUUG
CUCCGAUUAUUACUAUGGCUAAAUAUCAACAAGAGGGUGAUCGUCUGCUGCUGCCGCUGUCUGUUCAAGU
UCACCACGCUGUUUGUGAUGGUUUUCACGUUGCUCGUUUUAUUAAUCGUCUGCAAGAGCUGUGUAAUUCU
AAACUGAAAUAA
```

**CAT-III slow variant:**

```
AUGAACUACACAAAGUUCGACGUCAAGAACUGGGUCAGGAGGGAACAUUUCGAAUUCUACAGGCAUAGGC
UACCAUGCGGAUUCUCCCUAACAUCCAAGAUCGACAUCACAACACUAAAGAAGUCCCUAGACGACUCCGC
CUACAAGUUCUACCCAGUCAUGAUCUACCUAAUCGCCCAGGCCGUCAACCAGUUCGACGAACUAAGGAUG
GCCAUCAAGGACGACGAACUAAUCGUCUGGGACUCCGUCGACCCACAGUUCACAGUCUUCCAUCAGGAAA
CAGAAACAUUCUCCGCCCUAUCCUGCCCAUACUCCUCCGACAUCGACCAGUUCAUGGUCAACUACCUAUC
CGUCAUGGAAAGGUACAAGUCCGACACAAAGCUAUUCCCACAGGGAGUCACACCAGAAAACCAUCUAAAC
AUCUCCGCCCUACCAUGGGUCAACUUCGACUCCUUCAACCUAAACGUCGCCAACUUCACAGACUACUUCG
CCCCAAUCAUCACAAUGGCCAAGUACCAGCAGGAAGGAGACAGGCUACUACUACCACUAUCCGUCCAGGU
CCAUCAUGCCGUCUGCGACGGAUUCCAUGUCGCCAGGUUCAUCAACAGGCUACAGGAACUAUGCAACUCC
AAGCUAAAGUAG
```

**DDLB fast variant:**

AUGACUGAUAAAAUUGCUGUUCUGCUGGGUGGUACUUCUGCUGAGCGUGAGGUUUCUCUGAAUUCUGGUG
CUGCUGUUCUGGCUGGUCUGCGUGAGGGUGGUAUUGAUGCUUAUCCGGUUGAUCCGAAAGAGGUUGAUGU
UACUCAACUGAAAUCUAUGGGUUUUCAAAAAGUUUUUAUUGCUCUGCACGGUCGUGGUGGUGAGGAUGGU
ACUCUGCAAGGUAUGCUGGAGCUGAUGGGUCUGCCGUAUACUGGUUCUGGUGUUAUGGCUUCUGCUCUGU
CUAUGGAUAAACUGCGUUCUAAACUGCUGUGGCAAGGUGCUGGUCUGCCGGUUGCUCCGUGGGUUGCUCU
GACUCGUGCUGAGUUUGAGAAAGGUCUGUCUGAUAAACAACUGGCUGAGAUUUCUGCUCUGGGUCUGCCG
GUUAUUGUUAAACCGUCUCGUGAGGGUUCUUCUGUUGGUAUGUCUAAAGUUGUUGCUGAGAAUGCUCUGC
AAGAUGCUCUGCGUCUGGCUUUUCAACACGAUGAGGAGGUUCUGAUUGAGAAAUGGCUGUCUGGUCCGGA
GUUUACUGUUGCUAUUCUGGGUGAGGAGAUUCUGCCGUCUAUUCGUAUUCAACCGUCUGGUACUUUUUAU
GAUUAUGAGGCUAAAUAUCUGUCUGAUGAGACUCAAUAUUUUGUCCGGCUGGUCUGGAGGCUUCUCAAG
AGGCUAAUCUGCAAGCUCUGGUUCUGAAAGCUUGGACUACUCUGGGUUGUAAAGGUUGGGGUCGUAUUGA
UGUUAUGCUGGAUUCUGAUGGUCAAUUUUAUCUGCUGGAGGCUAAUACUUCUCCGGGUAUGACUUCUCAC
UCUCUGGUUCCGAUGGCUGCUCGUCAAGCUGGUAUGUCUUUUUCUCAACUGGUUGUUCGUAUUCUGGAGC
UGGCUGAUUAA

**DDLB slow variant:**

AUGACAGACAAGAUCGCCGUCCUACUAGGAGGAACAUCCGCCGAAAGGGAAGUCUCCCUAAACUCCGGAG
CCGCCGUCCUAGCCGGACUAAGGGAAGGAGGAAUCGACGCCUACCCAGUCGACCCAAAGGAAGUCGACGU
CACACAGCUAAAGUCCAUGGGAUUCCAGAAGGUCUUCAUCGCCCUACAUGGAAGGGGAGGAGAAGACGGA
ACACUACAGGGAAUGCUAGAACUAAUGGGACUACCAUACACAGGAUCCGGAGUCAUGGCCUCCGCCCUAU
CCAUGGACAAGCUAAGGUCCAAGCUACUAUGGCAGGGAGCCGGACUACCAGUCGCCCCAUGGGUCGCCCU
AACAAGGGCCGAAUUCGAAAAGGGACUAUCCGACAAGCAGCUAGCCGAAAUCUCCGCCCUAGGACUACCA
GUCAUCGUCAAGCCAUCCAGGGAAGGAUCCUCCGUCGGAAUGUCCAAGGUCGUCGCCGAAAACGCCCUAC
AGGACGCCCUAAGGCUAGCCUUCCAGCAUGACGAAGAAGUCCUAAUCGAAAAGUGGCUAUCCGGACCAGA
AUUCACAGUCGCCAUCCUAGGAGAAGAAAUCCUACCAUCCAUCAGGAUCCAGCCAUCCGGAACAUUCUAC
GACUACGAAGCCAAGUACCUAUCCGACGAAACACAGUACUUCUGCCCAGCCGGACUAGAAGCCUCCCAGG
AAGCCAACCUACAGGCCCUAGUCCUAAAGGCCUGGACAACACUAGGAUGCAAGGGAUGGGGAAGGAUCGA
CGUCAUGCUAGACUCCGACGGACAGUUCUACCUACUAGAAGCCAACACAUCCCCAGGAAUGACAUCCCAU
UCCCUAGUCCCAAUGGCCGCCAGGCAGGCCGGAAUGUCCUUCUCCCAGCUAGUCGUCAGGAUCCUAGAAC
UAGCCGACUAG

**DHFR fast variant:**

AUGAUUUCUCUGAUUGCUGCUCUGGCUGUUGAUCGUGUUAUUGGUAUGGAGAAUGCUAUGCCGUGGAAUC
UGCCGGCUGAUCUGGCUUGGUUUAAACGUAAUACUCUGAAUAAACCGGUUAUUAUGGGUCGUCACACUUG
GGAGUCUAUUGGUCGUCCGCUGCCGGGUCGUAAAAAUAUUAUUCUGUCUUCUCAACCGGGUACUGAUGAU

CGUGUUACUUGGGUUAAAUCUGUUGAUGAGGCUAUUGCUGCUUGUGGUGAUGUUCCGGAGAUUAUGGUUA
UUGGUGGUGGUCGUGUUUAUGAGCAAUUUCUGCCGAAAGCUCAAAAACUGUAUCUGACUCACAUUGAUGC
UGAGGUUGAGGGUGAUACUCACUUUCCGGAUUAUGAGCCGGAUGAUUGGGAGUCUGUUUUUUCUGAGUUU
CACGAUGCUGAUGCUCAAAAUUCUCACUCUUAUUGUUUUGAGAUUCUGGAGCGUCGUUAA

**DHFR slow variant:**

AUGAUCUCCCUAAUCGCCGCCCUAGCCGUCGACAGGGUCAUCGGAAUGGAAAACGCCAUGCCAUGGAACC
UACCAGCCGACCUAGCCUGGUUCAAGAGGAACACACUAAACAAGCCAGUCAUCAUGGGAAGGCAUACAUG
GGAAUCCAUCGGAAGGCCACUACCAGGAAGGAAGAACAUCAUCCUAUCCUCCCAGCCAGGAACAGACGAC
AGGGUCACAUGGGUCAAGUCCGUCGACGAAGCCAUCGCCGCCUGCGGAGACGUCCCAGAAAUCAUGGUCA
UCGGAGGAGGAAGGGUCUACGAACAGUUCCUACCAAAGGCCCAGAAGCUAUACCUAACACAUAUCGACGC
CGAAGUCGAAGGAGACACACAUUUCCCAGACUACGAACCAGACGACUGGGAAUCCGUCUUCUCCGAAUUC
CAUGACGCCGACGCCCAGAACUCCCAUUCCUACUGCUUCGAAAUCCUAGAAAGGAGGUAG

### Supplementary Tables

Supplementary Table 1. Force field parameters of the CG protein model

| Energy term | | Parameters |
|---|---|---|
| $E_{\text{bond}}$ | $K_{\text{b}}$ (kcal/mol/Å$^2$) | 50 |
| | $b_0$ (Å) | 3.81 |
| $E_{\text{angle}}$ | $\gamma$ (mol/kcal) | 0.1 |
| | $K_{\alpha}$ (kcal/mol/rad$^2$) | 106.4 |
| | $\theta_{\alpha}$ (rad) | 1.60 |
| | $\varepsilon_{\alpha}$ (kcal/mol) | 4.3 |
| | $K_{\beta}$ (kcal/mol/rad$^2$) | 26.3 |
| | $\theta_{\beta}$ (rad) | 2.27 |
| $E_{\text{dihedral}}$ | $K_{\text{D}_1}$ (kcal/mol/rad$^2$) | 0.6309 |
| | $\delta_1$ (rad) | 5.3291 |
| | $K_{\text{D}_2}$ (kcal/mol/rad$^2$) | 1.0621 |
| | $\delta_2$ (rad) | 4.5574 |
| | $K_{\text{D}_3}$ (kcal/mol/rad$^2$) | 0.045943 |
| | $\delta_3$ (rad) | 4.8229 |
| | $K_{\text{D}_4}$ (kcal/mol/rad$^2$) | 0.076257 |
| | $\delta_4$ (rad) | 2.1 |
| $E_{\text{elec}}$ | $q_i$ ($e$) | Amino acid dependent |
| | $\varepsilon_{\gamma}$ | 78.5 |
| | $l_{\text{D}}$ (Å) | 10 |
| $E_{\text{vdW}}$ | Native contacts [a] | |
| | $\varepsilon_{ij}$ (kcal/mol) | $\varepsilon_{ij}^{\text{HB}} + n_{\text{scal}} \cdot \varepsilon_{ij}^{\text{SC-SC}} + \varepsilon_{ij}^{\text{BB-SC}}$ |
| | $R_{ij}$ (Å) | $d_{ij}$ |
| | Non-native contacts [b] | |
| | $\varepsilon_{ij}$ (kcal/mol) | 0.000132 |
| | $R_{ij}$ (Å) | $\frac{2^{1/6}}{2}(\sigma_i + \sigma_j)$ |

[a] Native contacts include three terms: hydrogen bond (HB), sidechain-sidechain (SC-SC) and backbone-sidechain (BB-SC). The potential scaling factor $n_{\text{scal}}$ need to be further parameterized for a particular protein. $d_{ij}$ is the native contact distance.

[b] Non-native contacts have a fixed potential well depth. $\sigma_i$ is the collision diameter of the CG interaction site *i*.

Supplementary Table 2a. The protein name, PDB ID, protein length, experimental temperature $T^{\mathrm{exp}}$, experimental protein stability $\Delta G_{\mathrm{UN}}^{\mathrm{exp}}$, $n_{\mathrm{scal}}$ values used in pt-REMD, melting temperature $T_{\mathrm{m}}$, threshold used to identify folding states $Q_{\mathrm{eq}}$, estimated protein stability $\Delta G_{\mathrm{UN}}^{\mathrm{sim}}(T^{\mathrm{exp}})$, linear regression slope *m*, intercept *b*, coefficient of determination $R^2$ and optimized $n_{\mathrm{scal}}^{*}$ of α-class training-set proteins

| Protein | PDB ID | Length | Experimental parameters | | pt-REMD input and output parameters | | | | Linear regression parameters | | | |
|---|---|---|---|---|---|---|---|---|---|---|---|---|
| | | | $T^{\mathrm{exp}}$ (K) | $\Delta G_{\mathrm{UN}}^{\mathrm{exp}}$ (kcal/mol) | $n_{\mathrm{scal}}$ | $T_{\mathrm{m}}$ (K) | $Q_{\mathrm{eq}}$ | $\Delta G_{\mathrm{UN}}^{\mathrm{sim}}(T^{\mathrm{exp}})$ (kcal/mol) | *m* | *b* | $R^2$ | $n_{\mathrm{scal}}^{*}$ |
| EC298 | 1RYK | 69 | 298 | -2.72 [b] | 1.1 | 315.0 | 0.68 | -2.807 | -19.848 | 18.839 | 0.9952 | 1.0862 |
| | | | | | 1.2 | 331.1 | 0.66 | -5.119 | | | | |
| | | | | | 1.3 | 346.9 | 0.65 | -7.248 | | | | |
| | | | | | 1.4 | 361.5 | 0.64 | -8.713 | | | | |
| λ-repressor$_{6-85}$ | 1LMB | 80 | 298 | -4.25 [b] | 1.2 | 314.5 | 0.53 | -3.941 | -20.555 | 20.797 | 0.9995 | 1.2185 |
| | | | | | 1.3 | 330.0 | 0.54 | -5.802 | | | | |
| | | | | | 1.4 | 345.0 | 0.51 | -8.010 | | | | |
| | | | | | 1.5 | 359.8 | 0.50 | -10.057 | | | | |
| bACBP [a] | 2ABD | 86 | 298 | -6.40 [b] | 1.2 | 313.2 | 0.69 | -3.711 | -17.553 | 17.325 | 0.9970 | 1.3516 |
| | | | | | 1.3 | 327.3 | 0.69 | -5.406 | | | | |
| | | | | | 1.4 | 340.3 | 0.65 | -7.505 | | | | |
| | | | | | 1.5 | 354.8 | 0.67 | -8.863 | | | | |
| IM7 | 1CEI | 85 | 298 | -2.88 [b] | 1.0 | 305.8 | 0.62 | -1.809 | -16.987 | 14.735 | 0.9745 | 1.0369 |
| | | | | | 1.1 | 325.3 | 0.60 | -4.677 | | | | |
| | | | | | 1.2 | 342.8 | 0.60 | -5.528 | | | | |
| | | | | | 1.3 | 361.4 | 0.58 | -7.188 | | | | |
| Im9 | 1IMQ | 86 | 298 | -6.81 [c] | 1.1 | 309.7 | 0.53 | -2.458 | -18.194 | 17.111 | 0.9806 | 1.3147 |
| | | | | | 1.2 | 326.4 | 0.51 | -5.300 | | | | |
| | | | | | 1.3 | 340.6 | 0.51 | -6.723 | | | | |
| | | | | | 1.4 | 357.2 | 0.50 | -8.048 | | | | |
| Cytochrome B562 [a] | 256B | 106 | 293 | -6.60 [c] | 1.1 | 310.6 | 0.58 | -5.008 | -27.403 | 25.313 | 0.9982 | 1.1646 |
| | | | | | 1.2 | 322.8 | 0.59 | -7.402 | | | | |
| | | | | | 1.3 | 335.5 | 0.58 | -10.116 | | | | |
| | | | | | 1.4 | 347.5 | 0.57 | -13.238 | | | | |
| Average | | | | | | | | | | | | 1.1954 |

[a] Insufficiently sampled proteins whose pt-REMD results are the averages of those obtained from pt-REMD simulations starting from their native states and unfolded states. See details in Supplementary Table 3.

[b] Obtained from De Sancho et al.[11];

[c] Obtained from Leininger et al.[5];

Supplementary Table 2b. The protein name, PDB ID, protein length, experimental temperature $T^{\text{exp}}$, experimental protein stability $\Delta G_{\text{UN}}^{\text{exp}}$, $n_{\text{scal}}$ values used in pt-REMD, melting temperature $T_{\text{m}}$, threshold used to identify folding states $Q_{\text{eq}}$, estimated protein stability $\Delta G_{\text{UN}}^{\text{sim}}(T^{\text{exp}})$, linear regression slope *m*, intercept *b*, coefficient of determination $R^2$ and optimized $n_{\text{scal}}^{*}$ of β-class training-set proteins

| Protein | PDB ID | Length | Experimental parameters | | pt-REMD input and output parameters | | | | Linear regression parameters | | | |
|---|---|---|---|---|---|---|---|---|---|---|---|---|
| | | | $T^{\text{exp}}$ (K) | $\Delta G_{\text{UN}}^{\text{exp}}$ (kcal/mol) | $n_{\text{scal}}$ | $T_{\text{m}}$ (K) | $Q_{\text{eq}}$ | $\Delta G_{\text{UN}}^{\text{sim}}(T^{\text{exp}})$ (kcal/mol) | *m* | *b* | $R^2$ | $n_{\text{scal}}^{*}$ |
| ABP1 SH3 | 1JO8 | 58 | 298 | -3.07 [a] | 1.4 | 320.2 | 0.77 | -2.042 | -3.907 | 3.381 | 0.9952 | 1.6514 |
| | | | | | 1.5 | 334.6 | 0.78 | -2.535 | | | | |
| | | | | | 1.6 | 348.3 | 0.78 | -2.895 | | | | |
| | | | | | 1.7 | 362.7 | 0.78 | -3.224 | | | | |
| Fyn SH3 | 1SHF | 59 | 293 | -6.68 [a] | 1.2 | 307.6 | 0.54 | -2.888 | -12.596 | 12.167 | 0.9987 | 1.4962 |
| | | | | | 1.3 | 320.4 | 0.55 | -4.327 | | | | |
| | | | | | 1.4 | 335.4 | 0.55 | -5.410 | | | | |
| | | | | | 1.5 | 351.0 | 0.52 | -6.726 | | | | |
| CspB Bc | 1C9O | 66 | 298 | -1.81 [b] | 1.3 | 304.8 | 0.51 | -0.949 | -16.429 | 20.274 | 0.9969 | 1.3442 |
| | | | | | 1.4 | 319.6 | 0.49 | -2.8442 | | | | |
| | | | | | 1.5 | 334.7 | 0.49 | -4.5401 | | | | |
| | | | | | 1.6 | 349.0 | 0.49 | -5.8598 | | | | |
| CspA | 1MJC | 69 | 298 | -3.00 [c] | 1.4 | 307.9 | 0.50 | -1.6971 | -16.644 | 21.483 | 0.9980 | 1.4709 |
| | | | | | 1.5 | 320.8 | 0.51 | -3.6163 | | | | |
| | | | | | 1.6 | 334.2 | 0.50 | -5.2519 | | | | |
| | | | | | 1.7 | 347.3 | 0.49 | -6.7000 | | | | |
| Tenascin | 1TEN | 89 | 298 | -6.71 [a] | 1.3 | 307.4 | 0.41 | -1.6397 | -29.161 | 36.208 | 0.9999 | 1.4718 |
| | | | | | 1.4 | 323.9 | 0.39 | -4.7050 | | | | |
| | | | | | 1.5 | 338.7 | 0.39 | -7.5436 | | | | |
| | | | | | 1.6 | 354.4 | 0.39 | -10.4139 | | | | |
| Twitchin | 1WIT | 93 | 293 | -5.00 [a] | 1.3 | 310.1 | 0.46 | -3.0763 | -18.441 | 20.899 | 0.9995 | 1.4044 |
| | | | | | 1.4 | 324.6 | 0.46 | -4.9700 | | | | |
| | | | | | 1.5 | 338.7 | 0.45 | -6.6515 | | | | |
| | | | | | 1.6 | 353.3 | 0.44 | -8.6626 | | | | |
| Average | | | | | | | | | | | | 1.4732 |

[a] Obtained from De Sancho et al.[11];

[b] Obtained from Leininger et al.[5];

[c] Obtained from Reid et al.[12];

Supplementary Table 2c. The protein name, PDB ID, protein length, experimental temperature $T^{\text{exp}}$, experimental protein stability $\Delta G_{\text{UN}}^{\text{exp}}$, $n_{\text{scal}}$ values used in pt-REMD, melting temperature $T_{\text{m}}$, threshold used to identify folding states $Q_{\text{eq}}$, estimated protein stability $\Delta G_{\text{UN}}^{\text{sim}}(T^{\text{exp}})$, linear regression slope *m*, intercept *b*, coefficient of determination $R^2$ and optimized $n_{\text{scal}}^{*}$ of α/β-class training-set proteins

| Protein | PDB ID | Length | Experimental parameters | | pt-REMD input and output parameters | | | | Linear regression parameters | | | |
|---|---|---|---|---|---|---|---|---|---|---|---|---|
| | | | $T^{\text{exp}}$ (K) | $\Delta G_{\text{UN}}^{\text{exp}}$ (kcal/mol) | $n_{\text{scal}}$ | $T_{\text{m}}$ (K) | $Q_{\text{eq}}$ | $\Delta G_{\text{UN}}^{\text{sim}}(T^{\text{exp}})$ (kcal/mol) | *m* | *b* | $R^2$ | $n_{\text{scal}}^{*}$ |
| Hpr [a] | 1POH | 85 | 298 | -5.46 [b] | 1.0 | 297.3 | 0.51 | 0.165 | -26.153 | 25.765 | 0.9854 | 1.1940 |
| | | | | | 1.1 | 313.1 | 0.50 | -3.709 | | | | |
| | | | | | 1.2 | 327.6 | 0.50 | -5.864 | | | | |
| | | | | | 1.3 | 342.4 | 0.50 | -7.834 | | | | |
| Urm1 [a] | 2QJL | 99 | 298 | -3.48 [c] | 1.1 | 307.9 | 0.46 | -2.786 | -25.491 | 24.723 | 0.9854 | 1.1064 |
| | | | | | 1.2 | 325.8 | 0.46 | -6.506 | | | | |
| | | | | | 1.3 | 342.9 | 0.47 | -8.729 | | | | |
| | | | | | 1.4 | 359.6 | 0.48 | -10.543 | | | | |
| Src SH2 [a] | 1SPR | 103 | 298 | -7.24 [c] | 1.1 | 307.6 | 0.73 | -2.783 | -16.597 | 15.283 | 0.9910 | 1.3571 |
| | | | | | 1.2 | 324.6 | 0.74 | -4.740 | | | | |
| | | | | | 1.3 | 342.4 | 0.71 | -6.651 | | | | |
| | | | | | 1.4 | 359.0 | 0.71 | -7.678 | | | | |
| Azurin (apo) [a] | 1E65 | 128 | 298 | -5.29 [c] | 1.1 | 306.8 | 0.70 | -3.489 | -19.738 | 17.942 | 0.9795 | 1.1770 |
| | | | | | 1.2 | 321.6 | 0.66 | -6.424 | | | | |
| | | | | | 1.3 | 338.4 | 0.69 | -7.199 | | | | |
| | | | | | 1.4 | 355.6 | 0.65 | -9.810 | | | | |
| CheY [a] | 3CHY | 128 | 298 | -5.20 [d] | 1.0 | 309.0 | 0.50 | -3.906 | -64.410 | 60.542 | 0.9999 | 1.0207 |
| | | | | | 1.1 | 328.1 | 0.45 | -10.233 | | | | |
| | | | | | 1.2 | 347.2 | 0.43 | -16.788 | | | | |
| | | | | | 1.3 | 365.6 | 0.43 | $-\infty$ [f] | | | | |
| Ribonuclease H [a] | 2RN2 | 155 | 298 | -7.20 [e] | 1.0 | 310.5 | 0.23 | -3.924 | -42.080 | 38.167 | 0.9999 | 1.0781 |
| | | | | | 1.1 | 327.7 | 0.23 | -8.150 | | | | |
| | | | | | 1.2 | 344.0 | 0.22 | -12.230 | | | | |
| | | | | | 1.3 | 361.1 | 0.22 | -16.591 | | | | |
| Average | | | | | | | | | | | | 1.1556 |

[a] Insufficiently sampled proteins whose pt-REMD results are the averages of those obtained from pt-REMD simulations starting from their native states and unfolded states. See details in Table 3;

[b] Obtained from Nicholson et al.[13];

[c] Obtained from De Sancho et al.[11];

[d] Obtained from López-Hernéndez & Serrano[14];

[e] Obtained from Leininger et al.[5];

[f] Infinite $\Delta G_{\text{UN}}^{\text{sim}}$ due to the extremely high stability at this $n_{\text{scal}}$ value. It was considered as an outlier and not used in the linear regression.

Supplementary Table 3. Insufficiently sampled training-set proteins and their pt-REMD results

| Protein | PDB ID | REX input and output parameters | | | | | Average | | |
|---|---|---|---|---|---|---|---|---|---|
| | | $n_{\mathrm{scal}}$ | Starting state | $T_{\mathrm{m}}$ (K) | $Q_{\mathrm{eq}}$ | $\Delta G_{\mathrm{UN}}^{\mathrm{sim}}(T^{\mathrm{exp}})$ (kcal/mol) | $T_{\mathrm{m}}$ (K) | $Q_{\mathrm{eq}}$ | $\Delta G_{\mathrm{UN}}^{\mathrm{sim}}(T^{\mathrm{exp}})$ (kcal/mol) |
| bACBP | 2ABD | 1.2 | folded | 314.0 | 0.68 | -3.902 | 313.2 | 0.69 | -3.711 |
| | | | unfolded | 312.3 | 0.70 | -3.521 | | | |
| | | 1.3 | folded | 327.2 | 0.70 | -5.163 | 327.3 | 0.69 | -5.406 |
| | | | unfolded | 327.4 | 0.67 | -5.650 | | | |
| | | 1.4 | folded | 340.9 | 0.67 | -7.368 | 340.3 | 0.65 | -7.505 |
| | | | unfolded | 339.6 | 0.64 | -7.641 | | | |
| | | 1.5 | folded | 355.3 | 0.67 | -8.865 | 354.8 | 0.67 | -8.863 |
| | | | unfolded | 354.3 | 0.67 | -8.861 | | | |
| Cytochrome B562 | 256B | 1.1 | folded | 311.1 | 0.58 | -5.131 | 310.6 | 0.58 | -5.008 |
| | | | unfolded | 310.0 | 0.59 | -4.885 | | | |
| | | 1.2 | folded | 323.4 | 0.59 | -7.521 | 322.8 | 0.59 | -7.402 |
| | | | unfolded | 322.1 | 0.59 | -7.283 | | | |
| | | 1.3 | folded | 335.5 | 0.58 | -10.011 | 335.5 | 0.58 | -10.116 |
| | | | unfolded | 335.4 | 0.57 | -10.222 | | | |
| | | 1.4 | folded | 347.4 | 0.57 | -13.067 | 347.5 | 0.57 | -13.238 |
| | | | unfolded | 347.5 | 0.57 | -13.408 | | | |
| Hpr | 1POH | 1.0 | folded | 297.3 | 0.51 | 0.167 | 297.3 | 0.51 | 0.165 |
| | | | unfolded | 297.3 | 0.51 | 0.163 | | | |
| | | 1.1 | folded | 313.1 | 0.50 | -3.682 | 313.1 | 0.50 | -3.709 |
| | | | unfolded | 313.0 | 0.50 | -3.735 | | | |
| | | 1.2 | folded | 328.0 | 0.50 | -5.992 | 327.6 | 0.50 | -5.864 |
| | | | unfolded | 327.2 | 0.50 | -5.736 | | | |
| | | 1.3 | folded | 342.7 | 0.49 | -8.068 | 342.4 | 0.50 | -7.834 |
| | | | unfolded | 342.0 | 0.50 | -7.600 | | | |
| Urm1 | 2QJL | 1.1 | folded | 307.9 | 0.49 | -2.785 | 307.9 | 0.46 | -2.786 |
| | | | unfolded | 307.8 | 0.44 | -2.788 | | | |
| | | 1.2 | folded | 326.2 | 0.46 | -6.497 | 325.8 | 0.46 | -6.506 |
| | | | unfolded | 325.4 | 0.47 | -6.515 | | | |
| | | 1.3 | folded | 342.5 | 0.48 | -8.519 | 342.9 | 0.47 | -8.729 |
| | | | unfolded | 343.2 | 0.46 | -8.939 | | | |
| | | 1.4 | folded | 359.4 | 0.49 | -10.338 | 359.6 | 0.48 | -10.543 |
| | | | unfolded | 359.7 | 0.48 | -10.747 | | | |
| Src SH2 | 1SPR | 1.1 | folded | 307.5 | 0.74 | -2.676 | 307.6 | 0.73 | -2.783 |
| | | | unfolded | 307.6 | 0.72 | -2.890 | | | |
| | | 1.2 | folded | 325.0 | 0.73 | -4.752 | 324.6 | 0.74 | -4.740 |
| | | | unfolded | 324.1 | 0.74 | -4.728 | | | |
| | | 1.3 | folded | 342.0 | 0.71 | -6.686 | 342.4 | 0.71 | -6.651 |
| | | | unfolded | 342.8 | 0.71 | -6.617 | | | |
| | | 1.4 | folded | 358.8 | 0.72 | -7.635 | 359.0 | 0.71 | -7.678 |
| | | | unfolded | 359.2 | 0.71 | -7.721 | | | |

| | | | | | | | | | |
|---|---|---|---|---|---|---|---|---|---|
| Azurin (apo) | 1E65 | 1.1 | folded | 310.0 | 0.68 | -4.109 | 306.8 | 0.70 | -3.489 |
| | | | unfolded | 303.6 | 0.71 | -2.869 | | | |
| | | 1.2 | folded | 325.0 | 0.68 | -6.002 | 321.6 | 0.66 | -6.424 |
| | | | unfolded | 318.2 | 0.65 | -6.846 | | | |
| | | 1.3 | folded | 341.1 | 0.67 | -7.754 | 338.4 | 0.69 | -7.199 |
| | | | unfolded | 335.6 | 0.70 | -6.643 | | | |
| | | 1.4 | folded | 355.9 | 0.64 | -10.361 | 355.6 | 0.65 | -9.810 |
| | | | unfolded | 355.3 | 0.67 | -9.259 | | | |
| CheY | 3CHY | 1.0 | folded | 308.9 | 0.50 | -3.920 | 309.0 | 0.50 | -3.906 |
| | | | unfolded | 309.0 | 0.50 | -3.892 | | | |
| | | 1.1 | folded | 328.2 | 0.45 | -10.218 | 328.1 | 0.45 | -10.233 |
| | | | unfolded | 328.0 | 0.45 | -10.248 | | | |
| | | 1.2 | folded | 347.0 | 0.43 | -16.858 | 347.2 | 0.43 | -16.788 |
| | | | unfolded | 347.3 | 0.43 | -16.718 | | | |
| | | 1.3 | folded | 365.8 | 0.43 | $-\infty$ | 365.6 | 0.43 | $-\infty$ |
| | | | unfolded | 365.3 | 0.43 | $-\infty$ | | | |
| Ribonuclease H | 2RN2 | 1.0 | folded | 310.8 | 0.23 | -4.050 | 310.5 | 0.23 | -3.924 |
| | | | unfolded | 310.2 | 0.23 | -3.799 | | | |
| | | 1.1 | folded | 328.0 | 0.23 | -8.243 | 327.7 | 0.23 | -8.150 |
| | | | unfolded | 327.3 | 0.23 | -8.058 | | | |
| | | 1.2 | folded | 343.9 | 0.22 | -12.275 | 344.0 | 0.22 | -12.230 |
| | | | unfolded | 344.1 | 0.22 | -12.186 | | | |
| | | 1.3 | folded | 361.2 | 0.22 | -16.411 | 361.1 | 0.22 | -16.591 |
| | | | unfolded | 360.9 | 0.22 | -16.771 | | | |

Supplementary Table 4. The protein name, PDB ID, threshold of native contacts fraction $Q_{kin}$, protein folding rate constant $k_F$, protein folding lag time $t_0$, coefficient of determination $R^2$ for curve fitting, mean *in silico* protein folding time $\langle\tau_F^{sim}\rangle$, mean experimental protein folding time $\langle\tau_F^{exp}\rangle$ and the scaling factor $\alpha = \frac{\langle\tau_F^{exp}\rangle}{\langle\tau_F^{sim}\rangle}$ of training-set proteins

| Protein | PDB ID | $Q_{kin}$ | $k_F$ (ns$^{-1}$) | $t_0$ (ns) | $R^2$ | $\langle\tau_F^{sim}\rangle$ (ns) | $\langle\tau_F^{exp}\rangle$ (s) | $\frac{\langle\tau_F^{exp}\rangle}{\langle\tau_F^{sim}\rangle}$ |
|---|---|---|---|---|---|---|---|---|
| EC298 | 1RYK | 0.83 | 0.151 | 1.873 | 0.99813 | 8.51 | 0.00011 [b] | 12,926 |
| λ-repressor$_{6-85}$ | 1LMB | 0.68 | 0.098 | 2.601 | 0.99950 | 12.81 | 0.00003 [b] | 2,342 |
| bACBP | 2ABD | 0.80 | 0.172 | 2.639 | 0.99892 | 8.44 | 0.00095 [b] | 112,559 |
| IM7 | 1CEI | 0.64 | 0.055 | 1.844 | 0.99890 | 20.00 | 0.00075 [b] | 37,500 |
| Im9 | 1IMQ | 0.64 | 0.222 | 2.064 | 0.99840 | 6.57 | 0.00066 [b] | 100,457 |
| Cytochrome B562 | 256B | 0.75 | 0.053 | 3.045 | 0.99935 | 21.99 | 0.00125 [c] | 56,844 |
| ABP1 SH3 [a] | 1JO8 | 0.89 | 0.365 (0.527)<br>0.027 (0.473) | 1.187<br>1.635 | 0.99980 | 20.58 | 0.08547 [b] | 4,153,061 |
| Fyn SH3 | 1SHF | 0.75 | 0.218 | 1.871 | 0.99943 | 6.45 | 0.01060 [b] | 1,643,411 |
| CspB Bc | 1C9O | 0.52 | 0.099 | 1.705 | 0.99971 | 11.84 | 0.00073 [b] | 61,655 |
| CspA | 1MJC | 0.64 | 0.084 | 2.603 | 0.99956 | 14.51 | 0.00503 [d] | 346,657 |
| Tenascin [a] | 1TEN | 0.60 | 0.150 (0.940)<br>0.008 (0.060) | 2.262<br>24.845 | 0.99907 | 17.56 | 0.16611 [b] | 9,459,567 |
| Twitchin | 1WIT | 0.60 | 0.063 | 2.698 | 0.99968 | 18.62 | 0.66667 [b] | 35,803,974 |
| Hpr | 1POH | 0.68 | 0.026 | 9.184 | 0.99851 | 47.72 | 0.06711 [e] | 1,406,329 |
| Urm1 | 2QJL | 0.48 | 0.012 | 1.908 | 0.99973 | 82.93 | 0.07576 [b] | 913,542 |
| Src SH2 | 1SPR | 0.83 | 0.217 | 1.643 | 0.99868 | 6.26 | 0.00016 [b] | 25,559 |

| | | | | | | | | |
|---|---|---|---|---|---|---|---|---|
| Azurin (apo) | 1E65 | 0.78 | 0.008 | 12.400 | 0.99955 | 132.08 | 0.00735 [b] | 55,648 |
| CheY | 3CHY | 0.52 | 0.026 | 14.708 | 0.99848 | 53.04 | 0.02653 [f] | 500,189 |
| Ribonuclease H | 2RN2 | 0.40 | 0.022 | 12.110 | 0.99917 | 58.07 | 1.35135 [g] | 23,271,052 |
| | | | | | $\langle Q_{\text{kin}} \rangle = 0.66880,$ | | $\alpha$ = 4,331,293 | |

[a] Insufficiently sampled proteins. Their $k_F$ is shown as $k_i$ ($f_i$) and $t_0$ is shown as $t_i$, where $i$ = 1 or 2;

[b] Obtained from De Sancho et al.[11];

[c] Obtained from Wittung-Stafshede et al.[81];

[d] Obtained from Reid et al.[12];

[e] Obtained from Van Nuland et al.[82];

[f] Obtained from López-Hernéndez & Serrano[14];

[g] Obtained from Raschke & Marqusee[83];

Supplementary Table 5. The levels of $n_{\text{scal}}$ values used to parameterize an arbitrary protein

| Structural Class | Levels | | | | |
|---|---|---|---|---|---|
| | $\langle n^*_{\text{scal}} \rangle_{\text{class}}$ | $1.23 \times \langle n^*_{\text{scal}} \rangle_{\text{class}}$ | $1.46 \times \langle n^*_{\text{scal}} \rangle_{\text{class}}$ | $1.70 \times \langle n^*_{\text{scal}} \rangle_{\text{class}}$ | Overall highest |
| α | 1.1954 | 1.4704 | 1.7453 | 2.0322 | 2.5044 |
| β | 1.4732 | 1.8120 | 2.1508 | 2.5044 | 2.5044 |
| α/β | 1.1556 | 1.4213 | 1.6871 | 1.9644 | 2.5044 |
| Interface | 1.2747 | 1.5679 | 1.8611 | 2.1670 | 2.5044 |

Supplementary Table 6. Interaction site types and nonbonding parameters of the CG ribosome model

| | Chemical group | CG interaction site name | CG interaction site type | $R_i$ (Å) |
|---|---|---|---|---|
| rRNA and tRNA | Phosphate | P | P | 6.447660 |
| | Ribose | R | R | 5.231399 |
| | Adenine, Guanine | PU1 | BR [a] | 5.342436 |
| | | PU2 | | |
| | Uracil, Cytosine | PY | | |
| Ribosomal protein | ALA | SA | SA | 2.862278 |
| | CYS | SC | SC | 3.030648 |
| | ASP | SD | SD | 3.142894 |
| | GLU | SE | SE | 3.367386 |
| | PHE | SF | SF | 3.535755 |
| | GLY | SG | SG | 2.525540 |
| | HIS | SH | SH | 3.423509 |
| | ILE | SI | SI | 3.423509 |
| | LYS | SK | SK | 3.535755 |
| | LEU | SL | SL | 3.423509 |
| | MET | SM | SM | 3.423509 |
| | ASN | SN | SN | 3.199017 |
| | PRO | SP | SP | 3.086771 |
| | GLN | SQ | SQ | 3.423509 |
| | ARG | SR | SR | 3.704125 |
| | SER | SS | SS | 2.918401 |
| | THR | ST | ST | 3.142894 |
| | VAL | SV | SV | 3.311263 |
| | TRP | SW | SW | 3.816371 |
| | TYR | SY | SY | 3.591879 |

[a] Four different basic groups have the same CG interaction site type BR located at the centroid of each conjugated ring.

Supplementary Table 7. Equilibrium bond lengths, angles and dihedrals used for A-site and P-site NC-tRNA (or aa-tRNA)

| PDBid | Resolution (Å) | A-site tRNA | | | | P-site tRNA | | | |
|---|---|---|---|---|---|---|---|---|---|
| | | $b_0$ (CA-R) (Å) | $\theta_0$ (CA-R-PU2) (degree) | $\theta_0$ (CA-R-P) (degree) | $\varphi_0$ (CA-R-P-PU2) (degree) | $b_0$ (CA-R) (Å) | $\theta_0$ (CA-R-PU2) (degree) | $\theta_0$ (CA-R-P) (degree) | $\varphi_0$ (CA-R-P-PU2) (degree) |
| 5nwy | 2.9 | – | – | – | – | 4.66 | 143.32 | 99.51 | -170.62 |
| 5jte | 3.6 | 4.63 | 126.72 | 106.40 | 126.66 | 4.79 | 139.44 | 124.06 | -173.92 |
| 6i0y | 3.2 | – | – | – | – | 4.73 | 111.83 | 133.63 | -152.69 |
| 3jbu | 3.64 | – | – | – | – | 5.02 | 129.39 | 97.48 | -145.81 |
| 4uy8 | 3.8 | – | – | – | – | 4.93 | 111.98 | 134.20 | -154.60 |
| 6enj | 3.7 | 3.91 | 127.52 | 104.99 | 129.18 | – | – | – | – |
| 6enu | 3.1 | – | – | – | – | 4.43 | 136.54 | 109.20 | -163.82 |
| 5ju8 | 3.6 | – | – | – | – | 4.74 | 139.31 | 119.14 | -165.53 |
| Average | | 4.27 | 127 | 106 | 128 | 4.76 | 130 | 117 | -161 |

Supplementary Table 8. Original and rescaled $\langle\tau_{\text{trans}}^{\text{intrinsic}}\rangle_i$ , $\langle\tau_{\text{PT}}^{\text{intrinsic}}\rangle$ and $\langle\tau_{\text{TL}}^{\text{intrinsic}}\rangle$ data, as well as corresponding normalized codon usage frequencies $w_i$

| | *E. coli* (at 310 K) | | | | | | | |
|---|---|---|---|---|---|---|---|---|
| | Codon | Fluitt & Viljoen's data[30] (s) | $w_i$ [31] | Rescaled data (s) | Codon | Fluitt & Viljoen's data[30] (s) | $w_i$ [31] | Rescaled data (s) |
| $\langle\tau_{\text{trans}}^{\text{intrinsic}}\rangle_i$ | UUU | 0.136 | 0.022499 | 0.068164 | GUU | 0.026 | 0.018409 | 0.013031 |
| | UUC | 0.195 | 0.016260 | 0.097735 | GUC | 0.208 | 0.015070 | 0.104251 |
| | UUG | 0.050 | 0.013430 | 0.025060 | GUG | 0.042 | 0.025879 | 0.021051 |
| | UUA | 0.157 | 0.013880 | 0.078689 | GUA | 0.073 | 0.010980 | 0.036588 |
| | UCU | 0.055 | 0.008660 | 0.027566 | GCU | 0.039 | 0.015530 | 0.019547 |
| | UCC | 0.246 | 0.008840 | 0.123297 | GCC | 0.415 | 0.025409 | 0.208000 |
| | UCG | 0.096 | 0.008810 | 0.048116 | GCG | 0.044 | 0.032749 | 0.022053 |
| | UCA | 0.106 | 0.007640 | 0.053128 | GCA | 0.083 | 0.020579 | 0.041600 |
| | UGU | 0.075 | 0.005260 | 0.037590 | GGU | 0.035 | 0.024439 | 0.017542 |
| | UGC | 0.109 | 0.006400 | 0.054631 | GGC | 0.049 | 0.028619 | 0.024559 |
| | UGG | 0.168 | 0.015170 | 0.084203 | GGG | 0.081 | 0.011270 | 0.040598 |
| | UGA | 0.012 | 0.001040 | 0.006014 | GGA | 0.324 | 0.008440 | 0.162391 |
| | UAU | 0.053 | 0.016380 | 0.026564 | GAU | 0.077 | 0.032319 | 0.038593 |
| | UAC | 0.077 | 0.012160 | 0.038593 | GAC | 0.116 | 0.019109 | 0.058140 |
| | UAG | 0.019 | 0.000250 | 0.009523 | GAG | 0.036 | 0.018199 | 0.018043 |
| | UAA | 0.011 | 0.002060 | 0.005513 | GAA | 0.057 | 0.039419 | 0.028569 |
| | CUU | 0.260 | 0.011470 | 0.130314 | AUU | 0.097 | 0.030229 | 0.048617 |
| | CUC | 0.204 | 0.010930 | 0.102246 | AUC | 0.128 | 0.024589 | 0.064154 |
| | CUG | 0.035 | 0.051888 | 0.017542 | AUG | 0.266 | 0.027589 | 0.133321 |
| | CUA | 0.286 | 0.003930 | 0.143345 | AUA | 0.128 | 0.004910 | 0.064154 |
| | CCU | 0.143 | 0.007250 | 0.071672 | ACU | 0.055 | 0.009040 | 0.027566 |
| | CCC | 0.197 | 0.005580 | 0.098738 | ACC | 0.153 | 0.022869 | 0.076685 |
| | CCG | 0.134 | 0.022659 | 0.067162 | ACG | 0.129 | 0.014470 | 0.064656 |
| | CCA | 0.237 | 0.008480 | 0.118786 | ACA | 0.178 | 0.007670 | 0.089215 |
| | CGU | 0.028 | 0.020599 | 0.014034 | AGU | 0.085 | 0.009080 | 0.042603 |
| | CGC | 0.035 | 0.021459 | 0.017542 | AGC | 0.127 | 0.015900 | 0.063653 |
| | CGG | 0.397 | 0.005700 | 0.198979 | AGG | 0.461 | 0.001510 | 0.231056 |
| | CGA | 0.034 | 0.003700 | 0.017041 | AGA | 0.190 | 0.002470 | 0.095229 |
| | CAU | 0.296 | 0.012850 | 0.148357 | AAU | 0.109 | 0.018269 | 0.054631 |
| | CAC | 0.222 | 0.009460 | 0.111268 | AAC | 0.161 | 0.021479 | 0.080694 |
| | CAG | 0.231 | 0.029139 | 0.115779 | AAG | 0.102 | 0.010700 | 0.051123 |
| | CAA | 0.179 | 0.015100 | 0.089716 | AAA | 0.076 | 0.033879 | 0.038092 |
| $\langle\tau_{\text{PT}}^{\text{intrinsic}}\rangle$ | | 0.000679 | – | 0.000340 | | | | |
| $\langle\tau_{\text{TL}}^{\text{intrinsic}}\rangle$ | | 0.008381 | – | 0.004201 | | | | |

Supplementary Table 9. The structural information and the parameterization information of the initial 14 *E. coli* enzymes for screening. (* denotes the instable domain or interface that cannot be stabilized using the highest level of $n_{\text{scal}}$ and the median value 1.8611 was used.)

| PDB ID | Name | Length | Domain and Interfaces | Structural Class | Minimum $n^{*}_{\text{scal}}$ required to keep kinetic stability |
|---|---|---|---|---|---|
| 1AKE | Adenylate kinase | 214 | Domain 1: 1 - 214 | α/β | 1.1556 |
| 1AQ2 | Phosphoenolpyruvate carboxykinase | 540 | Domain 1: 1 - 42; 66 - 227 | α/β | 1.1556 |
| | | | Domain 2: 43 - 65; 284 - 342 | β | 1.4732 |
| | | | Domain 3: 228 - 283; 343 - 540 | α/β | 1.1556 |
| | | | 1 \| 2 Interface | – | 1.2747 |
| | | | 1 \| 3 Interface | – | 1.2747 |
| | | | 2 \| 3 Interface | – | 1.2747 |
| 1C2T | Glycinamide ribonucleotide transformylase | 212 | Domain 1: 1 - 212 | α/β | 1.4213 |
| 1FDR | Flavodoxin reductase | 248 | Domain 1: 1 - 96 | β | 1.4732 |
| | | | Domain 1: 97 - 248 | α/β | 1.1556 |
| | | | 1 \| 2 Interface | – | 1.5679 |
| 1PDA | Porphobilinogen deaminase | 313 | Domain 1: 1 - 99; 200 - 220 | α/β | 1.1556 |
| | | | Domain 2: 100 - 199 | α/β | 1.1556 |
| | | | Domain 3: 221 - 313 | α/β | 1.4213 |
| | | | 1 \| 2 Interface | – | 1.5679 |
| | | | 1 \| 3 Interface | – | 1.2747 |
| | | | 2 \| 3 Interface | – | 1.5679 |
| 1VHL | Dephospho-CoA kinase | 206 | Domain 1: 1 - 206 | α/β | 1.1556 |
| 2FMT | Methionyl-tRNA formyltransferase | 315 | Domain 1: 0 - 208 | α/β | 1.4213 |
| | | | Domain 2: 209 - 314 | α/β | 1.6871 |
| | | | 1 \| 2 Interface | – | 1.8611 |
| 2OFP | Ketopantoate reductase | 303 | Domain 1: 1 - 167 | α/β | 1.1556 |
| | | | Domain 2: 168 - 303 | α | 1.1954 |
| | | | 1 \| 2 Interface | – | 1.8611 * |
| 2ZCV | Quinone oxidoreductase 2 | 303 | Domain 1: 1~136; 171~193; 259~267 | α/β | 1.1556 |
| | | | Domain 2: 137~170; 194~258; 268~303 | α/β | 1.9644 |
| | | | 1 \| 2 Interface | – | 2.167 |
| 3CW7 | DNA-3-methyladenine glycosylase 2 | 282 | Domain 1: 1 - 112 | α/β | 1.1556 |
| | | | Domain 2: 113 - 230 | α/β | 1.1556 |
| | | | Domain 3: 231 - 282 | α | 1.1954 |

| | | | | | |
|---|---|---|---|---|---|
| | | | 1 \| 2 Interface | – | 1.8611 * |
| | | | 1 \| 3 Interface | – | 1.2747 |
| | | | 2 \| 3 Interface | – | 1.2747 |
| 3K6L | Peptide deformylase | 169 | Domain 1: 0 - 168 | α/β | 1.4213 |
| 3LBF | Protein-L-isoaspartate O-methyltransferase | 208 | Domain 1: 1 - 208 | α/β | 1.1556 |
| 4C5C | D-alanine--D-alanine ligase B | 306 | Domain 1: 1 - 84 | α/β | 1.1556 |
| | | | Domain 2: 85 - 110; 184 - 306 | α/β | 1.1556 |
| | | | Domain 3: 111 - 183 | α/β | 1.1556 |
| | | | 1 \| 2 Interface | – | 2.1670 |
| | | | 1 \| 3 Interface | – | 1.2747 |
| | | | 2 \| 3 Interface | – | 2.1670 |
| 4KJK | Dihydrofolate reductase | 159 | Domain 1: 1-159 | α/β | 1.4213 |

Supplementary Table 10. Parameterization results for CAT-III

| REX input and output parameters | | | | | Average | Linear regression parameters | | | |
|---|---|---|---|---|---|---|---|---|---|
| $n_{\mathrm{scal}}$ | Starting state | $T_{\mathrm{m}}$ (K) | $Q_{\mathrm{eq}}$ | $\Delta G_{\mathrm{UN}}^{\mathrm{sim}}(T^{\mathrm{exp}})$ (kcal/mol) | $\Delta G_{\mathrm{UN}}^{\mathrm{sim}}(T^{\mathrm{exp}})$ (kcal/mol) | $m$ | $b$ | $R^2$ | $n_{\mathrm{scal}}^{*}$ |
| 1.0 | folded | 323.5 | 0.67 | -9.027 | -6.910 | -14.688 | 8.006 | 0.9658 | 1.2471 |
| | unfolded | 303.5 | 0.55 | -4.792 | | | | | |
| 1.1 | folded | 343.2 | 0.71 | -9.700 | -7.695 | | | | |
| | unfolded | 322.8 | 0.77 | -5.691 | | | | | |
| 1.2 | folded | 361.2 | 0.71 | -12.438 | -9.847 | | | | |
| | unfolded | 340.8 | 0.78 | -7.256 | | | | | |

Supplementary Table 11. Experimentally measured rate constant ($k_{cat}$)[a] and Michaelis constant ($K_M$)[b] of the enzymatic reaction catalyzed by the fast and slow DDLB variants and the corresponding relative specific activity $k_{cat}^{slow}/k_{cat}^{fast}$.

| Biological replicates | $k_{cat}^{slow}$ (min$^{-1}$) | $K_M^{slow}$ (μM) | $k_{cat}^{fast}$ (min$^{-1}$) | $K_M^{fast}$ (μM) | $k_{cat}^{slow}/k_{cat}^{fast}$ |
|---|---|---|---|---|---|
| 1 | 986.05 ± 16.61 | 1181.26 ± 79.81 | 1045.70 ± 18.22 | 1759.32 ± 77.84 | 0.9430 |
| 2 | 895.40 ± 18.58 | 2272.66 ± 114.37 | 1125.37 ± 20.68 | 2414.56 ± 105.02 | 0.7956 |
| 3 | 1083.42 ± 16.94 | 2163.70 ± 80.36 | 1088.12 ± 24.36 | 2570.70 ± 133.06 | 0.9957 |
| 4 | 771.40 ± 9.00 | 1517.97 ± 44.98 | 958.48 ± 16.65 | 1633.22 ± 75.38 | 0.8048 |
| 5 | 601.89 ± 9.84 | 1568.39 ± 78.38 | 698.28 ± 13.87 | 2082.02 ± 118.30 | 0.8619 |
| $\langle k_{cat}^{slow}/k_{cat}^{fast}\rangle$[c] | 0.8802, 95%CI [0.8126, 0.9478] | | | | |

[a] $k_{cat}$ is presented as mean ± standard deviation errors of the curve fitting;

[b] $K_M$ is presented as mean ± standard deviation errors of the curve fitting;

[c] $\langle k_{cat}^{slow}/k_{cat}^{fast}\rangle$ is statistically less than 1 with a one-tailed t-test $p$-value of 0.0186.

Supplementary Table 12. Experimentally measured rate constant ($k_{cat}$)[a] of the enzymatic reaction catalyzed by the fast and slow DHFR variants and the corresponding relative specific activity $k_{cat}^{fast}/k_{cat}^{slow}$.

| Biological replicates | $k_{cat}^{slow}$ ($min^{-1}$) | $k_{cat}^{fast}$ ($min^{-1}$) | $k_{cat}^{fast}/k_{cat}^{slow}$ |
|---|---|---|---|
| 1 | 442.74 ± 21.46 | 220.25 ± 17.45 | 0.4975 |
| 2 | 301.75 ± 30.46 | 332.64 ± 26.29 | 1.1024 |
| 3 | 374.86 ± 21.71 | 309.66 ± 20.81 | 0.8261 |
| 4 | 425.21 ± 43.06 | 356.88 ± 44.13 | 0.8393 |
| 5 | 365.64 ± 14.51 | 467.94 ± 26.49 | 1.2798 |
| $\langle k_{cat}^{fast}/k_{cat}^{slow} \rangle$[b] | 0.9089, 95%CI [0.6838, 1.1539] | | |

[a] $k_{cat}$ is presented as mean ± standard deviation errors of the technical replicates;

[b] $\langle k_{cat}^{fast}/k_{cat}^{slow} \rangle$ is statistically indistinguishable from 1 with a two-tailed t-test *p*-value of 0.5323.

Supplementary Table 13. Limited-proteolysis mass spectrometry results for DDLB. The statistically significant peptides ($log_2$(effect size) > 1 or < -1, and $-log_{10}$(p-value) > 2) are highlighted in pink.

| Time | PK site / full-tryptic peptide[a] | Effect size ($log_2$)[b] | p-value ($-log_{10}$)[c] | Coefficients of variation[d] |
|---|---|---|---|---|
| 1 min | 44-51 | 0.09 | 0.21 | 0.1259 |
| | 100-119 | -0.84 | 0.50 | 0.1315 |
| | 130-147 | -0.40 | 0.82 | 0.0688 |
| | 32-43 | 0.75 | 1.01 | 0.1305 |
| | 2 | -0.51 | 1.11 | 0.1996 |
| | 138 | 0.72 | 1.68 | 0.0367 |
| | 194 | 0.85 | 1.85 | 0.0557 |
| | 102-119 | -0.83 | 1.90 | 0.0519 |
| | 203-215 | 0.97 | 1.95 | 0.0754 |
| | 140 | 0.81 | 2.12 | 0.0889 |
| | 17-31 | -0.69 | 2.32 | 0.1200 |
| | 2-9 | 1.01 | 2.52 | 0.0489 |
| | 157-168 | 0.51 | 2.68 | 0.0277 |
| | 192 | 1.41 | 4.70 | 0.0467 |
| 5 min | 104 | 0.06 | 0.03 | 0.4064 |
| | 102-119 | -0.18 | 0.04 | 0.1152 |
| | 100-119 | -0.77 | 0.41 | 0.0230 |
| | 130-147 | 0.41 | 0.42 | 0.3156 |
| | 44-51 | 0.13 | 0.48 | 0.0189 |
| | 17-31 | -0.49 | 0.94 | 0.2549 |
| | 138 | 0.70 | 1.37 | 0.1456 |
| | 32-43 | 0.86 | 1.59 | 0.0877 |
| | 194 | 0.90 | 1.62 | 0.1427 |
| | 2-9 | 0.85 | 1.63 | 0.1349 |
| | 191 | 1.18 | 1.85 | 0.1167 |
| | 157-168 | 0.34 | 1.96 | 0.0410 |
| | 289-300 | 9.38 | 1.96 | 0.1485 |
| | 140 | 0.84 | 2.22 | 0.0846 |
| | 192 | 1.57 | 5.01 | 0.0458 |
| 2 h | 100-119 | -0.27 | 0.11 | 0.1242 |
| | 104 | 0.55 | 0.31 | 0.4894 |
| | 44-51 | 0.09 | 0.33 | 0.0405 |
| | 130-147 | -0.22 | 0.47 | 0.1022 |
| | 136 | 0.71 | 0.80 | 0.2527 |
| | 102-119 | -0.36 | 0.86 | 0.0906 |
| | 32-43 | 0.75 | 0.89 | 0.1473 |
| | 17-31 | -0.33 | 0.91 | 0.0788 |
| | 194 | 0.80 | 1.06 | 0.2122 |
| | 138 | 0.75 | 1.44 | 0.1431 |
| | 140 | 0.88 | 1.64 | 0.1354 |
| | 157-168 | 0.43 | 1.88 | 0.0807 |
| | 191 | 1.15 | 1.99 | 0.1056 |
| | 192 | 1.44 | 4.22 | 0.0736 |

[a] Residue number of the PK site is shown for a half-tryptic peptide, while the residue range is presented for a full-tryptic peptide;

[b] Effect size is calculated by $\log_2\left(\frac{\text{Abundance in refolded samples}}{\text{Abundance in native samples}}\right)$. A positive effect size for a half-tryptic peptide and a negative effect-size for a full-tryptic peptide both indicate that the site is more exposed in the refolded form of the protein (vice versa);

[c] p-value is obtained from Welch's t-test;

[d] Coefficient of variation (standard deviation relative to mean) of peptide abundance in three (n=3) independent refolding reactions on three biological replicates.

Supplementary Table 14. Limited-proteolysis mass spectrometry results for DHFR. The statistically significant peptides ($log_2$(effect size) > 1 or < -1, and $-log_{10}$(p-value) > 2) are highlighted in pink.

| Time | PK site / full-tryptic peptide[a] | Effect size ($log_2$)[b] | p-value ($-log_{10}$)[c] | Coefficients of variation[d] |
|---|---|---|---|---|
| 1 min | 152 | 0.26 | 0.41 | 0.2118 |
| | 85 | 0.19 | 0.42 | 0.1199 |
| | 59-71 | -0.62 | 0.89 | 0.1112 |
| | 45-57 | -0.62 | 1.25 | 0.2211 |
| | 59-76 | -0.75 | 1.26 | 0.1090 |
| | 77-98 | -0.72 | 1.33 | 0.0644 |
| | 23 | 9.58 | 1.38 | 0.2979 |
| | 99-106 | -0.58 | 1.68 | 0.0687 |
| | 45-58 | -13.09 | 1.98 | 0.0000 |
| 5 min | 59-71 | -0.22 | 0.10 | 0.3849 |
| | 34-44 | 0.25 | 0.18 | 0.2016 |
| | 45-57 | -0.25 | 0.23 | 0.2435 |
| | 85 | 0.35 | 0.51 | 0.2252 |
| | 23 | 2.34 | 0.94 | 0.4224 |
| | 59-76 | -1.04 | 1.61 | 0.1692 |
| | 45-58 | -13.09 | 1.97 | 0.0000 |
| | 99-106 | -0.65 | 2.14 | 0.1580 |
| | 77-98 | -1.10 | 2.15 | 0.1091 |
| 2 h | 85 | -0.07 | 0.34 | 0.0696 |
| | 152 | 0.30 | 0.63 | 0.1593 |
| | 59-71 | -0.55 | 0.86 | 0.1576 |
| | 34-44 | 0.55 | 0.93 | 0.0407 |
| | 77-98 | -0.58 | 0.94 | 0.1789 |
| | 45-58 | -0.93 | 1.12 | 0.1000 |
| | 59-76 | -0.82 | 1.28 | 0.0613 |
| | 23 | 1.81 | 1.28 | 0.2705 |
| | 99-106 | -0.81 | 1.89 | 0.2495 |

[a] Residue number of the PK site is shown for a half-tryptic peptide, while the residue range is presented for a full-tryptic peptide;

[b] Effect size is calculated by $log_2\left(\frac{\text{Abundance in refolded samples}}{\text{Abundance in native samples}}\right)$. A positive effect size for a half-tryptic peptide and a negative effect-size for a full-tryptic peptide both indicate that peptide is more exposed in refolded protein (vice versa);

[c] p-value is obtained from Welch's t-test;

[d] Coefficient of variation (standard deviation relative to mean) of peptide abundance in three (n=3) independent refolding reactions on three biological replicates.

### Supplementary Figures

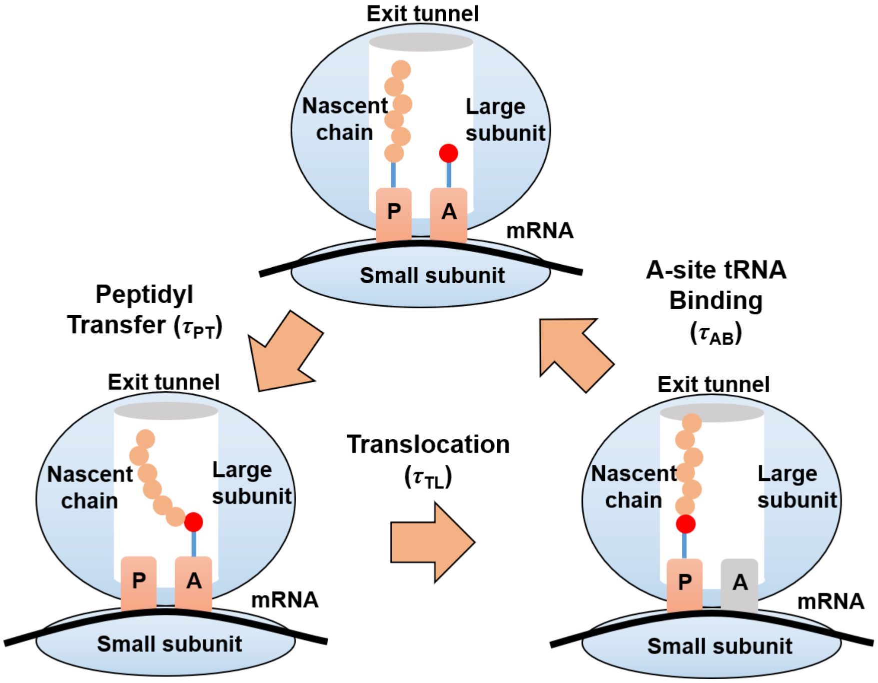

**Supplementary Figure 1. Scheme diagram of the on-ribosome continuous synthesis process.** The on-ribosome continuous synthesis process is simplified into three steps. First, aa-tRNA binds to the A-site. Then, the $NC_l$-tRNA at the P-site and the aa-tRNA at the A-site undergo the peptidyl transfer reaction catalyzed by the PTC and, forming $NC_{l+1}$-tRNA at the A-site. Finally, the P-site tRNA is translocated to the E-site and the A-site tRNA is translocated to the P-site, resulting in an empty A-site waiting proper aa-tRNA selection. The ribosome is shown in blue; the $NC_l$-tRNA, $NC_{l+1}$-tRNA and aa-tRNA are shown in orange for NC, pink for tRNA and red for the new amino acid; A-site and P-site are marked as “A” and “P”, respectively; the empty A-site is shown in grey; the exit tunnel is shown in white and the mRNA is shown in black. The dwell times before peptidyl transfer, translocation and A-site tRNA binding are denoted as $\tau_{PT}$, $\tau_{TL}$ and $\tau_{AB}$, respectively.

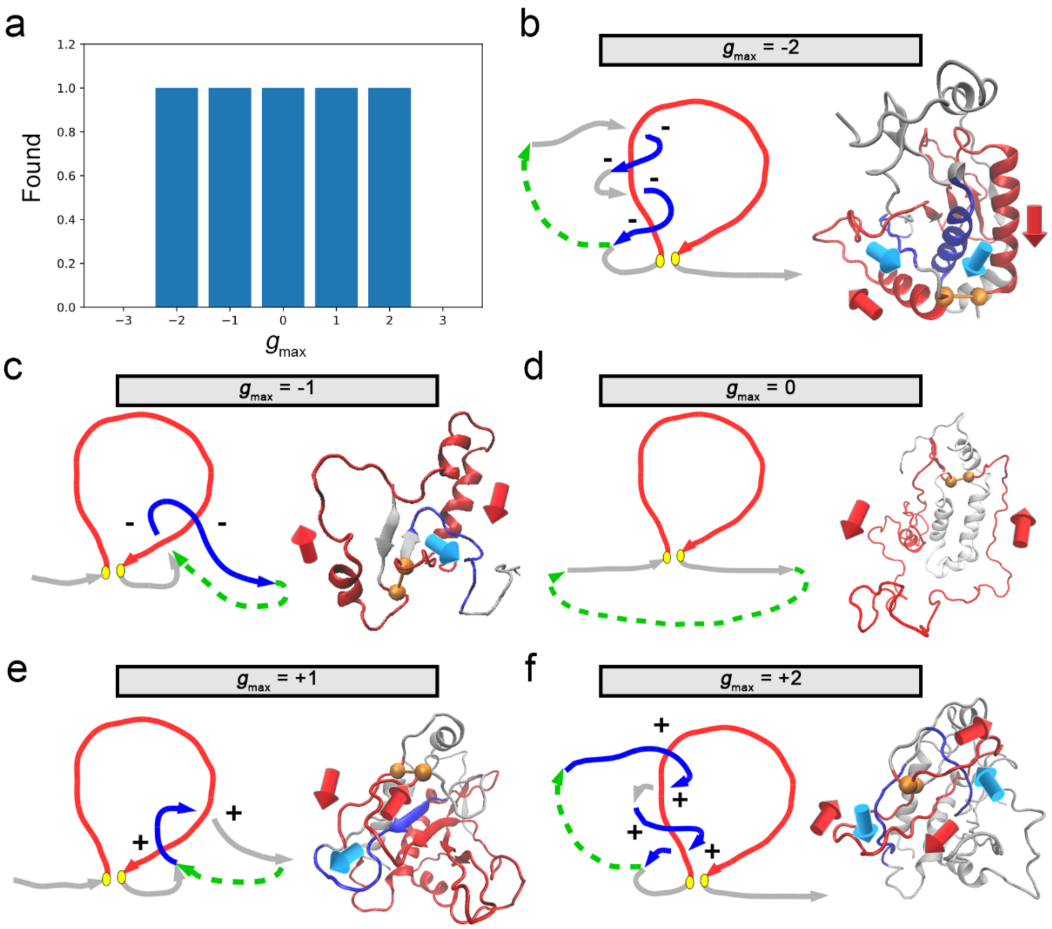

**Supplementary Figure 2. Test case of *g* for detecting topological entanglements.** a, The observed maximum *g* values ($g_{max}$) in the post-translational trajectories of the fast CAT-**III** variant (Found = 1 means the $g_{max}$ value is sampled in the trajectory; otherwise it is not sampled). The link diagrams (left) and the representative structures (right) are shown for each $g_{max}$ values in panel b, c, d, e and f. The closed loop formed by the native contact (yellow balls) are in red and the tail curves that form entanglements are in blue. The rest regions are shown in gray. The directions of those curves (from N-ter to C-ter) are marked in both the link diagrams and the representative structures. The +/- signs that are used to count the linking numbers are labelled near the curve intersections in the link diagrams. The imaginary curves that close the tail curves are shown in green dashed arrows.

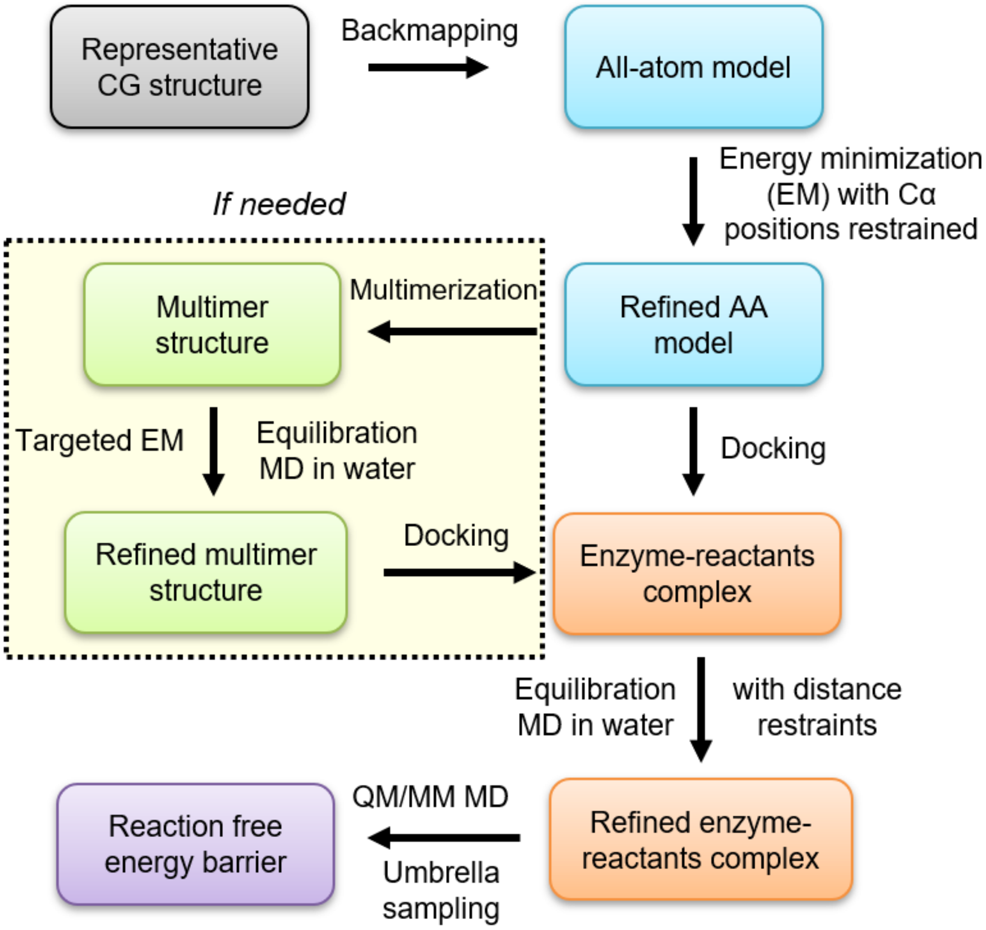

**Supplementary Figure 3. Scheme diagram of the estimation workflow for activation free energy barrier height.** The main workflow for estimating activation free energy barrier height for a representative coarse-grained (CG) structure is shown here. More details are described in Supplementary Methods Section 14.3.

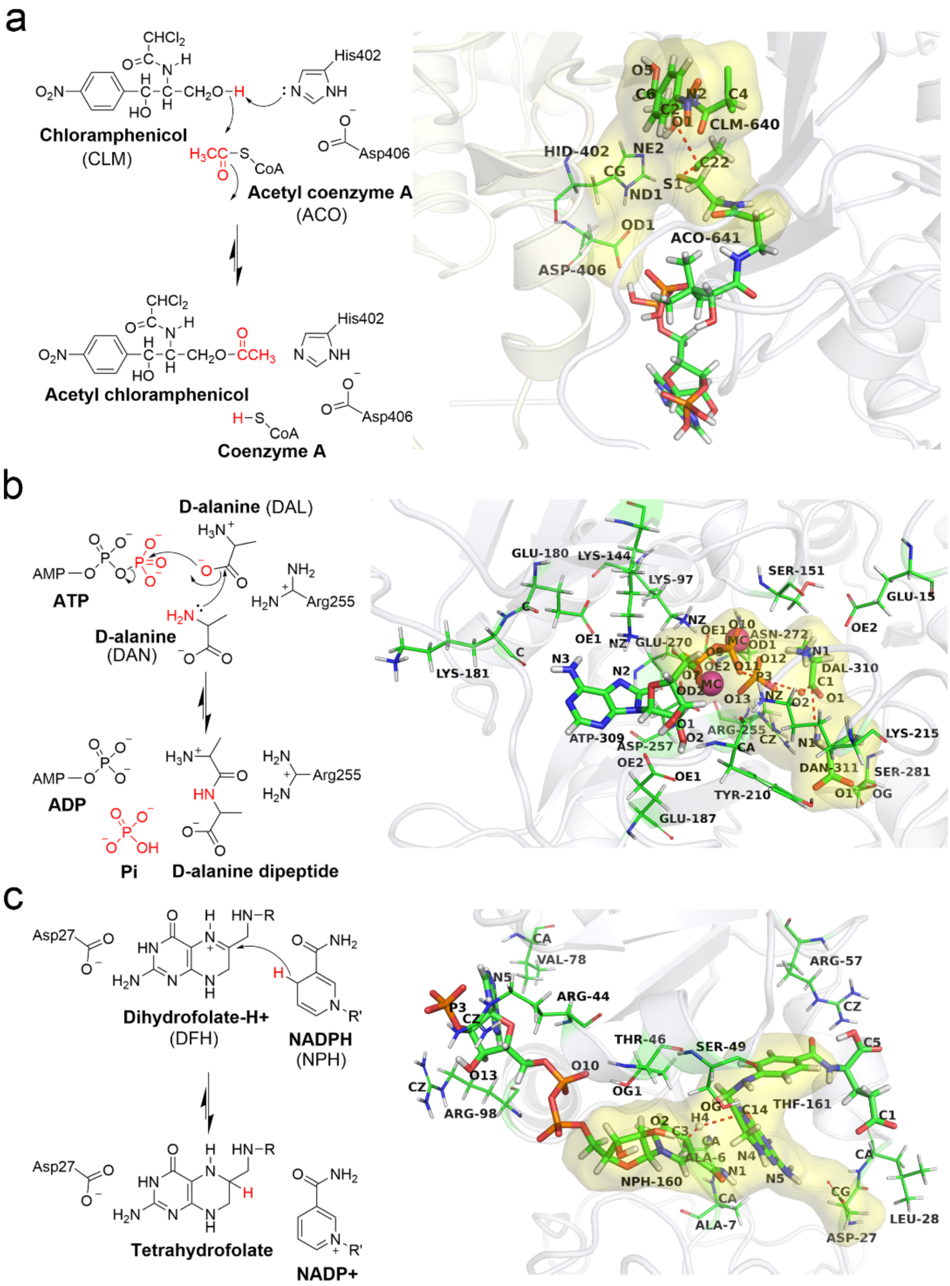

**Supplementary Figure 4. Reaction mechanisms and the ligand binding poses for candidate enzymes used in this study.** a, Reaction mechanism[66] (left) and ligand binding pose (right) of CAT-III; b, Reaction mechanism[68] (left) and ligand binding pose (right) of DDLB; c, Reaction mechanism[69] (left) and ligand binding pose (right) of DHFR. In reaction schemes, the moving groups are highlighted in red. In ligand binding pose diagrams, ligands are shown as bold sticks, whereas the amino acid residues that took part in the distance restraints are shown as thin sticks. The reaction coordinates are presented as red dotted lines. The QM regions are covered by yellow areas. The proteins are shown as Cartoon representation and different chains are shown in different colors.

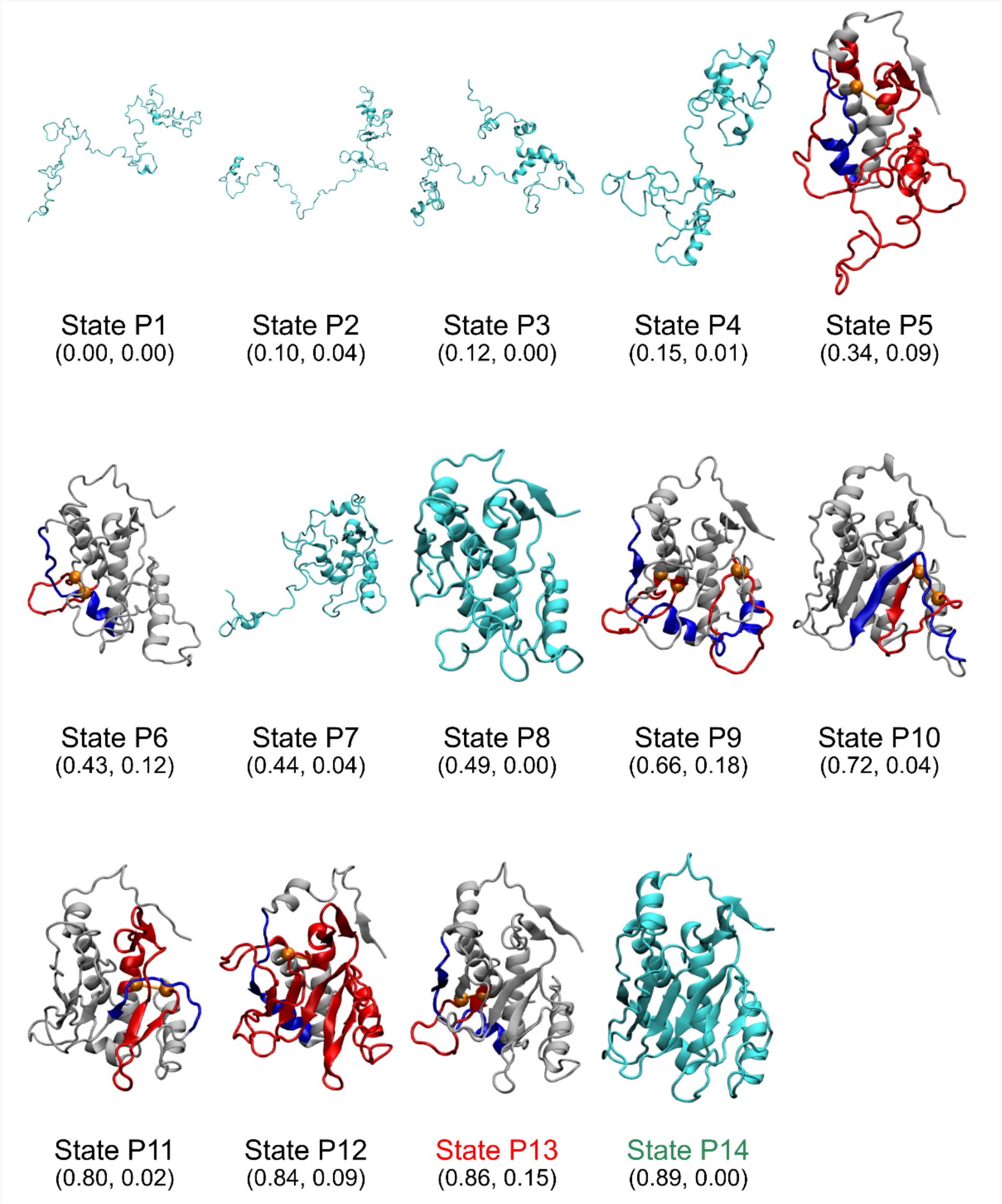

**Supplementary Figure 5. Representative structures of the metastable states of monomer CAT-III post-translational folding process.** The structures are represented with cartoon models. The ordinary intermediate structures, as well as the native structure, are shown in cyan. The entangled structures are shown in white, where the closed loop formed by the native contact (shown as two orange balls) is highlighted in red and the segment threading the red loop is highlighted in blue. The metastable name and the ($Q_{act}$, $G$) coordinates of the most probable cluster (microstate) in that metastable state are shown below each structure. The name of the near-native-like kinetics traps are highlighted in red and the native state is highlighted in green.

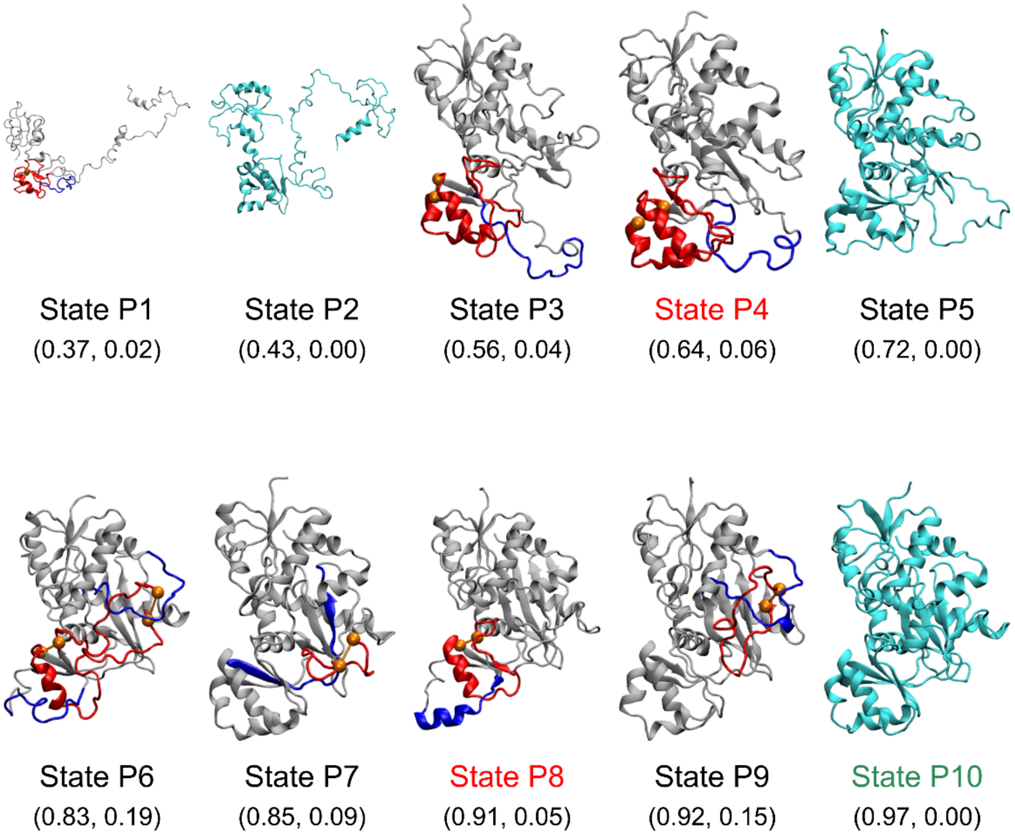

**Supplementary Figure 6. Representative structures of the metastable states of DDLB post-translational folding process.** The structures are represented with cartoon models. The ordinary intermediate structures, as well as the native structure, are shown in cyan. The entangled structures are shown in white, where the closed loop formed by the native contact (shown as two orange balls) is highlighted in red and the segment threading the red loop is highlighted in blue. The state name and the ($Q_{act}$, $G$) coordinates of the most probable cluster (microstate) in that metastable state are shown below each structure. The name of the near-native-like kinetics traps are highlighted in red and the native state is highlighted in green.

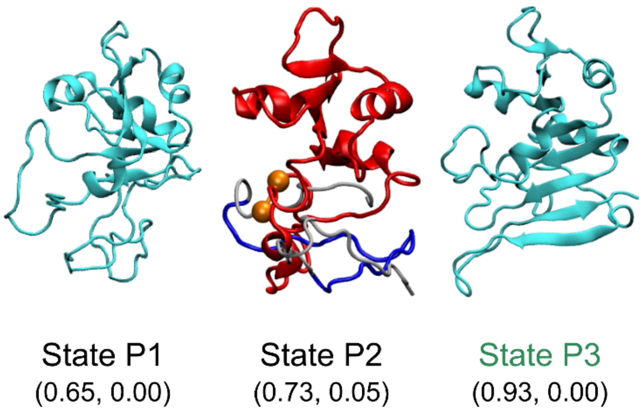

**Supplementary Figure 7. Representative structures of the metastable states of DHFR post-translational folding process.** The structures are represented with cartoon models. The ordinary intermediate structures, as well as the native structure, are shown in cyan. The entangled structures are shown in white, where the closed loop formed by the native contact (shown as two orange balls) is highlighted in red and the segment threading the red loop is highlighted in blue. The state name and the ($Q_{act}$, $G$) coordinates of the most probable cluster (microstate) in that metastable state are shown below each structure. The name of the native state is highlighted in green. No near-native-like kinetic traps were detected for DHFR.

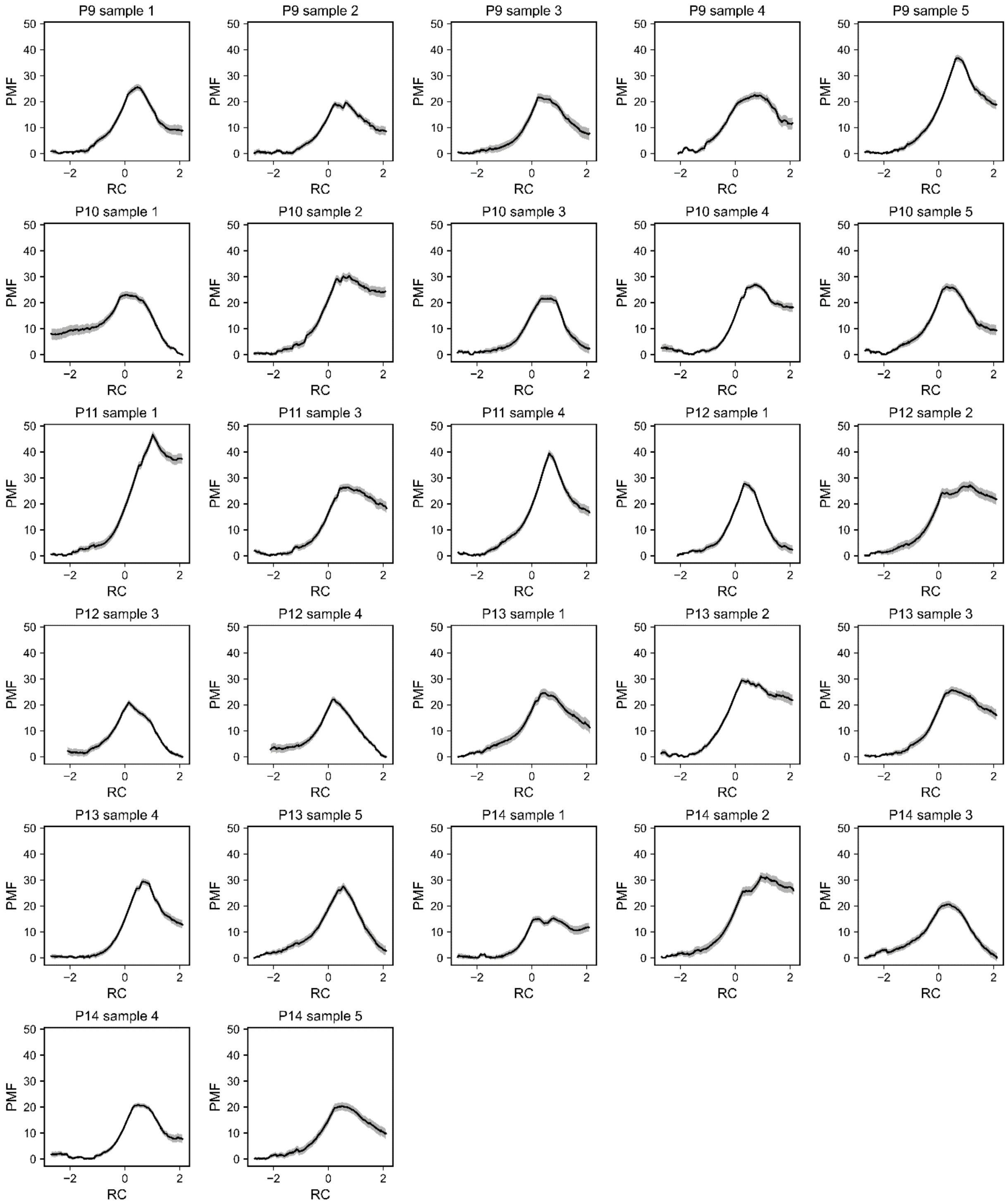

**Supplementary Figure 8. Potential of mean force (PMF) curves as a function of the reaction coordinate (RC) for the samples of CAT-III.** Only the samples that have successful substrates docking and QM/MM simulations are presented in this figure. Error bars (95% CIs) are shown as gray stripes.

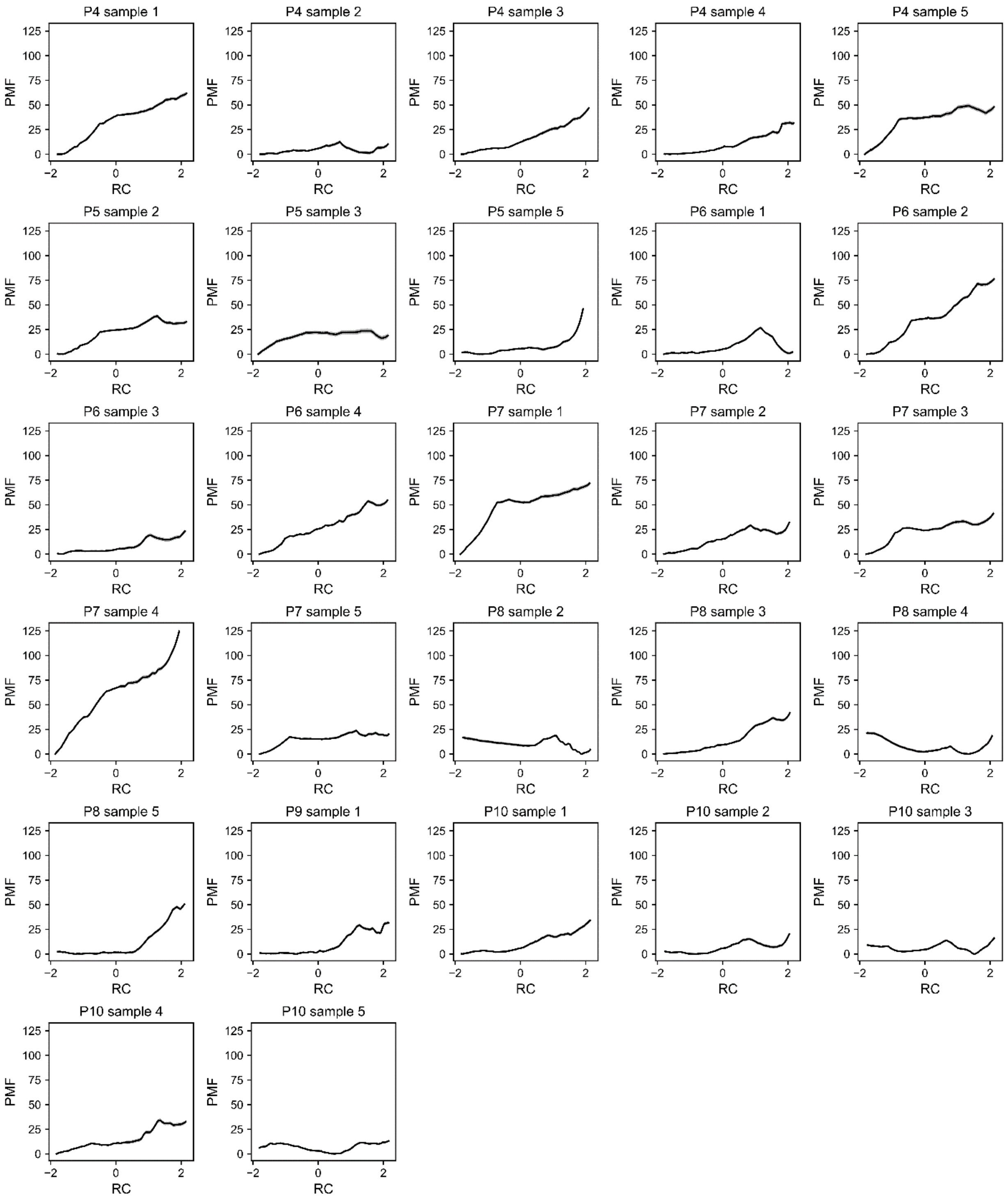

**Supplementary Figure 9. Potential of mean force (PMF) curves as a function of the reaction coordinate (RC) for the samples of DDLB.** Only the samples that have successful substrates docking and QM/MM simulations are presented in this figure. Error bars (95% CIs) are shown as gray stripes.

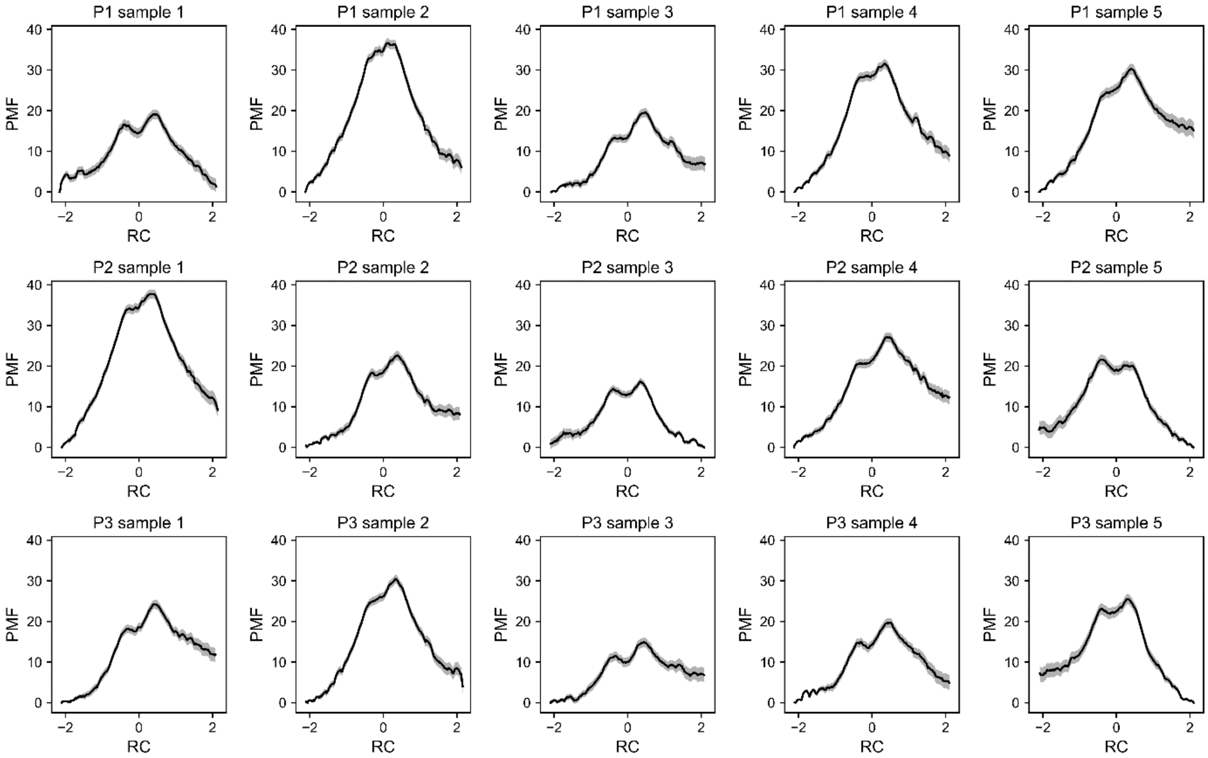

**Supplementary Figure 10. Potential of mean force (PMF) curves as a function of the reaction coordinate (RC) for the samples of DHFR.** Only the samples that have successful substrates docking and QM/MM simulations are presented in this figure. Error bars (95% CIs) are shown as gray stripes.

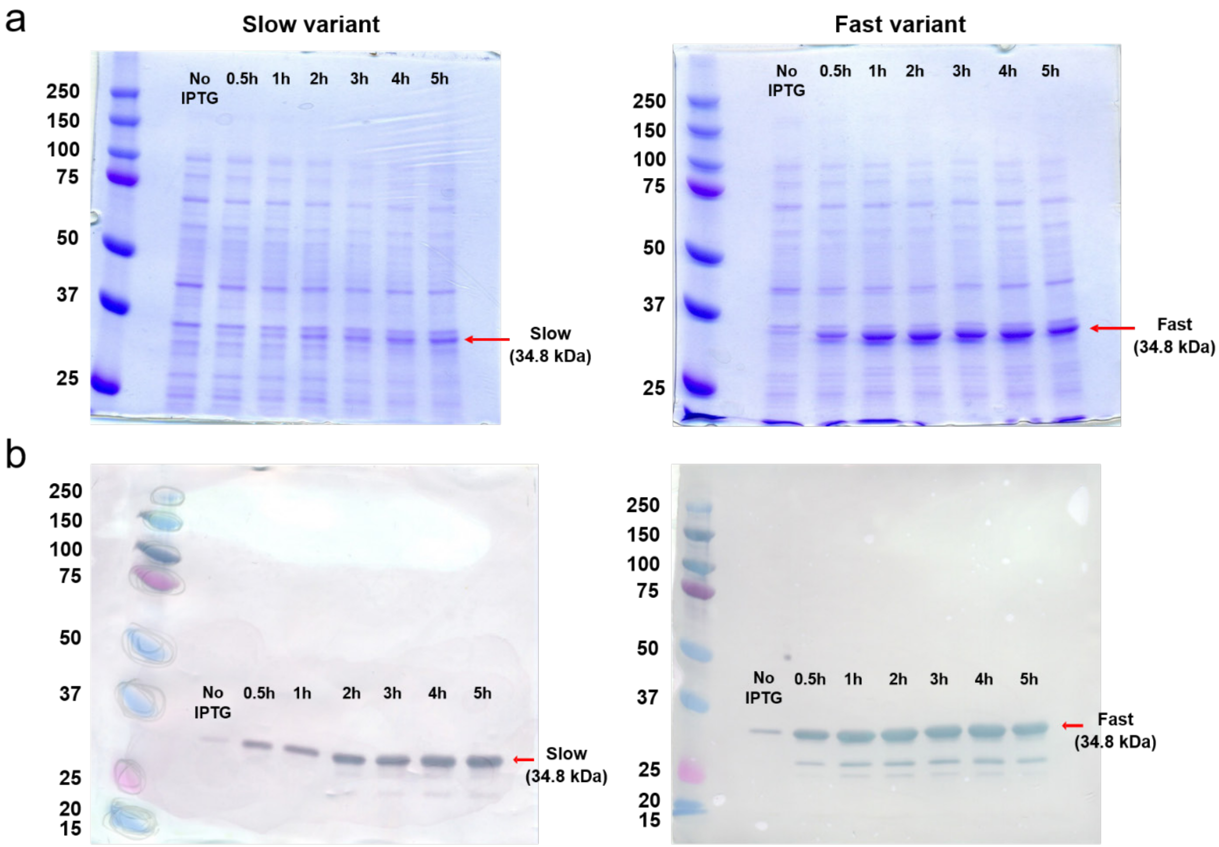

**Supplementary Figure 11. Time-course expression analysis of slow and fast DDLB variants by (a) SDS-PAGE and (b) Western blot ($His_6$-tag), respectively.** Both the gel band intensity in panel a and the $His_6$ band intensity in panel b are higher for the fast variant compared to the slow variant. The fast variant had a higher level of protein expression 5 hours after induction than the slow variant, consistent with the fast mRNA variant translating faster

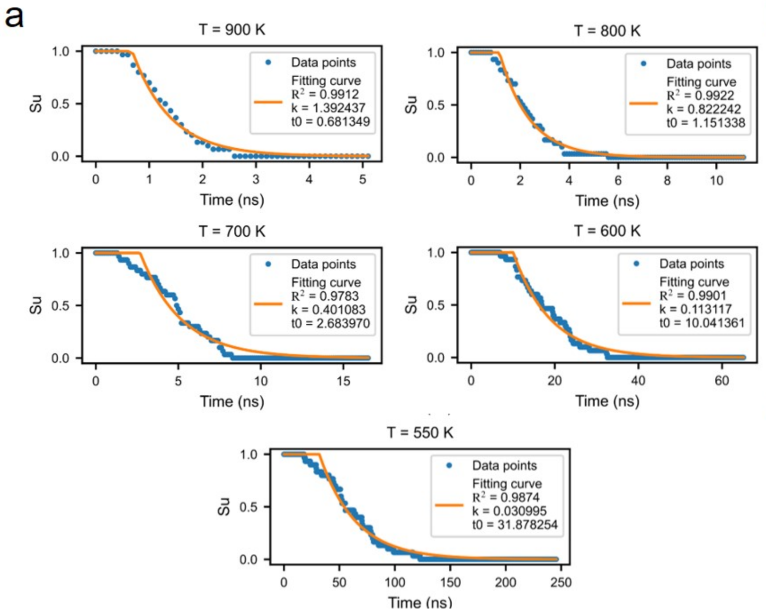

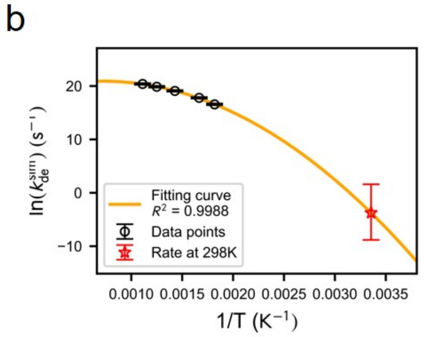

c

| | |
|---|---|
| $k_{\mathrm{de}}^{\mathrm{sim}}$ | 0.023 $s^{-1}$, [1.48 × $10^{-4}$, 4.92] $s^{-1}$ |
| $\frac{k_{\mathrm{uf}}^{\mathrm{sim}}}{k_{\mathrm{uf}}^{\mathrm{exp}}}$ | 143.89 |
| $\tau_{\mathrm{de}}^{\mathrm{rescale}}$ | 6.15 × $10^{3}$ s, [29.24, 9.75 × $10^{5}$] s |

**Supplementary Figure 12. Estimation of the disentangling timescale for CAT-III state P13**. a. Survival probabilities of entangled structures vs. time at different simulation temperatures that were fitted by an exponential function (orange). Fitted parameters (rate *k* and lag time $t_0$) are presented in the legends. b. Arrhenius plots for the disentangling rates that were fitted by a quadratic function. The rates at 298 K were extrapolated. Error bars are presented as 95% CIs estimated by bootstrapping. The rates and timescale at 298 K are presented in panel c with 95% CIs. $k_{\mathrm{uf}}^{\mathrm{sim}}/k_{\mathrm{uf}}^{\mathrm{exp}}$ is estimated from DHFR.

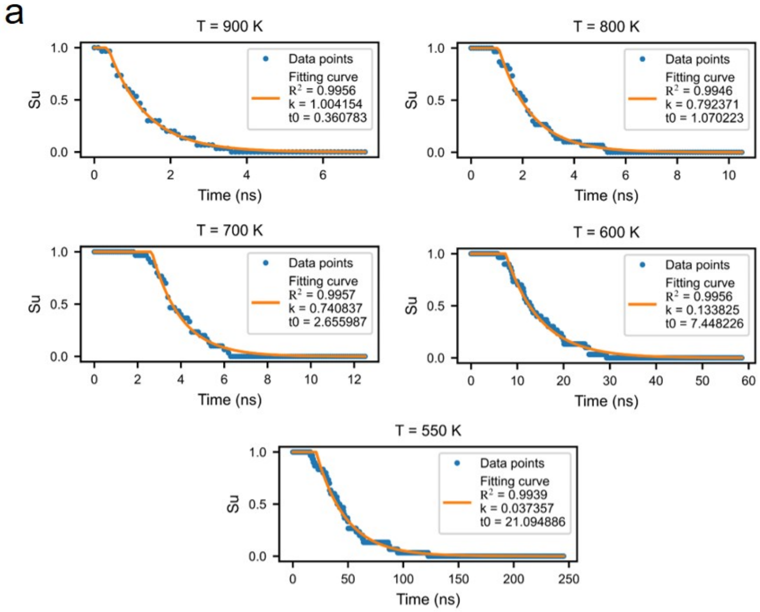

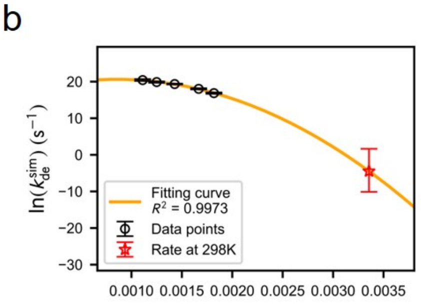

c

| | |
|---|---|
| $k_{\mathrm{de}}^{\mathrm{sim}}$ | 0.011 s$^{-1}$, [4.05 × 10$^{-5}$, 5.17] s$^{-1}$ |
| $\frac{k_{\mathrm{uf}}^{\mathrm{sim}}}{k_{\mathrm{uf}}^{\mathrm{exp}}}$ | 143.89 |
| $\tau_{\mathrm{de}}^{\mathrm{rescale}}$ | 1.26 × 10$^{4}$ s, [27.82, 3.56 × 10$^{6}$] s |

**Supplementary Figure 13. Estimation of the disentangling timescale for DDLB state P4**. a. Survival probabilities of entangled structures vs. time at different simulation temperatures that were fitted by an exponential function (orange). Fitted parameters (rate *k* and lag time $t_0$) are presented in the legends. b. Arrhenius plots for the disentangling rates that were fitted by a quadratic function. The rates at 298 K were extrapolated. Error bars are presented as 95% CIs estimated by bootstrapping. The rates and timescale at 298 K are presented in panel c with 95% CIs. $k_{\mathrm{uf}}^{\mathrm{sim}}/k_{\mathrm{uf}}^{\mathrm{exp}}$ is estimated from DHFR.

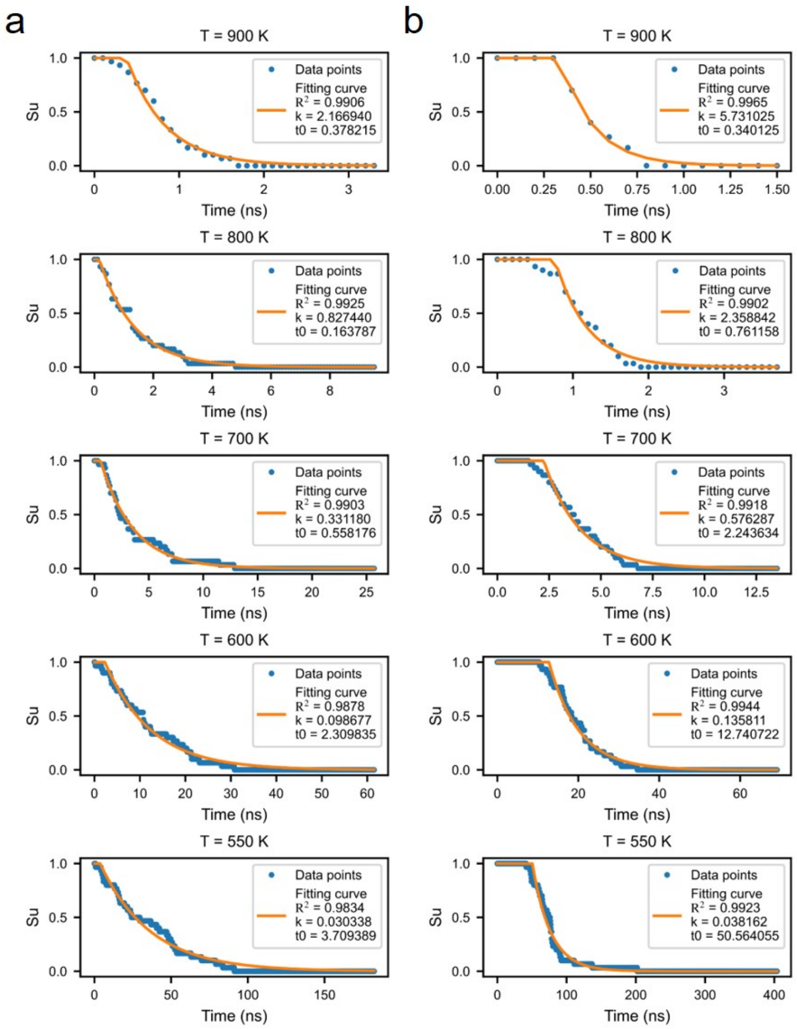

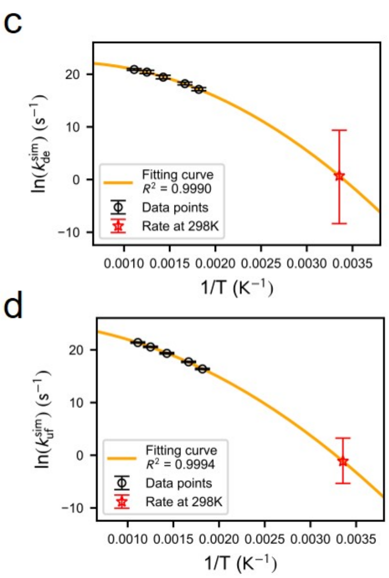

e

| | |
|---|---|
| $k_{\mathrm{de}}^{\mathrm{sim}}$ | 2.03 $s^{-1}$, [2.32 × $10^{-4}$, 1.16 × $10^{4}$] $s^{-1}$ |
| $k_{\mathrm{uf}}^{\mathrm{sim}}$ | 0.32 $s^{-1}$, [4.76 × $10^{-3}$, 25.72] $s^{-1}$ |
| $k_{\mathrm{uf}}^{\mathrm{exp}}$ | 2.24 × $10^{-3}$ $s^{-1}$ |
| $\frac{k_{\mathrm{uf}}^{\mathrm{sim}}}{k_{\mathrm{uf}}^{\mathrm{exp}}}$ | 143.89 |
| $\tau_{\mathrm{de}}^{\mathrm{rescale}}$ | 70.95 s, [4.44 × $10^{-3}$, 1.82 × $10^{6}$] s |

**Supplementary Figure 14. Estimation of the disentangling timescale for DHFR state P2**. Survival probabilities of entangled structures (a) and folded structures (b) vs. time at different simulation temperatures were fitted by an exponential function (orange). Fitted parameters (rate *k* and lag time $t_0$) are presented in the legends. Arrhenius plots for the disentangling rates (c) and unfolding rates (d) were fitted by a quadratic function. The rates at 298 K were extrapolated. Error bars are presented as 95% CIs estimated by bootstrapping. The rates and timescale at 298 K are presented in panel e with 95% CIs.

**Supplementary Videos**

**Supplementary Video I.** Co- and post-translational folding trajectory of a misfolded CAT-III that has a noncovalent lasso entanglement formed after synthesis. The conformation gets trapped in state P13 at the end. The nascent chain is shown in cyan, where the closed loop formed by the native contact is highlighted in red and the segment threading the red loop is highlighted in blue. At each nascent chain length, the conformation is obtained from the last frame of the simulation trajectory. The time of ejection, dissociation and post-translation processes are presented using the experimental timescale on the top left of the scene.

**Supplementary Video II.** Co- and post-translational folding trajectory of a correctly folded CAT-III without any topological entanglements formed. The nascent chain is colored based on secondary structure elements, where alpha-helices are magenta and beta-sheets are yellow. At each nascent chain length, the conformation is obtained from the last frame of the simulation trajectory. The time of ejection, dissociation and post-translation processes are presented using the experimental timescale on the top left of the scene.

**Supplementary Video III.** Co- and post-translational folding trajectory of a misfolded DDLB that has a noncovalent lasso entanglement formed during the synthesis and persisted in the post-translational dynamics. The conformation gets trapped in state P4 at the end. The nascent chain is shown in cyan, where the closed loop formed by the native contact is highlighted in red and the segment threading the red loop is highlighted in blue. At each nascent chain length, the conformation is obtained from the last frame of the simulation trajectory. The time of ejection, dissociation and post-translation processes are presented using the experimental timescale on the top left of the scene.

**Supplementary Video IV.** Co- and post-translational folding trajectory of a correctly folded DDLB without any topological entanglements formed. The nascent chain is colored based on secondary structure elements, where alpha-helices are magenta and beta-sheets are yellow. At each nascent chain length, the conformation is obtained from the last frame of the simulation trajectory. The time of ejection, dissociation and post-translation processes are presented using the experimental timescale on the top left of the scene.

### References

1. O'Brien EP, Christodoulou J, Vendruscolo M, Dobson CM. Trigger factor slows co-translational folding through kinetic trapping while sterically protecting the nascent chain from aberrant cytosolic interactions. *J Am Chem Soc* 2012, **134**(26)**:** 10920-10932.

2. Sharma AK, Bukau B, O'Brien EP. Physical origins of codon positions that strongly influence cotranslational folding: A framework for controlling nascent-protein folding. *J Am Chem Soc* 2016, **138**(4)**:** 1180-1195.

3. Fritch B, Kosolapov A, Hudson P, Nissley DA, Woodcock HL, Deutsch C*, et al.* Origins of the mechanochemical coupling of peptide bond formation to protein synthesis. *J Am Chem Soc* 2018, **140**(15)**:** 5077-5087.

4. Nissley DA, O'Brien EP. Structural Origins of FRET-Observed Nascent Chain Compaction on the Ribosome. *J Phys Chem B* 2018, **122**(43)**:** 9927-9937.

5. Leininger SE, Trovato F, Nissley DA, O'Brien EP. Domain topology, stability, and translation speed determine mechanical force generation on the ribosome. *Proc Natl Acad Sci* 2019, **116**(12)**:** 5523-5532.

6. Best RB, Chen Y-G, Hummer G. Slow protein conformational dynamics from multiple experimental structures: the helix/sheet transition of arc repressor. *Structure* 2005, **13**(12)**:** 1755-1763.

7. Karanicolas J, Brooks III CL. The origins of asymmetry in the folding transition states of protein L and protein G. *Protein Sci* 2002, **11**(10)**:** 2351-2361.

8. Betancourt MR, Thirumalai D. Pair potentials for protein folding: choice of reference states and sensitivity of predicted native states to variations in the interaction schemes. *Protein Sci* 1999, **8**(2)**:** 361-369.

9. Frishman D, Argos P. Knowledge-based protein secondary structure assignment. *Proteins: Struct Funct Bioinform* 1995, **23**(4)**:** 566-579.

10. Eastman P, Swails J, Chodera JD, McGibbon RT, Zhao Y, Beauchamp KA*, et al.* OpenMM 7: Rapid development of high performance algorithms for molecular dynamics. *PLoS Comp Biol* 2017, **13**(7)**:** e1005659.

11. De Sancho D, Doshi U, Munoz V. Protein folding rates and stability: how much is there beyond size? *J Am Chem Soc* 2009, **131**(6)**:** 2074-2075.

12. Reid KL, Rodriguez HM, Hillier BJ, Gregoret LM. Stability and folding properties of a model β-sheet protein, Escherichia coli CspA. *Protein Sci* 1998, **7**(2)**:** 470-479.

13. Nicholson EM, Scholtz JM. Conformational stability of the Escherichia coli HPr protein: test of the linear extrapolation method and a thermodynamic characterization of cold denaturation. *Biochemistry* 1996, **35**(35)**:** 11369-11378.

14. López-Hernéndez E, Serrano L. Structure of the transition state for folding of the 129 aa protein CheY resembles that of a smaller protein, CI-2. *Fold Des* 1996, **1**(1)**:** 43-55.

15. Sugita Y, Okamoto Y. Replica-exchange molecular dynamics method for protein folding. *Chem Phys Lett* 1999, **314**(1-2)**:** 141-151.

16. Kumar S, Rosenberg JM, Bouzida D, Swendsen RH, Kollman PA. The Weighted Histogram Analysis Method for Free-Energy Calculations on Biomolecules. I. The Method. *J Comput Chem* 1992, **13**(8)**:** 1011-1021.

17. Dawson NL, Lewis TE, Das S, Lees JG, Lee D, Ashford P*, et al.* CATH: an expanded resource to predict protein function through structure and sequence. *Nucleic Acids Res* 2016, **45**(D1)**:** D289-D295.

18. Dunkle JA, Wang L, Feldman MB, Pulk A, Chen VB, Kapral GJ*, et al.* Structures of the bacterial ribosome in classical and hybrid states of tRNA binding. *Science* 2011, **332**(6032)**:** 981-984.

19. Arenz S, Bock LV, Graf M, Innis CA, Beckmann R, Grubmüller H*, et al.* A combined cryo-EM and molecular dynamics approach reveals the mechanism of ErmBL-mediated translation arrest. *Nat Commun* 2016, **7:** 12026.

20. Berka K, Hanák O, Sehnal D, Banáš P, Navratilova V, Jaiswal D*, et al.* MOLE online 2.0: interactive web-based analysis of biomacromolecular channels. *Nucleic Acids Res* 2012, **40**(W1)**:** W222-W227.

21. Pravda L, Sehnal D, Toušek D, Navrátilová V, Bazgier V, Berka K*, et al.* MOLEonline: a web-based tool for analyzing channels, tunnels and pores (2018 update). *Nucleic Acids Res* 2018, **46**(W1)**:** W368-W373.

22. Zhang J, Pan X, Yan K, Sun S, Gao N, Sui S-F. Mechanisms of ribosome stalling by SecM at multiple elongation steps. *elife* 2015, **4:** e09684.

23. Bischoff L, Berninghausen O, Beckmann R. Molecular basis for the ribosome functioning as an L-tryptophan sensor. *Cell reports* 2014, **9**(2)**:** 469-475.

24. Su T, Cheng J, Sohmen D, Hedman R, Berninghausen O, von Heijne G*, et al.* The force-sensing peptide VemP employs extreme compaction and secondary structure formation to induce ribosomal stalling. *Elife* 2017, **6:** e25642.

25. Tian P, Steward A, Kudva R, Su T, Shilling PJ, Nickson AA*, et al.* Folding pathway of an Ig domain is conserved on and off the ribosome. *Proc Natl Acad Sci* 2018, **115**(48)**:** E11284-E11293.

26. Voorhees RM, Weixlbaumer A, Loakes D, Kelley AC, Ramakrishnan V. Insights into substrate stabilization from snapshots of the peptidyl transferase center of the intact 70S ribosome. *Nat Struct Mol Biol* 2009, **16**(5)**:** 528.

27. Huter P, Arenz S, Bock LV, Graf M, Frister JO, Heuer A*, et al.* Structural basis for polyproline-mediated ribosome stalling and rescue by the translation elongation factor EF-P. *Mol Cell* 2017, **68**(3)**:** 515-527. e516.

28. Sharma AK, Sormanni P, Ahmed N, Ciryam P, Friedrich UA, Kramer G*, et al.* A chemical kinetic basis for measuring translation initiation and elongation rates from ribosome profiling data. *PLoS Comp Biol* 2019, **15**(5)**:** e1007070.

29. Pai A, You L. Optimal tuning of bacterial sensing potential. *Mol Syst Biol* 2009, **5**(1)**:** 286.

30. Fluitt A, Pienaar E, Viljoen H. Ribosome kinetics and aa-tRNA competition determine rate and fidelity of peptide synthesis. *Comput Biol Chem* 2007, **31**(5-6)**:** 335-346.

31. Athey J, Alexaki A, Osipova E, Rostovtsev A, Santana-Quintero LV, Katneni U*, et al.* A new and updated resource for codon usage tables. *BMC Bioinformatics* 2017, **18**(1)**:** 391.

32. Karpinets TV, Greenwood DJ, Sams CE, Ammons JT. RNA: protein ratio of the unicellular organism as a characteristic of phosphorous and nitrogen stoichiometry and of the cellular requirement of ribosomes for protein synthesis. *BMC Biol* 2006, **4**(1)**:** 30.

33. Nagano N. EzCatDB: the enzyme catalytic-mechanism database. *Nucleic Acids Res* 2005, **33**(suppl_1)**:** D407-D412.

34. Nagano N, Nakayama N, Ikeda K, Fukuie M, Yokota K, Doi T*, et al.* EzCatDB: the enzyme reaction database, 2015 update. *Nucleic Acids Res* 2014, **43**(D1)**:** D453-D458.

35. Consortium U. UniProt: the universal protein knowledgebase. *Nucleic Acids Res* 2018, **46**(5)**:** 2699.

36. Berman HM, Westbrook J, Feng Z, Gilliland G, Bhat TN, Weissig H*, et al.* The protein data bank. *Nucleic Acids Res* 2000, **28**(1)**:** 235-242.

37. De Sancho D, Muñoz V. Integrated prediction of protein folding and unfolding rates from only size and structural class. *Phys Chem Chem Phys* 2011, **13**(38)**:** 17030-17043.

38. Leslie A. Refined crystal structure of type III chloramphenicol acetyltransferase at 1.75 Å resolution. *J Mol Biol* 1990, **213**(1)**:** 167-186.

39. Kauffman LH. *Knots and physics*, vol. 1. World scientific, 2001.

40. Baiesi M, Orlandini E, Seno F, Trovato A. Exploring the correlation between the folding rates of proteins and the entanglement of their native states. *Journal of Physics A: Mathematical and Theoretical* 2017, **50**(50)**:** 504001.

41. Steinhaus H. Sur la division des corps materiels en parties. Bull. Acad. Polon. Sci., C1. III vol IV: 801-804. 1956.

42. MacQueen J. Some methods for classification and analysis of multivariate observations. Proceedings of the fifth Berkeley symposium on mathematical statistics and probability; 1967: Oakland, CA, USA; 1967. p. 281-297.

43. Röblitz S, Weber M. Fuzzy spectral clustering by PCCA+: application to Markov state models and data classification. *Advances in Data Analysis and Classification* 2013, **7**(2)**:** 147-179.

44. Buchete N-V, Hummer G. Coarse master equations for peptide folding dynamics. *J Phys Chem B* 2008, **112**(19)**:** 6057-6069.

45. Scherer MK, Trendelkamp-Schroer B, Paul F, Pérez-Hernández G, Hoffmann M, Plattner N*, et al.* PyEMMA 2: A software package for estimation, validation, and analysis of Markov models. *J Chem Theory Comput* 2015, **11**(11)**:** 5525-5542.

46. O'Brien EP, Ziv G, Haran G, Brooks BR, Thirumalai D. Effects of denaturants and osmolytes on proteins are accurately predicted by the molecular transfer model. *Proc Natl Acad Sci* 2008, **105**(36)**:** 13403-13408.

47. Moore BL, Kelley LA, Barber J, Murray JW, MacDonald JT. High–quality protein backbone reconstruction from alpha carbons using Gaussian mixture models. *J Comput Chem* 2013, **34**(22)**:** 1881-1889.

48. Rotkiewicz P, Skolnick J. Fast procedure for reconstruction of full-atom protein models from reduced representations. *J Comput Chem* 2008, **29**(9)**:** 1460-1465.

49. Tsolis AC, Papandreou NC, Iconomidou VA, Hamodrakas SJ. A consensus method for the prediction of 'aggregation-prone'peptides in globular proteins. *PLoS One* 2013, **8**(1)**:** e54175.

50. Gutierres M, Bonorino C, Rigo M. ChaperISM: improved chaperone binding prediction using position-independent scoring matrices. *Bioinformatics* 2020, **36**(3)**:** 735-741.

51. Schneidman-Duhovny D, Inbar Y, Nussinov R, Wolfson HJ. Geometry-based flexible and symmetric protein docking. *Proteins: Struct Funct Bioinform* 2005, **60**(2)**:** 224-231.

52. Trott O, Olson AJ. AutoDock Vina: improving the speed and accuracy of docking with a new scoring function, efficient optimization, and multithreading. *J Comput Chem* 2010, **31**(2)**:** 455-461.

53. Grossfield A. WHAM: the weighted histogram analysis method. 2.0.10 ed.

54. Darden T, York D, Pedersen L. Particle Mesh Ewald: An N⋅ log (N) Method for Ewald Sums in Large Systems. *J Chem Phys* 1993, **98**(12)**:** 10089-10092.

55. Ryckaert J-P, Ciccotti G, Berendsen HJ. Numerical Integration of the Cartesian Equations of Motion of a System with Constraints: Molecular Dynamics of *n*-alkanes. *J Comput Phys* 1977, **23**(3)**:** 327-341.

56. Case DA, Betz RM, Cerutti DS, Cheatham III TE, Darden TA, Duke RE*, et al.* AMBER. 17 ed. San Francisco: University of California; 2017.

57. Maier JA, Martinez C, Kasavajhala K, Wickstrom L, Hauser KE, Simmerling C. ff14SB: improving the accuracy of protein side chain and backbone parameters from ff99SB. *J Chem Theory Comput* 2015, **11**(8)**:** 3696-3713.

58. Wang J, Wolf RM, Caldwell JW, Kollman PA, Case DA. Development and Testing of a General AMBER Force Field. *J Comput Chem* 2004, **25**(9)**:** 1157-1174.

59. Bayly CI, Cieplak P, Cornell W, Kollman PA. A Well-Behaved Electrostatic Potential Based Method Using Charge Restraints for Deriving Atomic Charges: the RESP Model. *J Phys Chem* 1993, **97**(40)**:** 10269-10280.

60. Frisch M, Trucks G, Schlegel H, Scuseria G, Robb M, Cheeseman J*, et al.* Gaussian 09 Revision D. 01, 2009. *Gaussian Inc Wallingford CT* 2009.

61. Gaus M, Cui Q, Elstner M. DFTB3: extension of the self-consistent-charge density-functional tight-binding method (SCC-DFTB). *J Chem Theory Comput* 2011, **7**(4)**:** 931-948.

62. Gaus M, Goez A, Elstner M. Parametrization and benchmark of DFTB3 for organic molecules. *J Chem Theory Comput* 2012, **9**(1)**:** 338-354.

63. Gaus M, Lu X, Elstner M, Cui Q. Parameterization of DFTB3/3OB for sulfur and phosphorus for chemical and biological applications. *J Chem Theory Comput* 2014, **10**(4)**:** 1518-1537.

64. Kubillus M, Kubar T, Gaus M, Rezac J, Elstner M. Parameterization of the DFTB3 method for Br, Ca, Cl, F, I, K, and Na in organic and biological systems. *J Chem Theory Comput* 2014, **11**(1)**:** 332-342.

65. Lu X, Gaus M, Elstner M, Cui Q. Parametrization of DFTB3/3OB for magnesium and zinc for chemical and biological applications. *J Phys Chem B* 2014, **119**(3)**:** 1062-1082.

66. Leslie A, Moody P, Shaw WV. Structure of chloramphenicol acetyltransferase at 1.75-A resolution. *Proc Natl Acad Sci* 1988, **85**(12)**:** 4133-4137.

67. Jiang Y, Zhang H, Tan T. Rational Design of Methodology-Independent Metal Parameters Using a Nonbonded Dummy Model. *J Chem Theory Comput* 2016, **12**(7)**:** 3250-3260.

68. Shi Y, Walsh CT. Active-site mapping of Escherichia coli D-Ala-D-Ala ligase by structure-based mutagenesis. *Biochemistry* 1995, **34**(9)**:** 2768-2776.

69. Liu CT, Francis K, Layfield JP, Huang X, Hammes-Schiffer S, Kohen A*, et al.* Escherichia coli dihydrofolate reductase catalyzed proton and hydride transfers: Temporal order and the roles of Asp27 and Tyr100. *Proc Natl Acad Sci* 2014, **111**(51)**:** 18231-18236.

70. Lin J. Divergence measures based on the Shannon entropy. *IEEE Trans Inf Theory* 1991, **37**(1)**:** 145-151.

71. Ellsworth BA, Tom NJ, Bartlett PA. Synthesis and evaluation of inhibitors of bacterial D-alanine: D-alanine ligases. *Chem Biol* 1996, **3**(1)**:** 37-44.

72. Zawadzke LE, Bugg TD, Walsh CT. Existence of two D-alanine: D-alanine ligases in Escherichia coli: cloning and sequencing of the ddlA gene and purification and characterization of the DdlA and DdlB enzymes. *Biochemistry* 1991, **30**(6)**:** 1673-1682.

73. Daub E, Zawadzke LE, Botstein D, Walsh CT. Isolation, cloning, and sequencing of the Salmonella typhimurium ddlA gene with purification and characterization of its product, D-alanine: D-alanine ligase (ADP forming). *Biochemistry* 1988, **27**(10)**:** 3701-3708.

74. Lee J, Yennawar NH, Gam J, Benkovic SJ. Kinetic and structural characterization of dihydrofolate reductase from Streptococcus pneumoniae. *Biochemistry* 2010, **49**(1)**:** 195-206.

75. Srinivasan B, Rodrigues JoV, Tonddast-Navaei S, Shakhnovich E, Skolnick J. Rational design of novel allosteric dihydrofolate reductase inhibitors showing antibacterial effects on drug-resistant Escherichia coli escape variants. *ACS Chem Biol* 2017, **12**(7)**:** 1848-1857.

76. To P, Xia Y, Devlin T, Fleming KG, Fried SD. A Proteome-Wide Map of Chaperone-Assisted Protein Refolding in the Cytosol. *bioRxiv* 2022**:** 2021.2011.2020.469408.

77. Neidhardt FC, Bloch PL, Smith DF. Culture medium for enterobacteria. *J Bacteriol* 1974, **119**(3)**:** 736-747.

78. Feng Y, De Franceschi G, Kahraman A, Soste M, Melnik A, Boersema PJ*, et al.* Global analysis of protein structural changes in complex proteomes. *Nat Biotechnol* 2014, **32**(10)**:** 1036-1044.

79. Kong AT, Leprevost FV, Avtonomov DM, Mellacheruvu D, Nesvizhskii AI. MSFragger: ultrafast and comprehensive peptide identification in mass spectrometry–based proteomics. *Nat Methods* 2017, **14**(5)**:** 513-520.

80. To P, Whitehead B, Tarbox HE, Fried SD. Nonrefoldability is pervasive across the E. coli proteome. *J Am Chem Soc* 2021, **143**(30)**:** 11435-11448.

81. Wittung-Stafshede P, Gray HB, Winkler JR. Rapid formation of a four-helix bundle. Cytochrome b 562 folding triggered by electron transfer. *J Am Chem Soc* 1997, **119**(40)**:** 9562-9563.

82. Van Nuland NA, Meijberg W, Warner J, Forge V, Scheek RM, Robillard GT*, et al.* Slow cooperative folding of a small globular protein HPr. *Biochemistry* 1998, **37**(2)**:** 622-637.

83. Raschke TM, Marqusee S. The kinetic folding intermediate of ribonuclease H resembles the acid molten globule and partially unfolded molecules detected under native conditions. *Nat Struct Biol* 1997, **4**(4)**:** 298.
